# Functional genomics of the stable fly, *Stomoxys calcitrans*, reveals mechanisms underlying reproduction, host interactions, and novel targets for pest control

**DOI:** 10.1101/623009

**Authors:** Pia U. Olafson, Serap Aksoy, Geoffrey M. Attardo, Greta Buckmeier, Xiaoting Chen, Craig J. Coates, Megan Davis, Justin Dykema, Scott J. Emrich, Markus Friedrich, Christopher J. Holmes, Panagiotis Ioannidis, Evan N. Jansen, Emily C. Jennings, Daniel Lawson, Ellen O. Martinson, Gareth L. Maslen, Richard P. Meisel, Terence D. Murphy, Dana Nayduch, David R. Nelson, Kennan J. Oyen, Tyler J. Raszick, José M. C. Ribeiro, Hugh M. Robertson, Andrew J. Rosendale, Timothy B. Sackton, Sonja L. Swiger, Sing-Hoi Sze, Aaron M. Tarone, David B. Taylor, Wesley C. Warren, Robert M. Waterhouse, Matthew T. Weirauch, John H. Werren, Richard K. Wilson, Evgeny M. Zdobnov, Joshua B. Benoit

## Abstract

**Background:** The stable fly, *Stomoxys calcitrans*, is a major blood-feeding pest of livestock that has near worldwide distribution, causing an annual cost of over $2 billion for control and product loss in the United States alone. Control of these flies has been limited to increased sanitary management practices and insecticide application for suppressing larval stages. Few genetic and molecular resources are available to help in developing novel methods for controlling stable flies.

**Results:** This study examines stable fly biology by utilizing a combination of high-quality genome sequencing, microbiome analyses, and RNA-seq analyses targeting multiple developmental stages and tissues. In conjunction, manual curation of over 1600 genes was used to examine gene content related to stable fly reproduction, interactions with their host, host-microbe dynamics, and putative routes for control. Most notable was establishment of reproduction-associated genes and identification of expanded vision, chemosensation, immune repertoire, and metabolic detoxification pathway gene families.

**Conclusions:** The combined sequencing, assembly, and curation of the male stable fly genome followed by RNA-seq and downstream analyses provide insights necessary to understand the biology of this important pest. These resources and knowledge will provide the groundwork for expanding the tools available to control stable fly infestations. The close relationship of *Stomoxys* to other blood-feeding (*Glossina*) and non-blood-feeding flies (medflies, *Drosophila*, house flies) will allow for understanding the evolution of blood feeding among Cyclorrhapha flies.

## INTRODUCTION

Livestock ectoparasites are detrimental to cattle industries in the US and worldwide, impacting both confined and rangeland operations. Flies from the Muscidae family commonly occupy these settings, including the nonbiting house fly and face fly and the blood-feeding (hematophagous) stable fly and horn fly. These muscid flies exhibit different larval and adult biologies, varying in larval developmental substrates, as well as adult nutrient sources and feeding frequency [1, 2]. As such, control efforts against these flies are not one size fits all. The stable fly, *Stomoxys calcitrans* (L.), in particular, is a serious hematophagous pest with a cosmopolitan host range, feeding on bovids, equids, cervids, canines, and occasionally humans throughout much of the world. The stable fly’s painful bites disrupt livestock feeding behavior [3-6]; these bites can be numerous during heavy infestation, leading to reductions of productivity by over $2 billion USD [7]. In Australia, Brazil, and Costa Rica, dramatic increases in stable fly populations have coincided with the expansion of agricultural production where the vast accumulation of post-harvest byproducts are recognized as nutrient sources for development of immature stages [8-10].

Stable fly larvae occupy and develop in almost any type of decomposing vegetative materials, e.g. spent hay, grass clippings, residues from commercial plant processing, that are often contaminated with animal wastes [11]. The active microbial communities residing in these developmental substrates, e.g. plant, soil, manure, are required for larval development and likely provide essential nutrients [12]. Even though stable flies are consistently exposed to microbes during feeding and grooming activities, biological transmission (uptake, development, and subsequent transmission of a microbial agent by a vector) of pathogens has not been demonstrated for organisms other than the helminth *Habronema microstoma* [13]. Stable flies have been implicated in mechanical transmission (transfer of pathogens from an infected host or a contaminated substrate to a susceptible host, association between specific vector and pathogen is not necessary) of Equine infectious anemia, African swine fever, West Nile, and Rift Valley Viruses, *Trypanosoma* spp., and *Besnoitia* spp. (reviewed by [13]). The apparent low vector competence of stable flies implicates the importance of immune system pathways not only in regulating larval survival in microbe-rich environments but also in the inability of pathogens to survive and replicate in the adult midgut following ingestion [14-16].

Stable fly mate location and recognition are largely dependent upon visual cues and contact pheromones [17, 18], and gravid females identify suitable oviposition sites through a combination of olfactory and contact chemostimuli along with physical cues [19, 20]. Since stable flies infrequently associate with their hosts, feeding only 1 to 2 times per day, on-animal and pesticide applications are less effective control efforts than those that integrate sanitation practices with fly population suppression by way of traps [21]. Given the importance of chemosensory and vision pathways, repellents have been identified that target stable fly chemosensory inputs and current trap technologies exploit stable fly visual attraction [22-24]. However, despite these efforts, consistent control of stable fly populations remains challenging and development of novel control mechanisms is greatly needed.

Although both sexes feed on sugar, adults are reliant on a bloodmeal for yolk deposition and egg development, as well as seminal fluid production [25, 26]. Blood feeding evolved independently on at least five occasions within the Diptera, in the Culicimorpha, Psychodomorpha, Tabanomorpha, Muscoidea, and Hippoboscoidea [27]. The Muscinae appear to have a high propensity for developing blood feeding; which has occurred at least four times within this subfamily-once in each of the *domestica*-, *sorbens*-and *lusoria*-groups and again in the Stomoxini [28]. Unlike other groups of blood-feeding Diptera where non-blood feeding ancestors are distantly related and / or difficult to discern, stomoxynes are imbedded with the subfamily Muscinae of the Muscidae, featuring many non-blood feeding species. Contrasting blood-feeding culicimorphs and tabanimorphs, stable flies exhibit gonotrophic discordance [29, 30], requiring 3-4 blood meals for females to develop their first clutch of eggs and an additional 2-3 for each subsequent clutch of eggs. These unique aspects of stable flies offer opportunities for comparative analysis of the genomic features underlying these key biological traits.

Even with the importance of the stable fly as a pest, little is known about the molecular mechanisms underlying the biology of *S. calcitrans*. To further our understanding of this critical livestock pest, we report a draft genome sequence of the stable fly. The quality of this genome is high and includes *in silico* annotation that was aided by extensive developmental and tissue-specific RNA-seq data focusing on the feeding and reproduction of *S. calcitrans*. Manual curation and comparative analyses focused on aspects related to host interactions, reproduction, control, and regulation of specific biological processes. Our study significantly advances the understanding of stable fly biology including the identification of unique molecular and physiological processes associated with this blood-feeding fly. These processes can serve as novel targets which will assist in both developing and improving control of this important livestock pest.

## RESULTS AND DISCUSSION

### Genome assembly and annotation supported by comparative and functional genomics

Whole genome shotgun sequencing of adult males resulted in the 66x coverage draft assembly of 971 MB of total sequence. Scaffolds (12,042) and contigs (125,702) had N50 lengths of 504.7 and 11.3 kb, respectively. The sequence was ∼20% smaller compared to the genome predicted by propidium iodide analyses (∼1150 MB, [31]). This difference is likely the result of heterochromatin and other repetitive regions that were unassembled, as genome size is not significantly different between the sexes [31], and is comparable to differences documented for other insect genomes [32-34]. Further details of the stable fly genome assembly and analyses are provided as supplementary information (Additional Files 1 and 2). There were 16,102 predicted genes/pseudogenes that included 2,003 non-protein coding genes, and a total of 22,450 mRNA transcripts were predicted with over 90-95% supported by RNA-seq (Additional File 2:Table S2). Manual curation and analyses allowed preliminary chromosome arm assignment and identification of repeat elements from the genome (Additional File 1, Sections 4 and 5) and included the combined analyses and correction of over 1,600 genes focused on gene families underlying reproduction, immunity, host sensing, feeding, and insecticide resistance.

Completeness of the genome was assessed through identification of sets of benchmarking universal single-copy orthologs (BUSCOs) among flies, and BUSCOs identified were comparable to those in other flies for both the genome and predicted gene set (Additional File 1: Fig. S1). Further, the number of genes with significant alignment to *Drosophila* spp. genes was comparable to other published fly genomes (Fig. 1). Lastly, CEGMA genes and those associated with autophagy were all identified from the *S. calcitrans* genome (Additional File 2, Tables S3 and S4), which can be used as an additional metric of genome completeness as these are highly conserved among flies [32, 35]. These metrics indicate that the genome is of sufficient quality for subsequent comparative analyses with other insects. Comparison of protein orthologs revealed only 47 *Stomoxys* species-specific protein families relative to other higher flies (Fig. 1). Based on gene ontology, there was enrichment for zinc finger transcription factors in relation to all genes, which has also been documented in other insect systems [36, 37].

**Figure 1.**
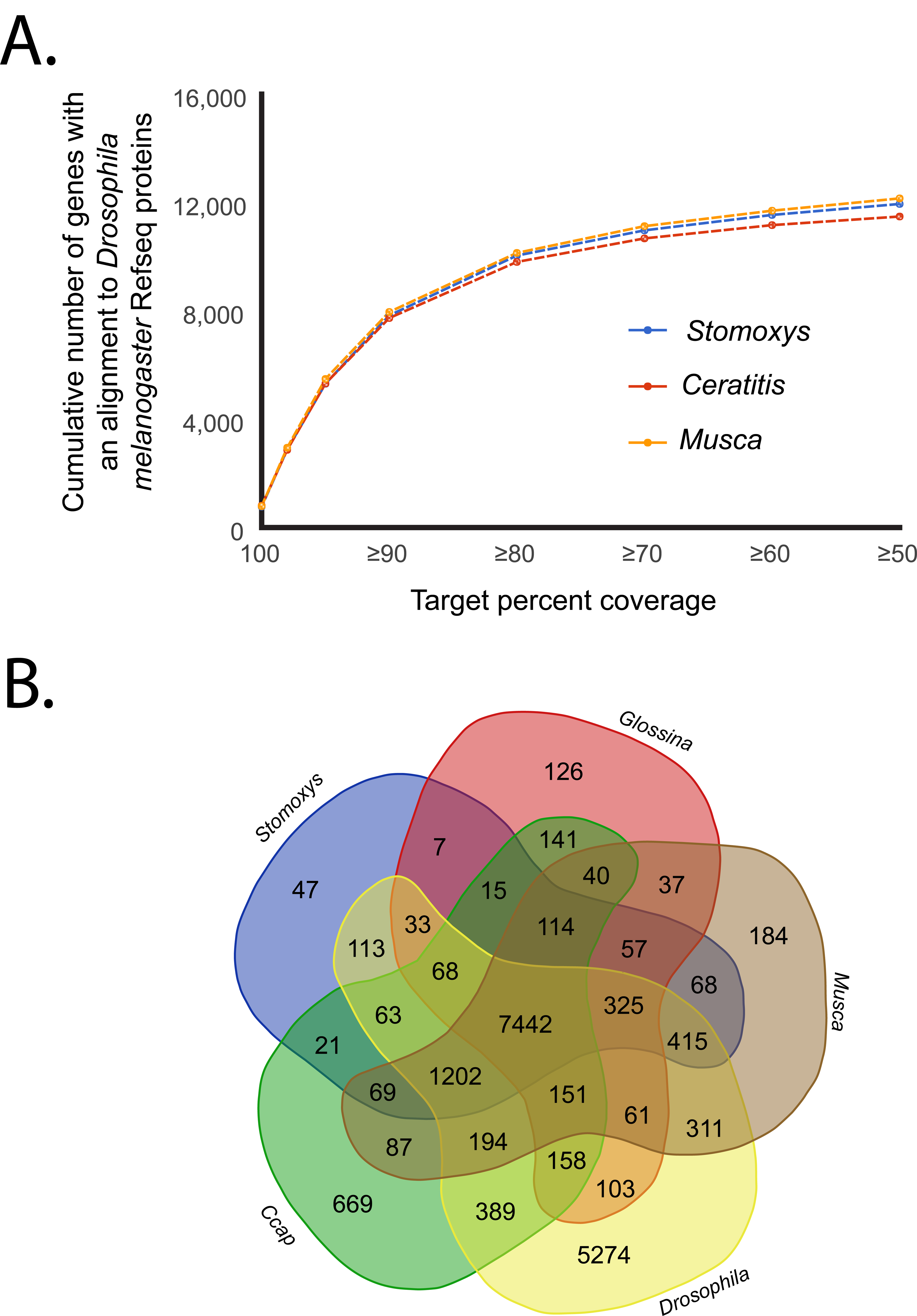
Quality assessment of Stomoxys genome. A. Number of genes with alignment to *Drosophila melanogaster* genome. **B**. Ortholog group comparison between *Stomoxys* and other higher flies based on comparison to the OrthoDB8 database. All *Drosophila* species were merged for the analysis.

RNA sequencing produced a comprehensive catalog of expression profiles for all genes sampling both sexes, as well as different developmental stages, tissues, and specific organs (Additional File 2: Tables S5 – S13). Specifically, RNA collected from whole females (teneral and mated, reproductive), whole males (teneral), male reproductive tracts (mated), female reproductive tracts (mated), male heads (fed, mated), female heads (fed, mated), third instar larva, and pooled female/male salivary glands were examined to assist in addressing core questions of this study. The RNA-seq datasets were validated by quantitative RT-PCR on a sample of 25 genes with a Pearson’s correlation of 0.85 (Additional File 1: Section 3; Additional File 2: Table S14). RNA-seq datasets are discussed below in relation to the targeted areas of focus.

### Unique duplications in the yolk protein gene family and evidence of male-biased genes with putative seminal fluid function

Yolk proteins (YPs) in flies act as a primary source of nutrients for developing embryos. These proteins function as a source of amino acids and also likely function as transporters of other essential nutrients such as lipids and vitamins [38]. While many insects utilize YPs classified as vitellogenins, cyclorrhaphan (higher) flies such as *Stomoxys* [39], *Drosophila* [40], *Musca* [41], *Calliphora* [42], and *Glossina* [43] utilize an alternative class of proteins derived from lipase enzymes termed YPs [44]. The number of yolk protein genes varies among species. The species-specific expansions/contractions observed within this class of genes may reflect reproductive demand within those species. Our analysis identified 8 putative *Stomoxys* YP homologs relative to YP gene family members in *Musca* [36], four of which we identified during our analysis.

To understand the evolutionary relationships between characterized YPs we performed a phylogenetic analysis of the predicted YPs from *S. calcitrans, M. domestica, G. morsitans* and *Drosophila melanogaster* (Fig. 2). Based on this analysis, the yolk protein gene family expanded in *S. calcitrans* and *M. domestica* sometime after their divergence from *Drosophila*. Of the *Musca* and *Stomoxys* specific YPs, 3 members are orthologous between the two species suggesting derivation from a common ancestor. However, the remaining yolk protein genes are paralogous and appear to originate from independent duplication events occurring after the divergence of the *Stomoxys* and *Musca* lineages. The lineage-specific expansions suggest that duplications in this gene family may confer a reproductive advantage by increasing reproductive capacity. In support of this role, all 8 YP genes are upregulated in reproductively active *S. calcitrans* females. Expression is not observed in the female reproductive tract, which suggests these genes are expressed and translated in the fat body, secreted into the hemolymph and transported to the ovaries as observed in other higher flies (Fig. 2).

**Figure 2.**
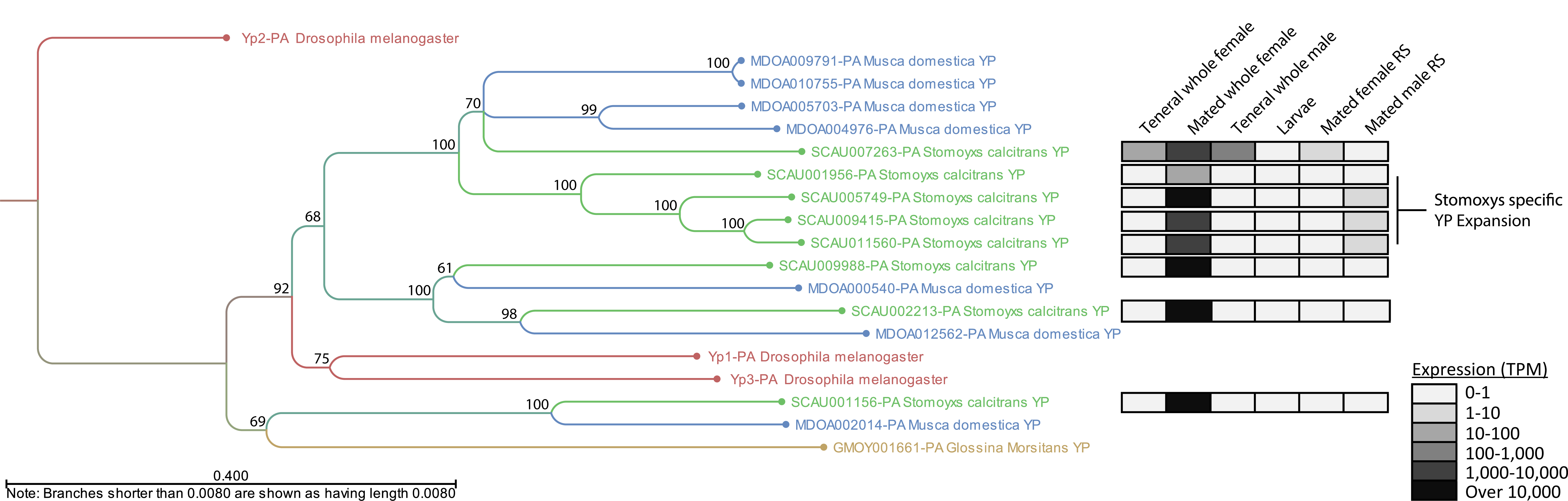
Maximum Likelihood Phylogenetic analysis of yolk protein genes from *Drosophila melanogaster, Glossina morsitans, Musca domestica* and *Stomoxys calcitrans*. Numerical annotations indicate bootstrap values for each branch point in the tree. Heat map of gene expression (transcripts per million, TPM) is based on RNA-seq data (Additional File 2, Tables S5–13). RS, reproductive system.

To identify male-specific reproductive genes and putative seminal proteins we performed an analysis comparing RNA-seq data derived from male and female reproductive tract tissues (Fig. 3). Genes with a male reproductive expression of at least 50 reads per kilobase of transcript per million mapped reads (RPKM) or higher and that were expressed at least 5-fold higher in males relative to females were selected as male biased. This analysis resulted in the classification of 763 genes with male reproductive tract biased expression (Additional File 2: Table S18).

**Figure 3.**
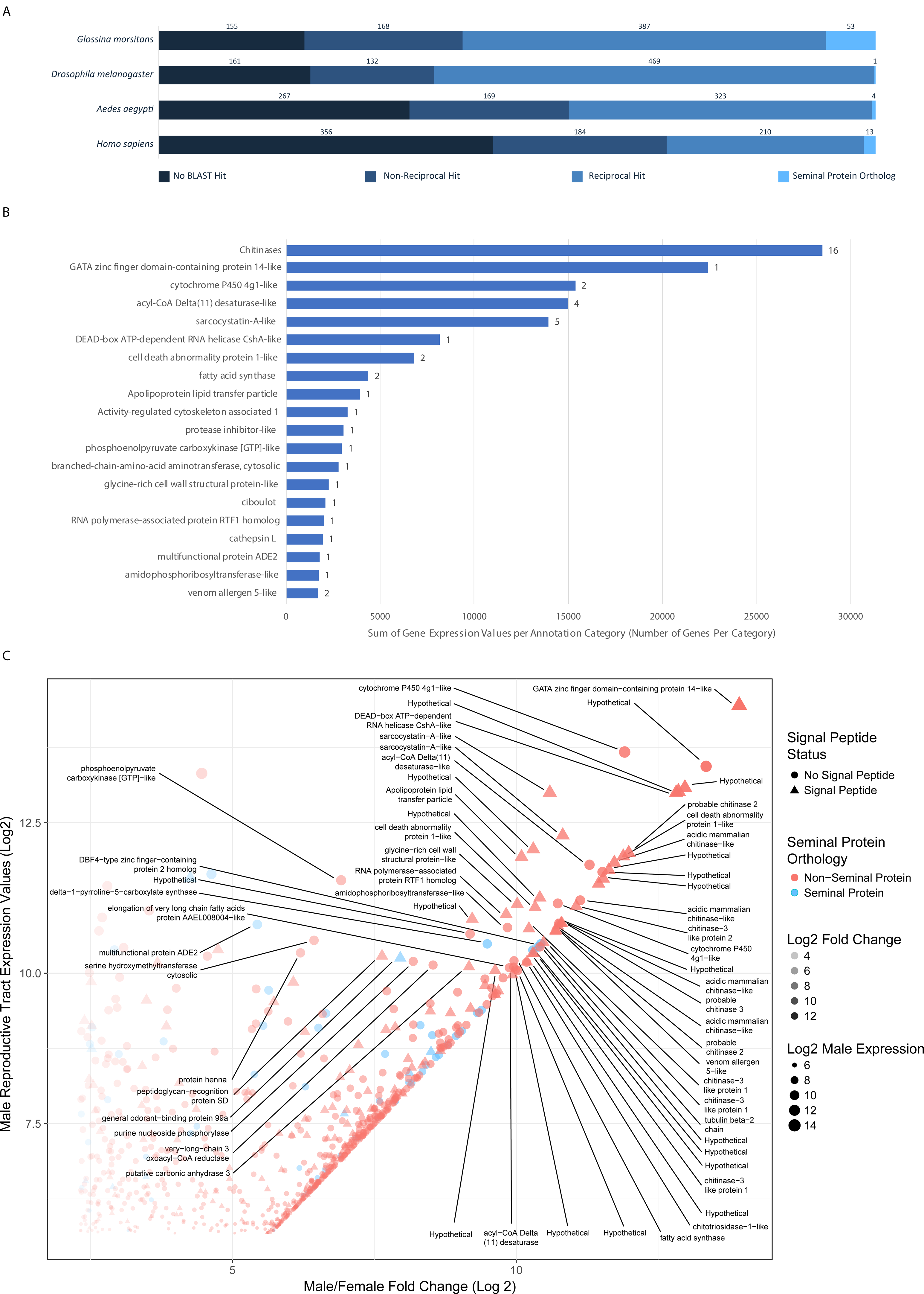
Analysis of male reproductive biased genes in *Stomoxys calcitrans*. A. Results of reciprocal BLAST analysis of male reproductive tract biased genes. **B.** Expression analysis of top 20 most abundant gene classes as annotated by BLAST best hits. Bar length represents combined expression values in RPKM for the genes included in that category. Numbers associated with the bars represent the number of genes in that functional classification. **C.** Scatter plot of the 763 *Stomoxys* male reproductive biased genes. The plot shows on the x axis-log_2_ fold change expression in males relative to females and the y axis represents the log_2_ transformed expression value in RPKM in the male reproductive tissue. Triangular points are genes predicted to contain signal peptides and blue points are genes with orthology to seminal proteins in other species. Genes with a log_2_ expression value above 10 and log_2_ Male/Female Fold Change value above 5 are annotated with putative functional descriptions.

We performed a reciprocal BLAST analysis to identify orthologs of the male biased *Stomoxys* genes in other species in which seminal proteins are characterized including *G. morsitans* [45], *D. melanogaster* [46, 47], *Ae. aegypti* [48] and *Homo sapiens* [49] (Fig. 3). In general, the number of orthologous sequences identified corresponded to the basic phylogenetic relationships between the species tested. However, these relationships did not hold for the number of gene orthologs associated with seminal function. Reciprocal analysis with *Drosophila* identified 469 1:1 orthologs of male biased *Stomoxys* genes, amounting to the largest number of orthologs identified between species included in this analysis. In contrast, of those 469 orthologs only 1 is associated with seminal fluid function in *D. melanogaster*.Comparison with *G. morsitans* identified the second highest number of orthologous proteins (387). Of those, 53 were associated with seminal function, suggesting a greater similarity in the constitution of seminal secretions between *Glossina* and *Stomoxys* consistent with their closer phylogenetic relationship compared to *Drosophila*. Of note, none of the *Stomoxys* male biased proteins were orthologous to seminal proteins across all four species. In *Drosophila*, male-biased genes evolve at a faster rate, especially those expressed in reproductive tissues, and they tend to lack identifiable orthologs compared with genes expressed in an unbiased pattern [50, 51]. There is evidence for this in *Musca* as well [52], and these may be due to selection pressure resulting in rapid evolution of male-biased genes [51]. One gene, however, a catalase (XM_013259723), was orthologous to seminal protein genes in *Aedes, Glossina* and *Homo sapiens* (Fig 3A). As catalases function to reduce oxidative stress, this finding could reflect a conserved mechanism that protects the sperm from oxidative damage.

The 763 male biased *Stomoxys* genes were annotated by BLAST and gene ontology analysis and then categorized by best hit annotation. Of those genes, 216 lacked significant BLAST hits or were homologous to hypothetical proteins with no functional associations. Of the remaining genes that had significant hits to annotated proteins, certain categories were highly expressed in terms of both the number of genes and the relative level of expression within the male reproductive tract (Fig. 3B+C).

The top category for which function could be assigned was comprised of 16 genes with chitinase activity, 12 of which are clustered on scaffolds KQ079939 (7 genes) and KQ080089 (5 genes). Chitinases confer anti-fungal activity in honey bee seminal secretions, preventing the transfer of pathogenic spores during copulation [53]. Such protective properties would be beneficial to *Stomoxys* given the high probability of exposure to fungi in the moist and microbe rich substrates in which females oviposit. The second most highly expressed category consisted of a single gene, XM_013245551, which is the most highly expressed gene in the male reproductive library. While it is annotated as a GATA zinc-finger domain containing protein, further analysis reveals little in the way of conserved domains to indicate its function. The high level of expression of this gene suggests it is an important participant in the functions of this tissue. Cytochrome P450s were an additional class of male biased reproductive tract genes. Cytochrome P450 proteins are involved in a wide array of processes and are associated with molecular modifications, such as hormone biosynthesis and detoxification of xenobiotics [54, 55]. This analysis provides some insight into genetic associations with male reproductive functions in *Stomoxys* and highlights a number of interesting targets for functional analysis in the future. Such analyses could provide key targets for *Stomoxys* control strategies including sterile male production and novel reproductive inhibitors.

### Evidence of muscid-specific odorant binding proteins and odorant receptor lineages

Chemosensory pathways rely on gene families encoding odorant binding proteins, carrier proteins for lipid molecules, as well as odorant, gustatory, and ionotropic receptors that display different affinities for air-borne molecules, mediating the insect’s response to its environment. The *Stomoxys* and *Musca* genomes encode numerous lineage-specific expansions and contain signatures of many pseudogenizations/deletions of chemosensory pathway genes relative to *Drosophila* (Figs. 4 – 6, Additional File 1: Section 7), consistent with the birth-and-death model of evolution proposed for these gene families [56].

**Figure 4.**
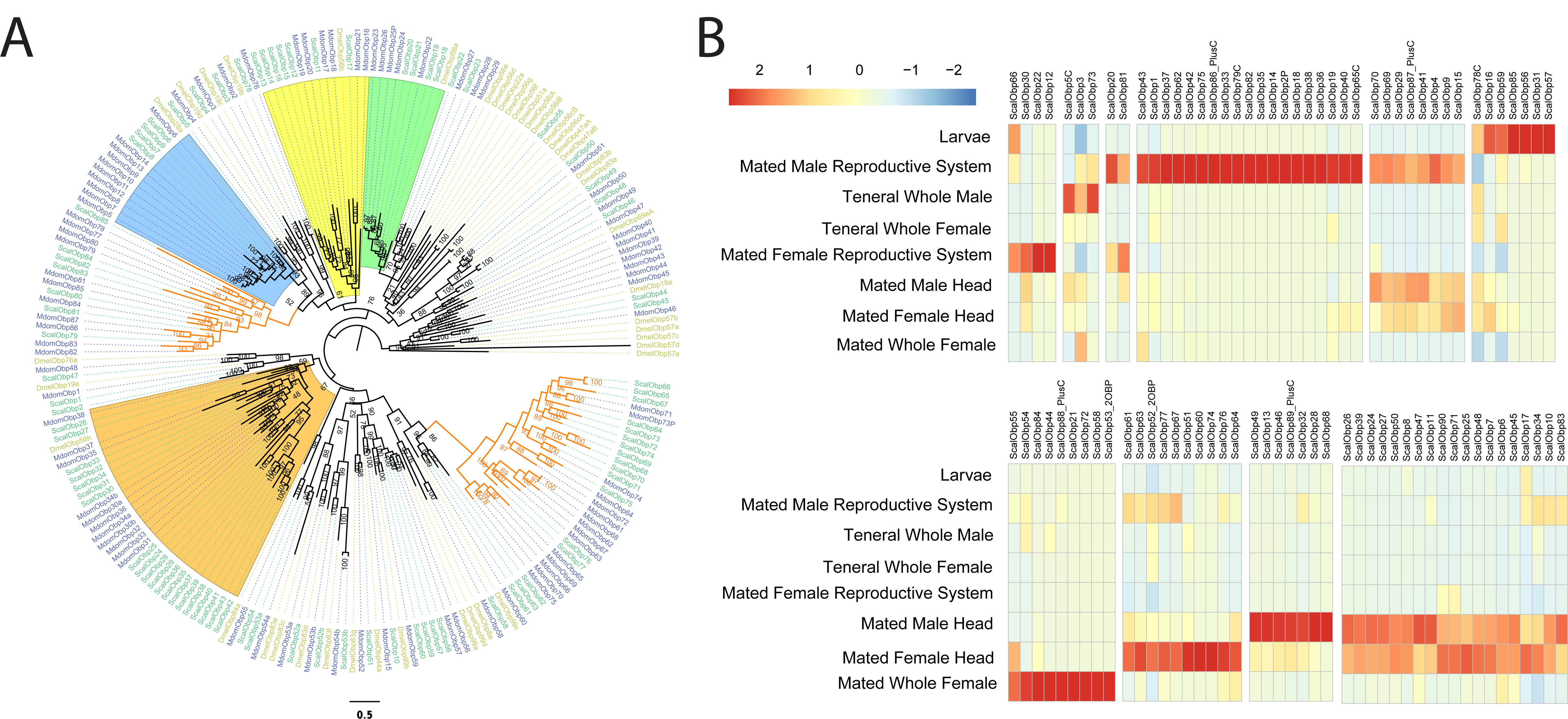
Phylogenetic tree of the *Stomoxys calcitrans* OBPs with those of *Drosophila melanogaster* and *Musca domestica* along with RNA-seq expression. A. Maximum likelihood phylogeny was constructed using the web server version of IQ-TREE software ([177]; best-fit substitution model, branch support assessed with 1000 replicates of UFBoot bootstrap approximation). The *S. calcitrans* and *M. domestica* gene/protein names are highlighted in teal and blue, respectively, while *D. melanogaster* names are in mustard. Clades that are expanded in the muscids relative to *Drosophila* are shaded in orange, blue, yellow, and green. B. OBP transcripts with detectable expression by RNASeq among tissues and developmental stages, based on complete results within Additional File 2, Table S22.

**Figure 5.**
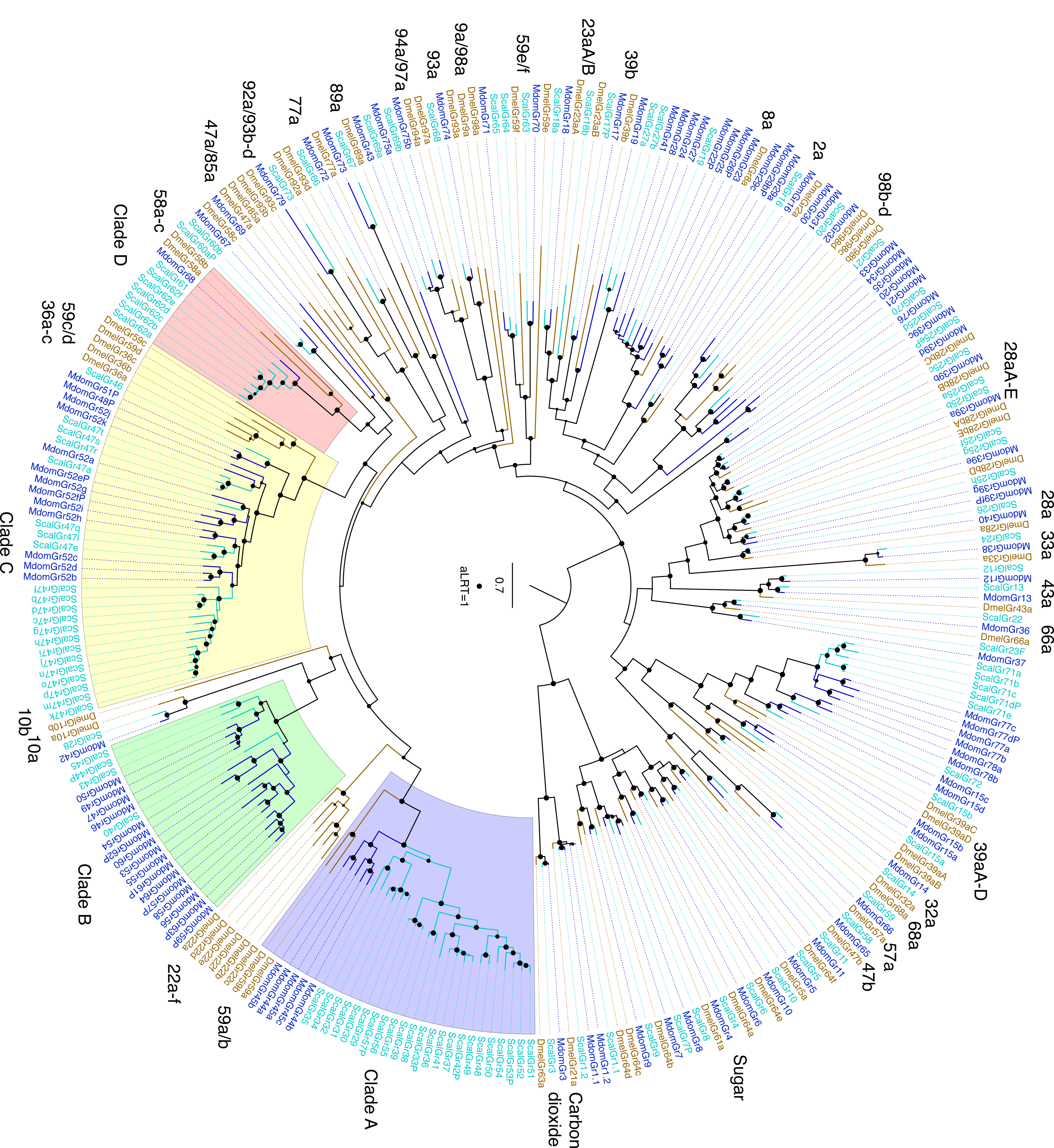
Phylogenetic tree of the *Stomoxys calcitrans* GRs with those of *Drosophila melanogaster* and *Musca domestica.* Maximum likelihood tree rooted by declaring the distantly-related and divergent carbon dioxide and sugar receptor subfamilies as the outgroup. The *S. calcitrans* and *M. domestica* gene/protein names are highlighted in blue and teal, respectively, while *D. melanogaster* names are in mustard. Support levels from the approximate Likelihood-Ratio Test (aLRT) from PhyML v3.0 are shown on branches. Subfamilies and individual or clustered *Drosophila* genes are indicated outside the circle to facilitate finding them in the tree. Four clades of candidate bitter receptors that are expanded in the muscids are highlighted. Pseudogenic sequences are indicated with the suffix P. Scale bar indicates amino acid substitutions per site.

**Figure 6.**
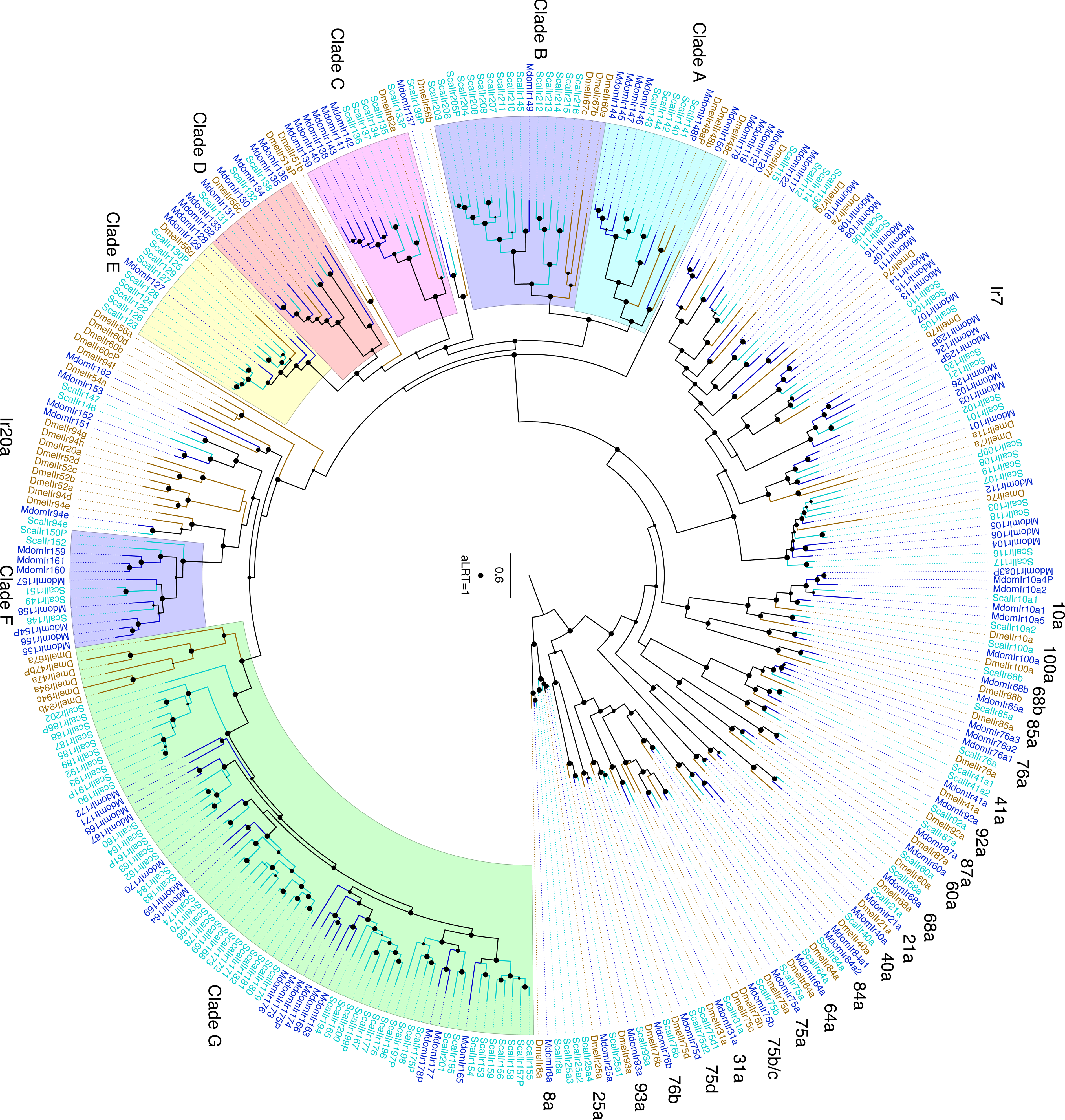
Phylogenetic tree of the *Stomoxys calcitrans* IRs with those of *Drosophila melanogaster* and *Musca domestica.* Maximum likelihood tree rooted by declaring the Ir8a/25a lineage as the outgroup. The *S. calcitrans* and *M. domestica* gene/protein names are highlighted in blue and teal, respectively, while *D. melanogaster* names are in mustard. Support levels from the approximate Likelihood-Ratio Test from PhyML v3.0 are shown on branches. Subfamilies, clades, and individual *Drosophila* genes are indicated outside the circle to facilitate finding them in the tree. Pseudogenic sequences are indicated with the suffix P. Scale bar indicates amino acid substitutions per site.

#### Odorant Binding Protein (OBP) Gene Family

The *Stomoxys* OBP family is comprised of 90 genes, more than half of which are organized as tandem clusters across three scaffolds, consistent with OBP gene organization in other dipteran genomes [57] (Fig. 4, Additional File 2: Table S19). Further, two lineages of OBPs appear unique to *Stomoxys* and *Musca* (Fig. 4A, orange lineage). Expression of 44 *Obp*s was detected in heads of both mated, adult females and males (Fig. 4B), of which 7 *Obps* and 10 *Obps* were highly enriched in heads of mated males and females, respectively. This expression pattern may indicate a role for the genes in mediating chemosensory interactions between the sexes. There is a major expansion of 31 *Stomoxys* (ScalObp11-43) and 23 *Musca* (MdomObp16-38) genes related to the *DmelObp56* gene cluster (*DmelObp56a,b,d,e,h*), with 20 of these *Stomoxys* genes related to *DmelObp56h* (Fig. 4A, orange shade). *DmelObp56h* has a role in male mating behavior, as gene silencing results in distinct changes in the cuticular hydrocarbon profile of male *Drosophila* and in reduction of 5-T, a hydrocarbon that is produced by males and thought to delay onset of courtship [103]. While *DmelObp56h* is expressed exclusively in adult *Drosophila* female and male heads, *Stomoxys* transcripts in this expansion have a diverse expression profile and are detected in not only head tissue of mated males and females, but also in the reproductive tract (RT) tissue of mated males, in larvae, and in mated, whole adult females. Whether these genes have roles in muscid mating behavior is unknown.

Twenty-one *Obps* were detected by RNASeq in reproductive tract (RT) tissues of mated, adult males and females (Fig. 4). This was not unexpected given expression of *Obp*s in non-sensory tissues and deduced roles not related to chemosensation that are reported in other dipteran species [58, 59]. The transfer of OBPs from males to females in seminal fluid occurs in *Drosophila, Glossina*, and *Ae. aegypti* [47, 60, 61], and this may account for *Obp* enrichment in *Stomoxys* male RT. *ScObp 12* and *22* are highly enriched in the RT tissue of mated females, suggesting putative roles for these genes in female reproduction (Fig. 4).

#### Odorant Receptor (OR) Gene Family

The *Stomoxys* OR family is comprised of 74 gene models, including a lineage of four and three ORs that appears unique to *Stomoxys* and *Musca*, respectively (Additional File 1, Fig. S7; Additional File 2: Table S21). Seventeen *Or*s were enriched in the RT of mated males, and these may have a role in sperm activation, as proposed for *An. gambiae* [62]. During stable fly mating, males ‘perch and dart’ towards females after visual and contact pheromone recognition [27]. If sperm transfer is successful, females seldom re-mate [63, 64] suggesting a shift in post-mating behavior. Interestingly, three *Or*s (*ScalOr22, 54*, and *55*) were highly enriched in the RT of mated females relative to all other tissues (Additional File 1, Fig. S7b), suggesting these *Or*s may have a role in female reproduction, such as perceiving male pheromones transferred during copulation.

As in *Musca* and *Glossina* sp., there is an expansion of *Stomoxys* OR genes related to *DmelOr67d* and *DmelOr45a* [33, 52], and smaller subsets of *Stomoxys* ORs duplicated relative to *Drosophila* are present in the *Stomoxys* genome (Additional File 1: Section 7). The ScalOr expansion related to *DmelOr67d* is present with five intact genes (*ScalOr50, ScalOr54-57*) and three pseudogenes (*ScalOr51-53*) organized across six scaffolds; *ScalOr54 and 55* were highly enriched in the RT of mated *Stomoxys* females. In *Drosophila, DmelOr67d* has a role in recognizing a male-specific mating pheromone, cis-vaccenyl acetate [65], that regulates mating behaviors [66]. Seven ScalOr genes (*ScalOr22 – 28*) are related to *DmelOr45a*, which is expressed solely during the larval stage in *Drosophila* and mediates the response to octyl acetate, a repellent substance that induces an escape response in larvae [67]. Expression of the *Scal* orthologs, however, is not restricted to the larval stage, suggesting an expanded role for this receptor in *Stomoxys* adult responses.

Eleven *Ors* were detected in third instar larvae, none of which were highly enriched during this immature lifestage. Further evaluation by non-quantitative RT-PCR detected an additional 9 *Or*s expressed in first and second instar but not in third instar larvae (Additional File 2: Table S21). This suggested that stable flies differentially utilize odorant receptors throughout immature development. Expression of all 20 of these *Ors* was not exclusive to the larval stages, and the absence of larval-specific receptors in the stable fly may be a result of exposure to related compounds during the immature and adult stages, e.g. host dung, detritus. This is in contrast with mosquito species that occupy a larval aquatic habitat distinct from that of the adult. Similarly, both female and male stable flies are blood feeders and sex-biased receptors to enhance host or nutrient localization in one gender over the other may be less critical.

### Gustatory and ionotropic receptor gene family expansions support importance of bitter taste perception

#### Gustatory Receptor (GR) Gene Family

The gustatory receptor family is the more ancient of the two families that make up the insect chemoreceptor superfamily [68-71], and comprises several highly divergent lineages, most involved in taste but some in olfaction [72]. The OR family arose from a GR lineage at the base of the Insecta [73, 74]. While the family is generally divided into three major and divergent subfamilies (sugar or sweet receptors, the carbon dioxide receptors, and the bitter taste receptors), a lineage within the bitter taste receptor clade has evolved into an important receptor for fructose [75], and there are others involved in courtship [76].

The carbon dioxide, sugar, and fructose receptors are relatively well conserved in *Stomoxys* and *Musca*, as is the case for many other insects. However, the bitter taste receptors reveal considerable gene family evolution both with respect to the available relatives of these muscid flies (*Drosophila* and *Ceratitis*) and between these two muscids. For example, a major expansion (*ScalGr29-57*) encoding 49 candidate bitter taste receptors occurs in *Stomoxys*, comparable to a similarly complicated set in *Musca* (*MdomGr43-64*, encoding 35 proteins). Together, these form four major expanded clades in the muscids (Clade A – D, Fig. 5). Insect carbon dioxide receptors are comprised of heterodimerized GRs encoded by highly conserved gene lineages, Gr1 – 3 [77]. *Stomoxys* has the same set of carbon dioxide receptors as *Musca*, Gr1 (*DmelGr21a*) and Gr3 (*DmelGr63a*), with a duplication of the Gr1 lineages present in inverted orientation like *Musca*. The absence of the Gr2 lineage, which is also absent from *Drosophila*, helps confirm that this loss occurred before the Muscidae and Drosophilidae split, but after they separated from the Tephritidae because the Gr2 lineage is present in *Ceratitis* [34].

The expression of 35 *Gr*s was detected in heads of mated females and males, of which 6 and 5 *Grs* were highly enriched in the female and male tissue, respectively. These 11 were candidate bitter taste receptors, predominantly members of the expanded clades A, C, and D (Additional File 1, Fig. S3b; Additional File 2, Table S22). Interestingly, 27 *Gr*s were enriched in the male RT tissues (Additional File 1, Fig. S3b; Additional File 2, Table S20). Evidence from *Drosophila* supports the expression of GRs in neurons that innervate testes and oviducts [78], suggesting that these *Stomoxys* GRs may have a role in mediating reproductive system function. Twenty-three *Grs* were detected in larvae, all of which were candidate bitter taste receptors, suggesting they mediate larval bitter sensing. Determination of the ligand specificities of these muscid receptors is required to fully understand the ecological significance of the differential expansions and contractions of their bitter taste abilities.

Stable flies rely on chemosensory input to mediate localization of nutritional resources, such as volatiles emitted by cattle [19, 20, 79-82] and volatiles/tastants produced from plant products [25, 83, 84]. Stable fly adults likely respond to bacterial communities occupying various substrates when identifying ideal ovipositional sites [12, 19, 82, 85], processing microbial volatiles and assessing substrate suitability for moisture level and temperature.

#### Ionotropic Receptor (IR) Gene Family

The ionotropic receptor family is a variant lineage of the ancient ionotropic glutamate receptor family [68, 86-88]. Like the GRs they are involved in both olfaction and gustation, as well as sensing light, temperature, and humidity [88]. In *Drosophila* and most other insects examined to date, Ir8a and 25a, both of which function as co-receptors with other IRs [88], are highly conserved both in sequence and length and in being phylogenetically most closely related to the ionotropic glutamate receptors from which this variant ionotropic receptor family of chemoreceptors evolved [86, 87]. While *Stomoxys* has the expected single conserved ortholog of Ir8a, surprisingly it has four paralogs of Ir25a (*ScalIr25a1-4*), the functions of which are enigmatic, as such duplications of Ir25a have seldom been observed in other insects (Fig. 6).

The *Ir7a-g* and *11a* genes in *Drosophila* are expressed in larval and adult gustatory organs [87], but ligands for these receptors are unknown. This subfamily is considerably expanded in both *Musca* and *Stomoxys* and, given their complexities, they are not named for their *Drosophila* relatives. Rather, these are part of the numbered series from Ir101, in the case of *Stomoxys* to Ir121 and in *Musca* to Ir126 (Fig. 6). These IR gene family expansions strongly suggest an expanded gustatory capacity.

A large clade of “divergent” IRs in *Drosophila* is involved in gustation and is known as the Ir20a subfamily of 33 proteins [89-91]. This clade of mostly intronless genes is considerably expanded in *Musca* to 53 members (*MdomIr127-179*), and even more so in *Stomoxys* to 96 members (*ScalIr122-217*). This subfamily consists mostly of minor expansions in *Drosophila* and major expansions in the two muscids, labeled clades A-G in the tree (Fig. 6). Of note is clade B, which has just one *Musca* gene (*MdomIr149*), but 15 *Stomoxys* genes (*ScalGr203-217*), apparently related to DmelIr60e and 67b/c. Information on putative IR ligands in *Drosophila* are limited to carbonation sensing and specific carbohydrates [92, 93].

RNA-seq detected 23 *Ir*s in heads of mated females and males of which 2 and 7 were enriched in the female and male tissue, respectively. Interestingly, 5 *Ir*s were detected in the female RT while 56 *Ir*s were detected in the male RT with 46 highly enriched in this male tissue (Additional File 1, Fig. S4b; Additional File 2, Table S22), suggesting a potential critical role in fly reproduction or reproductive behaviors. Given the striking expansions and expression pattern of genes within this family, further functional studies in *Stomoxys* are warranted. Characterizing the gene families involved in olfaction and gustation provides a means to identify compounds that are repellent or attractive to stable flies, facilitating design of behavior-based control technologies.

### Expansion of the long wavelength-sensitive Rh1 opsin subfamily with tuning substitution evidence of diversified wavelength sensitivities

#### Global opsin conservation

As is typical for the generally fast flying calyptrate flies, stable fly adults of both sexes are equipped with large compound eyes in the lateral head and three ocelli positioned in the dorsal head cuticle [94, 95]. Both, achromatic motion tracking and color-specific perception tasks begin with the harvest of photons by members of the opsin class of G-protein coupled transmembrane receptors which differ in their wavelength absorption optima. Our genomic survey in the stable fly revealed conservation of most opsin gene subfamilies observed in *Drosophila* (Fig. 7a; Additional File 1, Fig. S8 and Table S24). This included the UV sensitive opsin paralogs *Rh3*, the blue sensitive opsin *Rh5*, several homologs of the long wavelength (LW) sensitive opsin *Rh1* but a 1:1 ortholog of LW opsin *Rh6*, all of which are expressed in subsets of photoreceptors in the compound eye retina [96]. In addition, we found 1:1 orthologs of the ocellus-specific opsin Rh2 and the recently characterized UV-sensitive, deep brain opsin Rh7 [97] (Fig. 7a; Additional File 1, Fig. S8 and Table S24). Overall, these findings are consistent with the electrophysiological and positive phototactic sensitivity of *Stomoxys* to light in the UV, blue, and green range of visible light [24, 98].

**Figure 7.**
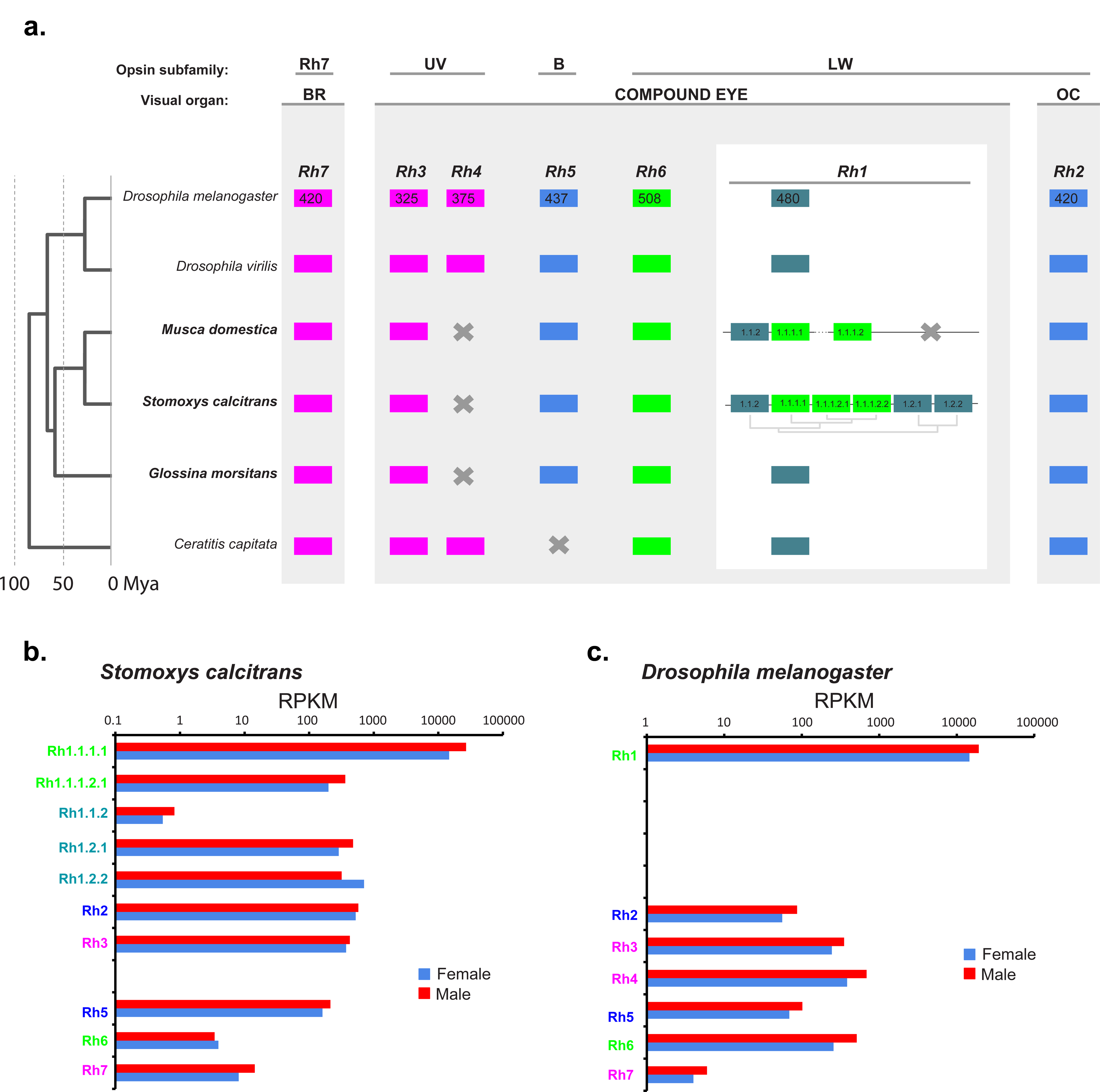
Phylogenetic and genomic organization of dipteran opsin gene relationships. A. Phylogenetic tree of dipteran opsin gene relationships, and genomic organization and evolution of the *Stomoxys calcitrans* Rh1 opsin subfamily [178]. Protein sequences were aligned with MUSCLE [179]. Ambiguous alignment regions were filtered using Gblocks [180] using least stringent settings. Maximum liklihood tree was estimated in MEGA version 6.0 [181] applying the Jones-Taylor-Thornton (JTT) model of amino acid sequence evolution and assuming Gamma Distributed substitution rates across sites with 3 categories. **B.** Transcript abundance differences between *Stomoxys* and *Drosophila* opsins.

*Drosophila* and other higher Diptera including the Mediterranean fruit fly, *Ceratitis capitata*, possess a second UV sensitive opsin gene *Rh4* (Fig. 7) [34, 99], which is not detected in the stable fly genome. The same result was previously obtained in the tsetse fly [33]. As global BLAST searches failed to detect Rh4 orthologs in any calyptrate genome, it can be concluded that the Rh4 opsin subfamily was lost during early calyptrate evolution.

#### An Rh1 opsin gene cluster in muscid Diptera

A unique aspect of the *Stomoxys* opsin gene repertoire is the existence of six homologs of the LW opsin *Rh1* (Fig. 7; Additional File 1, Table S23). Most higher Diptera sampled so far, including related species like the tsetse fly [33] and the black blowfly *Phormia regina* [100], possess a singleton Rh1 gene. Three Rh1 homologs, however, were detected in the *Musca* draft genome [52]. Moreover, in both *Musca* and *Stomoxys*, these Rh1 homologs are closely linked and anchored as a cluster by homologous flanking genes (Fig. 7; Additional File 1, Fig. S9). This suggests that the Rh1 tandem gene clusters of the two species are homologous and date back to an ancestral cluster in the last common ancestor of muscid Calyptratae. Consistent with this, the *Stomoxys* and *Musca* Rh1 homologs formed a monophyletic unit in maximum likelihood trees estimated from amino acid or nucleotide sequence alignments of dipteran Rh1 homologs (Additional File 1, Fig. S10). Moreover, each of the three *Musca* Rh1 homologs grouped with strong support as 1:N orthologs with different members of the *Stomoxys* Rh1 gene cluster (Fig. 7; Additional File 1, Fig. S10). Two of the *Stomoxys* Rh1 genes (Rh1.2.1 and Rh1.2.2), however, lack *Musca* orthologs albeit rooting deeply into the muscid Rh1 gene clade (Fig. 7; Additional File 1, Fig. S10). Integrating the information on gene linkage, it is possible to conclude that the six Rh1 paralogs of *Stomoxys* originated by three early duplications before separating from the *Musca* lineage. While the latter subsequently lost one of the two earliest paralogs, the *Stomoxys* Rh1 cluster continued to expand by minimally one but possibly two subsequent tandem gene duplications (Fig. 7a and Additional File 1, Fig. S9).

#### A tuning site amino acid replacement differentiates two muscid Rh1 paralog subclusters

While exceptional for other higher Diptera, tandem duplicated LW opsin gene clusters have been found in mosquito and water strider species [101, 102]. In both cases, evidence of functional paralog diversification has been detected in the form of amino acid changes that affect opsin wavelength sensitivity, i.e. at tuning sites. Integrating data from butterflies and *Drosophila*, the water strider study identified one high confidence tuning site that very likely affects the blue vs green range wavelength specificity in LW opsins: site 17 based on the numbering system developed for butterflies, which corresponds to residue 57 in *Drosophila* Rh1 [102-104]. Based on this criterion, the three oldest *Stomoxys* Rh1 gene cluster paralogs preserved the blue-shifted wavelength specificity of the Rh1 singleton homologs of other dipteran species (λ_max_ 480nm in *Drosophila*) given their conservation of the ancestral methionine state at tuning site 17 (Fig. 7a; Additional File 1, Fig. S11). The three younger *Stomoxys* Rh1 paralogs, by contrast, share a leucine residue at tuning site 17, which is extremely rare across insect LW opsins. In a survey of over 100 insect LW opsins, it was detected only in the corresponding two Rh1 orthologs of *Musca* in addition to one more distantly related species in thrips (Thysanoptera) [102]. Further, the physicochemical similarity of the tuning site 17 leucine in the three youngest *Stomoxys* Rh1 paralogs to the pervasively conserved isoleucine residue at tuning site 17 in the green-sensitive Rh6 opsins (λ_max_ 515nm in *Drosophila*) represents compelling evidence that this shared derived replacement substitution defines a green-sensitive subcluster in the *Stomoxys* Rh1 paralog group (Fig. 7a; Additional File 1, Fig. S11). Of note, *Stomoxys* is also characterized by an expanded visual sensitivity in the red range [24, 98]. While this aspect could likewise be related to the Rh1 opsin cluster expansion, the red-sensitivity of *Stomoxys* is shared with other calyptrate species, which preserved the ancestral condition of a singleton Rh1 homolog (including *Glossina* and *Calliphora*) [105, 106]. This suggests that the red-sensitivity peak of *Stomoxys* is mediated by accessory filter pigments instead of one of the newly emerged Rh1 gene cluster paralogs.

#### Dramatic transcript abundance differences between muscid Rh1 gene cluster members

As expected, all *Stomoxys* opsin genes were characterized by significant transcript levels in head vs other adult body regions (Additional File 1, Table S24). Moreover, the head tissue derived RNA-seq data provided evidence that a single member of the Rh1 paralog clusters, named Rh1.1.1.1, maintained the ancestral function of Rh1 as the major motion detection specific opsin. The singleton opsin Rh1 of *Drosophila* is expressed in six motion detection-specialized outer photoreceptors (R1-6) per ommatidium while opsins Rh5 and Rh6 are differentially expressed in a single color-vision specialized photoreceptor (R8) per ommatidium [96]. In the *Drosophila* modENCODE expression catalogue, this is reflected in up to 200 fold higher transcript abundance of Rh1 opsin compared to Rh5 or Rh6 in the adult heads of both sexes [107] (Fig. 7c; Additional File 1, Table S24). A similarly massive transcript abundance was detected for the Rh1.1.1.1 homolog of both *Stomoxys* and *Musca*, while the remaining Rh1 cluster member genes were characterized by low to very low transcript levels (Fig. 7b; Additional File 1, Table S24).

The dramatic expression level differences between the Rh1 cluster paralogs in *Stomoxys* may reflect restrictions to smaller subsets of specialized photoreceptors or low level expression throughout the retina. An attractive possibility for the former scenario is the male-specific ‘love spot’ region [108]. In *Stomoxys*, this dorsofrontal expansion of the male eyes, which plays a role in fast flight mating partner pursuit, translates into about 4 percent more ommatidia (∼4,250) compared to females (∼4,050) [94], which may be reflected in the 1.8-fold expression level difference of Rh1.1.1.1 between the adult head transcriptomes of male vs female *Stomoxys* [109]. The presence of similarly enlarged male eyes in other calyptrate species (*Fannia fannia, M. domestica, S. calcitrans, C. erythrocephala, Chrysomya megacephala*), however, suggests that either this male-specific compound eye region was already present in the last common ancestor of calyptrate flies or species are predisposed to evolving this trait [94, 110] (Additional File 1, Fig. S8). The evolutionary origin of the muscid Rh1 cluster is thus not tightly correlated with that of the ‘love spot’ region. Moreover, none of the Rh1 cluster paralogs are expressed in a strictly male-specific manner (Fig. 7b; Additional File 1, Table S24). In contrast, one paralog (Rh1.2.2) is characterized by 2 fold higher expression in the female head (Fig. 7b; Additional File 1, Table S24). Single cell analysis of the Rh1 cluster paralog expression specificity may reveal yet unknown specialized subregions in the stable fly compound eye retina.

#### A shift in Stomoxys Rh5 vs Rh6 opsin transcript abundance compared to Drosophila

A notable difference in the relative opsin transcript levels between *Stomoxys* and *Drosophila* provides tentative evidence that some of the *Stomoxys* Rh1 gene cluster paralogs may have adopted functions in the color-sensitive R8 photoreceptors. These photoreceptors express either the blue sensitive Rh5 opsin or the green-sensitive Rh6 opsin in *Drosophila* [96]. In the *Drosophila* modENCODE expression catalogue, Rh6 transcripts are 3-5 fold more abundant in the whole head transcriptome in comparison to Rh5, consistent with the expression of Rh6 in about 70% of the *Drosophila* R8 photoreceptors (Fig. 7c; Additional File 1, Table S24). In the head transcriptomes of both male and female *Stomoxys*, by contrast, Rh6 transcripts are 2 orders of magnitude less abundant than Rh5 transcripts (Fig. 7b; Additional File 1, Table S24). In the heads of both sexes, Rh6 opsin transcript abundance is the second lowest of all opsin genes and even below that of the deep brain opsin Rh7, which is expressed in only about 20 pacemaker cells in *Drosophila* [97]. The lower transcript abundance of *Stomoxys* Rh6 relative to *Drosophila* could reflect a partial (but pronounced) replacement of ancestral Rh6 expression in R8 photoreceptors by the blue-sensitive Rh5 opsin or members of the Rh1 gene cluster. Of possible significance in this context, stable flies exhibit strong positive phototaxis in response to UV-and blue range light sources [22, 23, 111, 112]. Moreover, positive phototaxis to blue light increases in female flies after fertilization [24]. Whether and how these visual preferences relate to the high ratio of blue (Rh5) vs green-sensitive (Rh6) opsin expression also awaits resolution through single cell expression and wavelength-sensitivity studies of the unexpectedly complex opsin gene repertoire of the stable fly.

### Immune system gene family expansions may reflect adaptation to larval development in microbe-rich substrates

Analysis of the *Stomoxys* genome revealed extensive conservation of immune system signaling pathways coupled with dramatic expansions of some gene families involved in both recognition and effector functions. The insect immune system – best characterized from work in the model organism *Drosophila melanogaster* – includes both cellular defenses (e.g., macrophage-like cells that phagocytose pathogenic microorganisms) and a humoral defense system that results in the production of antimicrobial effector molecules [113]. The humoral immune system can be divided into recognition proteins, which detect pathogenic bacteria and fungi; signaling pathways, which are activated by recognition proteins and result in the translocation of transcription factors to the nucleus to induce gene expression; and effectors, which are (typically) secreted and ultimately act to clear infections.

Previous comparative work suggests that at least some parts of the immune system are deeply conserved across Dipterans and indeed most insects. Genes encoding immune signaling proteins, in particular, are generally preserved as single-copy orthologs across a wide range of insects [33, 52, 114-120], with only rare exceptions [121]. Despite the strong conservation of the basic structure of the main signaling pathways in insect immunity, there is considerable evidence for variation in both the gene content and protein sequence of the upstream inputs (recognition proteins) and downstream outputs (effector proteins) of the immune system (e.g., [116-119, 122, 123]).

We find major components of the Toll, Imd, JAK/STAT, p38, and JNK pathways in the *S. calcitrans* genome (Additional File 1, Table S27), largely conserved as single-copy orthologs [113]. A description and full lists of putative computationally annotated and manually curated immune-related genes in *S. calcitrans* is provided (Additional File 1, Section 9; Additional File 2, Tables S25 and S26). These findings are consistent with previous reports for many other Dipterans, and supports the conclusion that the intracellular signaling mechanisms of innate immunity have been stable during the evolutionary history of Dipterans. In contrast, the gene families encoding upstream recognition proteins and downstream effector proteins tend to be expanded in *S. calcitrans* and *M. domestica* relative to other Dipterans (Table 1).

**Table 1.** Number of Immune-Related Gene Families Annotated by Hidden Markov Models. Numbers in parentheses are those numbers annotated after manual curation of the genome.

|  |  | *Scal* | *Mdom* | *Gmor* | *Aaeg* | *Dmel* |
| --- | --- | --- | --- | --- | --- | --- |
|  | ATT | 11 (12)* | 10 | 4 | 1 | 4 |
|  | DEF | 5 (11)* | 5 | 0 | 4 | 1 |
|  | DIPT | 3 (1)* | 4 | 0 | 1 | 3 |
|  | CEC | 5 (10)* | 12 | 2 | 9 | 5 |
| **canonical effectors** | LYS | 23 | 32 | 4 | 7 | 13 |
|  | TPX | 5 | 6 | 6 | 5 | 8 |
|  | PPO | 19 | 23 | 4 | 25 | 10 |
|  | GPX | 1 | 1 | 0 | 3 | 2 |
|  | HPX | 12 | 12 | 8 | 19 | 10 |
| **non-canonical effectors** | TSF | 4 | 6 | 3 | 5 | 3 |
|  | NIM | 25 | 23 | 10 | 8 | 17 |
|  | PGRP | 17 | 17 | 4 | 10 | 13 |
|  | BGBP | 4 | 3 | 3 | 7 | 7 |
| **canonical recognition** | TEP | 16 | 22 | 4 | 8 | 6 |
|  | CTL | 78 | 41 | 11 | 43 | 38 |
|  | FREP | 49 | 38 | 7 | 34 | 14 |
|  | GALE | 15 | 13 | 8 | 12 | 6 |
|  | IGSF | 1 | 1 | 1 | 0 | 1 |
|  | MD2L | 8 | 12 | 5 | 26 | 8 |
|  | SRCA | 3 | 3 | 2 | 2 | 3 |
|  | SRCB | 15 | 18 | 11 | 13 | 14 |
| **other recognition** | SRCC | 7 | 8 | 4 | 5 | 9 |

Based on Hidden Markov Model profiles and manual curation, we analyzed four canonical recognition families with well-characterized immune roles and an additional eight families with less well-defined roles (Table 1). For three of the four canonical pattern recognition receptor families (PGRP, NIM, TEP), and four of the other families (CTL, GALE, FREP, and SRCB), the *Stomoxys* genome encodes either the most or second-most after *M. domestica*, members among the 5 Dipteran genomes screened (*S. calcitrans* plus *Aedes aegypti, D. melanogaster*,. *domestica*, and *G. morsitans*). A similar pattern holds for downstream effector proteins: the *S. calcitrans* genome encodes either the most or second-most after *M. domestica* for attacins (ATT), defensins (DEF), cecropins (CEC) and lysozymes (LYS). Not unexpectedly, three classes of AMPs were originally characterized in *D. melanogaster* but are missing from the *M. domestica* genome (Metchnikowin, Drosocin, Drosomycin) and are also not detected in the *Stomoxys* genome.

Several of the expanded AMP gene families were found clustered on individual scaffolds, possibly arising from tandem duplications (Additional File 1, Section 9). For example, the 11 *Stomoxys defensin* genes are located on a single scaffold (KQ079966), and these phylogenetically separate into three lineages that are grouped into two regions along the scaffold, i.e. one includes five *defensins* present upstream of the other that includes six *defensins* (Additional File 1, Fig S15). Within the downstream cluster, three genes demonstrate larval-biased expression patterns while the other three appear induced upon blood-feeding in adults (Additional File 2, Table S26). In contrast, *defensin* genes in the upstream cluster are all detected in newly emerged adults and are upregulated in response to blood-feeding. This expression pattern is consistent with that reported for *Stomoxys midgut defensin 1* (*Smd1*) and *Smd2*, which were present in this upstream cluster [124].

The *Stomoxys* genome encodes 17 PGRPs, 11 in the short subfamily, 5 in the long subfamily, and 1 ambiguous. Of the members of the short subfamily, six – all orthologs of the PGRP-SC gene family in *D. melanogaster* – appear to be expressed exclusively in larvae and, based on sequence properties and conservation of residues required for amidase activity [125], are predicted to be both secreted and catalytic (Additional File 1, Fig. S13). PGRP-SC genes in both *D. melanogaster* [126] and *M. domestica* [127] are also expressed in larvae, but this expression is not exclusive suggesting the possibility that larval-specific expression may be an *S. calcitrans* innovation.

In combination with the previously reported expansions of many effector and recognition immune components in the house fly [52, 116], our analysis of the *Stomoxys* genome suggests that Muscidae likely have expanded the diversity of their immune repertoires, sometimes dramatically, despite differences in adult feeding ecology (blood feeder vs generalist). One hypothesis is that the shared diversification of immune receptors and effectors is driven by larval ecology (e.g., the shared requirement for bacteria during development and the septic environment larvae inhabit), while additional *M. domestica* specific expansions (e.g. in TEPs) are accounted for by the saprophytic adult feeding behavior of that species.

### Immunomodulatory and anti-hemostatic products are prominent in the sialome

Blood-feeding insects salivate while probing their host skin for a blood meal. Development of a sophisticated salivary potion that disarms their hosts’ hemostasis is among the adaptations to blood feeding found in hematophagous animals [128, 129]. Blood clotting inhibitors, anti-platelet compounds, vasodilators and immunomodulators are found in salivary gland homogenates or saliva of blood sucking arthropods [129]. To determine the genes associated with salivation, we mapped the reads from four RNA-seq libraries (male and female salivary glands -SG, as well as male and female whole bodies – WB) to the *S. calcitrans* predicted gene set. We used an ?^2^ test to identify those that were significantly over-expressed in SG (Fig. 8), as in [130] (Additional File 2, Tables S2 and S29). A subset of SG transcripts with 100-fold higher expression than in WB was analyzed in more detail (Additional File 2, Table S30). The SG 100-fold overexpressed set was comprised of 139 transcripts, 18 of which were found to be splice variants, or identical to other transcripts, as verified by their scaffold coordinates. The non-redundant set comprised of 121 transcripts was classified into three major groups: Putative Secreted, Putative Housekeeping and Unknown; these groups were further classified into finer functional categories (Additional File 2: Table S29).

**Figure 8.**
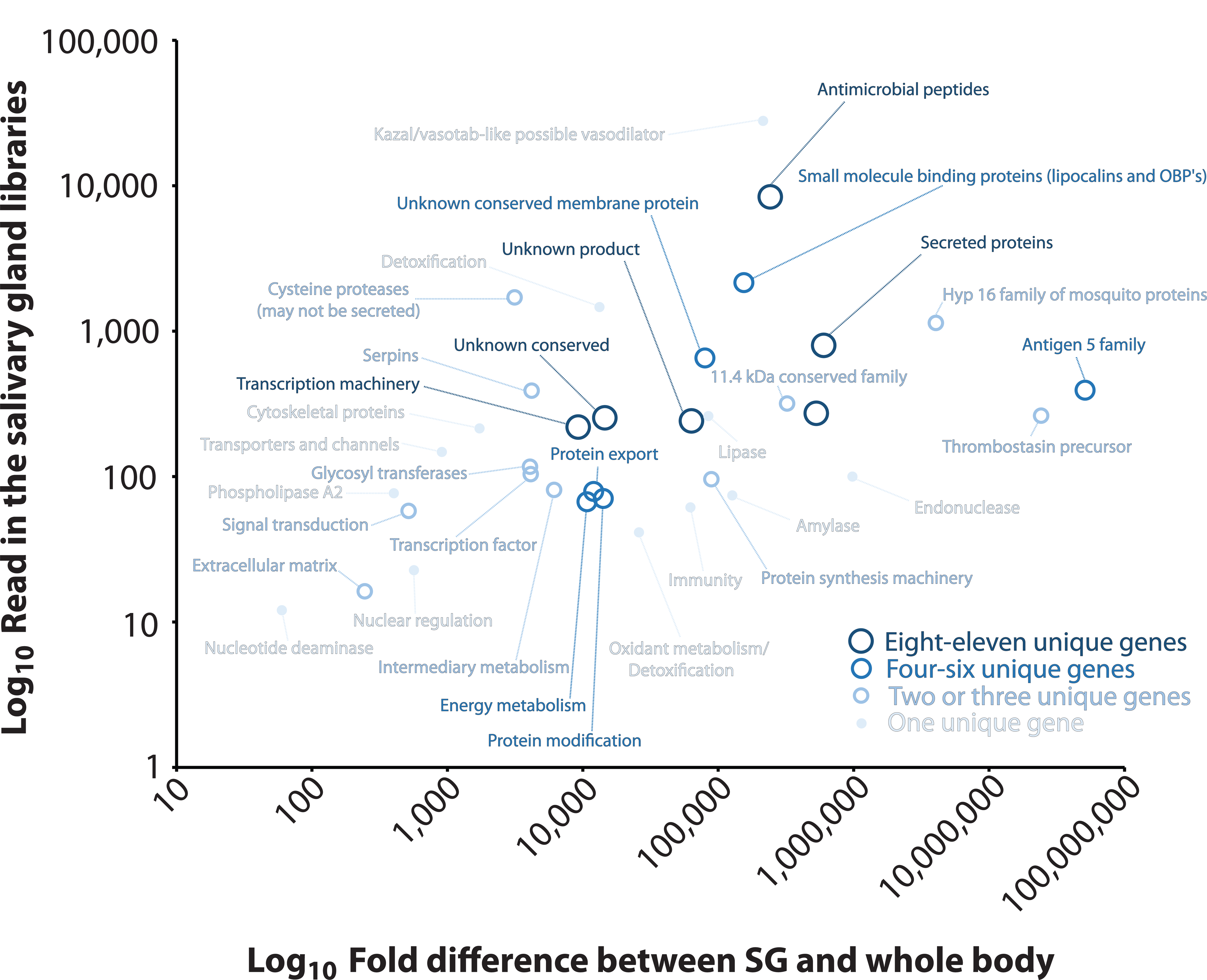
Analysis of salivary gland biased genes in *Stomoxys calcitrans*. Number of Illumina reads versus fold enrichment in the salivary gland related to the whole body. Each point represents the average among all genes in that specific category. Expression levels are based on results in Additional File 2, Table S29.

In congruence with the SDS gel of *S. calcitrans* SG [131], the antigen 5 family returned 62% of the total reads mapped to the 121 SG-enriched transcripts (Fig. 8). Members of this family in *S. calcitrans* may function as inhibitors of the classical complement system [132]. Thrombostasin [133] members are represented by two transcripts (29% of reads with strong gel bands, [131]), which are precursors for anti-thrombin peptides previously identified in *S. calcitrans*. They accrued 29 % of the reads and are strongly represented in the gel bands. The Hyp 16 family of peptides (unknown function, 4.9% of accrued reads) and one transcript encoding an endonuclease (1.2% of accrued reads) were also noted. Together, these groups of transcripts account for 97% of the reads that are over expressed in the salivary glands of *S. calcitrans.*

There was a wide variety of other transcripts represented in the last 3% of the reads, and serine proteases, nucleotide deaminase, amylase, phospholipase A2 and lipases were found enriched in the *S. calcitrans* sialome. These enzymes are also found enriched in other sialomes and their functions have been reviewed [129, 134, 135]. Two of eight serine proteases were found 5-15 times overexpressed in female salivary glands when compared to male glands. These two products produce best matches to vitellin-degrading proteases from *M. domestica* (XM_005191887.2) and may be indeed female enriched enzymes that were hitchhiked to the salivary set due to their similarities to overexpressed salivary enzymes. No other peptides were found above five-fold expressed in either salivary gland gender.

Several antimicrobial peptides appeared enriched in the *S. calcitrans* sialome, including lysozyme, attacins, defensins, diptericin, a GGY rich peptide and sarcotoxin. Of these, only the GGY peptide and diptericin were identified in the Sanger-based sialome description [131]. These peptides may help to control microbial growth in the ingested blood. Regarding polypeptides with anti-proteolytic activity, in addition to thrombostasin precursors discussed above, two CDS coding for serpins (however, with very low expression) and one coding for a Kazal domain-containing peptide (accruing 0.3 % of the reads) were identified. While serpins may modulate clotting and inflammation-related proteases, the Kazal domain peptide may be related in function to vasotab, a vasodilatory peptide from a tabanid fly [136].

Finally, 24 transcripts accruing 0.19 % of the reads could not be functionally classified and were thus assigned to the “unknown” class. These include membrane proteins (XP_013114823.1 and XP_013117270.1) that are over one thousand fold over expressed in the salivary glands (the first of which was identified in the Sanger sialotranscriptome) and are attractive targets for gene disruption experiments to elucidate the contribution of these proteins to the salivary function of *S. calcitrans*.

### Expanded cytochrome P450 gene family may enhance metabolic detoxification of insecticides

The family of *Stomoxys* cytochrome P450s (CYPs) identified from the current genome assembly represent a substantial expansion relative to other sequenced dipteran genomes. Arthropod CYPs have diverse roles in insect physiology, including ecdysteroid biosynthesis and xenobiotic detoxification [137, 138]. The CYP gene family size varies among insects with dipterans having large arrays, i.e. 145 in *Musca*, 86 in *Drosophila*, and 77 in *Glossina*. The 214 *Stomoxys* CYPs that were identified encode representatives from each of the CYP clans that are typically found in insects, *i.e.* mitochondrial, CYP2, CYP3, and CYP4 (Fig. 9; Additional File 2, Table S31). As in *Musca*, expansions in *Stomoxys* were primarily in clans 3 and 4. The CYP4 clan (62 genes) was represented by the CYP4 (51 genes) family, while the CYP3 clan (107 genes) comprised the largest expansion of CYPs in *Stomoxys*, predominated by the CYP6 (81 genes) and CYP9 (16 genes) families. Together, members of the CYP4 and CYP6 families represent 62% of the *Stomoxys* CytP450s, which is comparable to what was observed in *Musca* [52]. Upregulation of genes in the CYP4, CYP6, and CYP9 families has been associated with resistance to spinosad and pyrethroid insecticides in *Musca, Anopheles*, and *Drosophila* [139-141], but this has yet to be investigated in *Stomoxys*. Interestingly, tandemly duplicated arrangements (“blooms”) of 11 CYP4D (scaffold KQ080140), 18 CYP6A (scaffold KQ080692), and 16 CYP9F (scaffold KQ080085) genes are present in the *Stomoxys* genome. Given that *Stomoxys* inhabits conventional livestock production settings that utilize chemical fly control measures, the initial gene duplication that eventually lead to these expanded clusters may have been favored because of environmental exposure to xenobiotic pressure [142].

**Figure 9.**
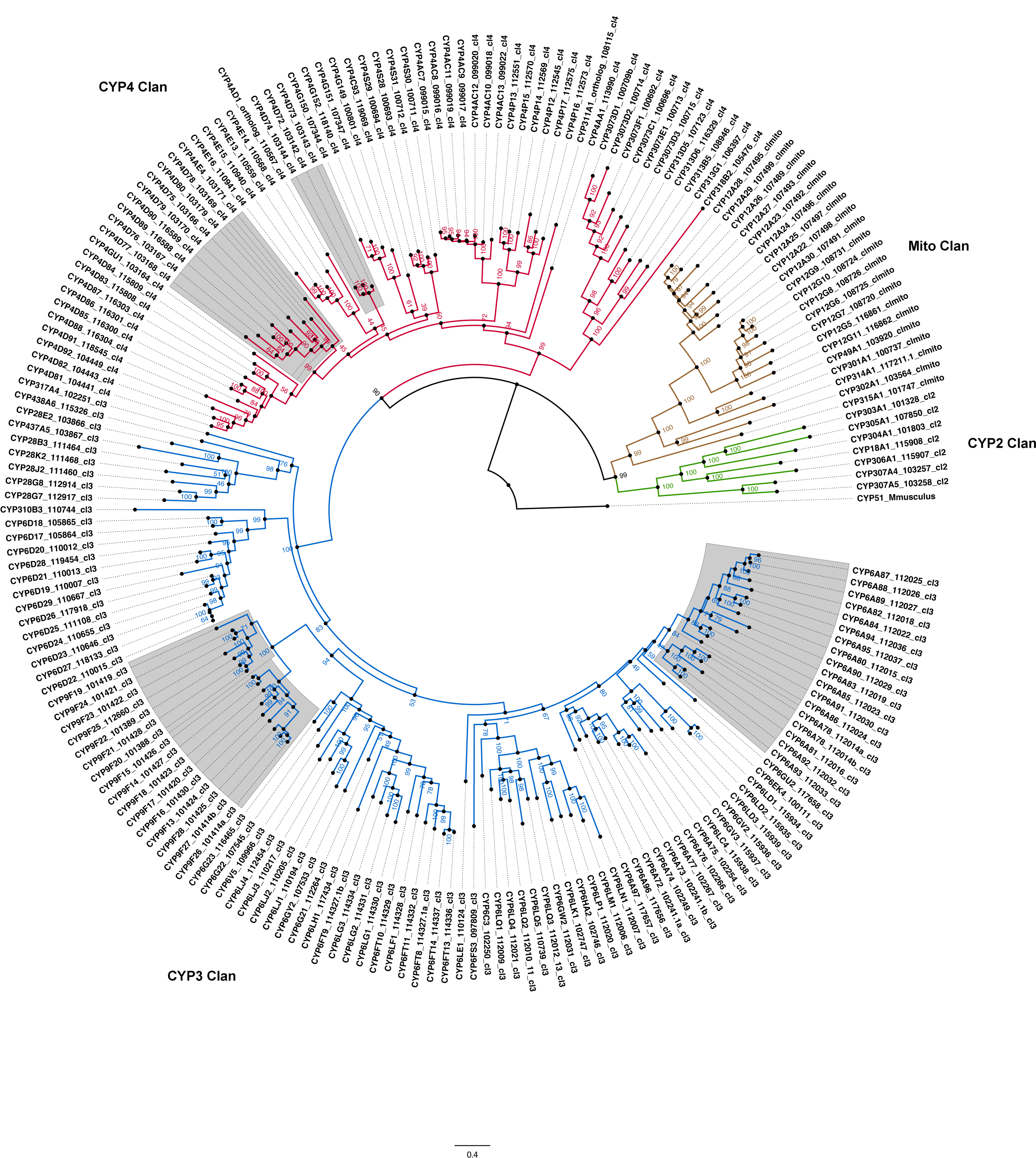
Phylogenetic analysis of cytochrome P450 genes from *Stomoxys calcitrans*. Amino acid sequences from each family were aligned with the MUSCLE algorithm [179], and the alignments trimmed with the trimAl tool using the –strictplus option [182]. The trimmed alignment was used to construct a maximum likelihood phylogeny, rooted with *Mus musculus* CYP51 as the outgroup, with the web server version of IQ-TREE software (best-fit substitution model, branch support assessed with 1000 replicates of UFBoot bootstrap approximation; [177]). The CYP clades are presented in different colors, and CYP gene clusters that are found in tandem within the genome are shaded in grey. P450 gene names were assigned based on comparative analyses.

Metabolic detoxification of insecticides can be further mediated by the carboxylesterase and glutathione-S-transferase gene families, members of which were identified from the *Stomoxys* genome assembly and were comparable to the gene families from *Drosophila* and *Musca* (Additional File 1, Section 10, Figs. S16, S17, S19). Further, 26 members of the Cys-loop gated ion channel (CysLGIC) superfamily, targets for several classes of insecticides [143, 144], were identified in *Stomoxys* (Additional File 1, Section 10, Figure S20).

Stable fly population reduction relies on integrating cultural control practices, trap deployment, and application of insecticides, although the latter is considered less effective for control of adult populations given that stable flies do not spend as much time on their mammalian host and can disperse across long distances. Interactions of stable fly adults with crops and various structures to which insecticides have been applied provides avenues for exposure, and use of insecticide fogging in outbreak situations contributes to this exposure.

There are limited studies evaluating levels of stable fly insecticide resistance in field populations, but anecdotal reports of product failure for control of stable fly populations in Brazil (Thaddeu Barros, pers. comm.) and Costa Rica (Arturo Solorzano, pers. comm.) underscores its importance. Pyrethroid resistance was reported in Europe and the US [145-147], and the phenotype was attributed, in part, to target site insensitivity at the *knockdown resistance* (*kdr*) locus of the *voltage-sensitive sodium channel* gene [148, 149]; there is evidence indicating additional mechanisms are involved. Metabolic detoxification of insecticides may contribute to pyrethroid resistance in stable flies, especially given the robust expansion of the CytP450 gene family. Descriptions of these insecticide resistance (ion channel and enzymatic) gene families facilitates further functional studies to define these mechanisms in stable fly populations.

### Microbiota and Lateral Gene Transfers

Similar to mosquitoes, stable flies will consume blood and nectar for nourishment [25, 83, 150]. This is different to the closely-related tsetse flies, which are obligate blood feeders. Due to this limited food source, tsetse flies harbor an obligate symbiont, *Wigglesworthia glossinidae*, that provides B vitamins that are present at low levels in blood [33, 151, 152]). DNA sequencing, along with analysis of the assembled *Stomoxys* genome, did not reveal a distinct microbial symbiont. A preliminary study of culturable bacteria among adult *Stomoxys* collected from four US (Texas) dairies revealed that there are a variety of bacteria associated with adult stable flies (Fig. 10; Additional File 1, Section 11; Additional File 2, Table S30). Among those cultured, *Aeromonas* was the most prevalent. *Aeromonas* sp. are frequently found in aquatic environments, such as irrigation water, and have been cultured from arthropods [153]. Similar to mosquitoes, harbored bacterial communities are likely acquired during the larval stage or as adults during ingestion of nectar or water sources, as strict blood feeders usually have reduced gut microbiota [154-156]. In addition, the *Stomoxys* bacterial communities have similarity to those isolated from other flies associated with filth and decomposition (e.g. blow flies and other related species, [157, 158]), suggesting these bacterial components may be retained from larval development or acquired by adults when visiting sites for oviposition. Of particular interest, multiple potential pathogens were cultured, such as *Staphylococcus* sp. and *Bacillus* sp. associated with bovine mastitis and *Vibrio* sp., suggesting the potential of this fly to act as a reservoir for pathogens. Unlike bacterial community surveys of stable flies from dairies in Brazil, *Salmonella* and *Escherichia coli* were not prevalent in this US sampling [159, 160].

**Figure 10.**
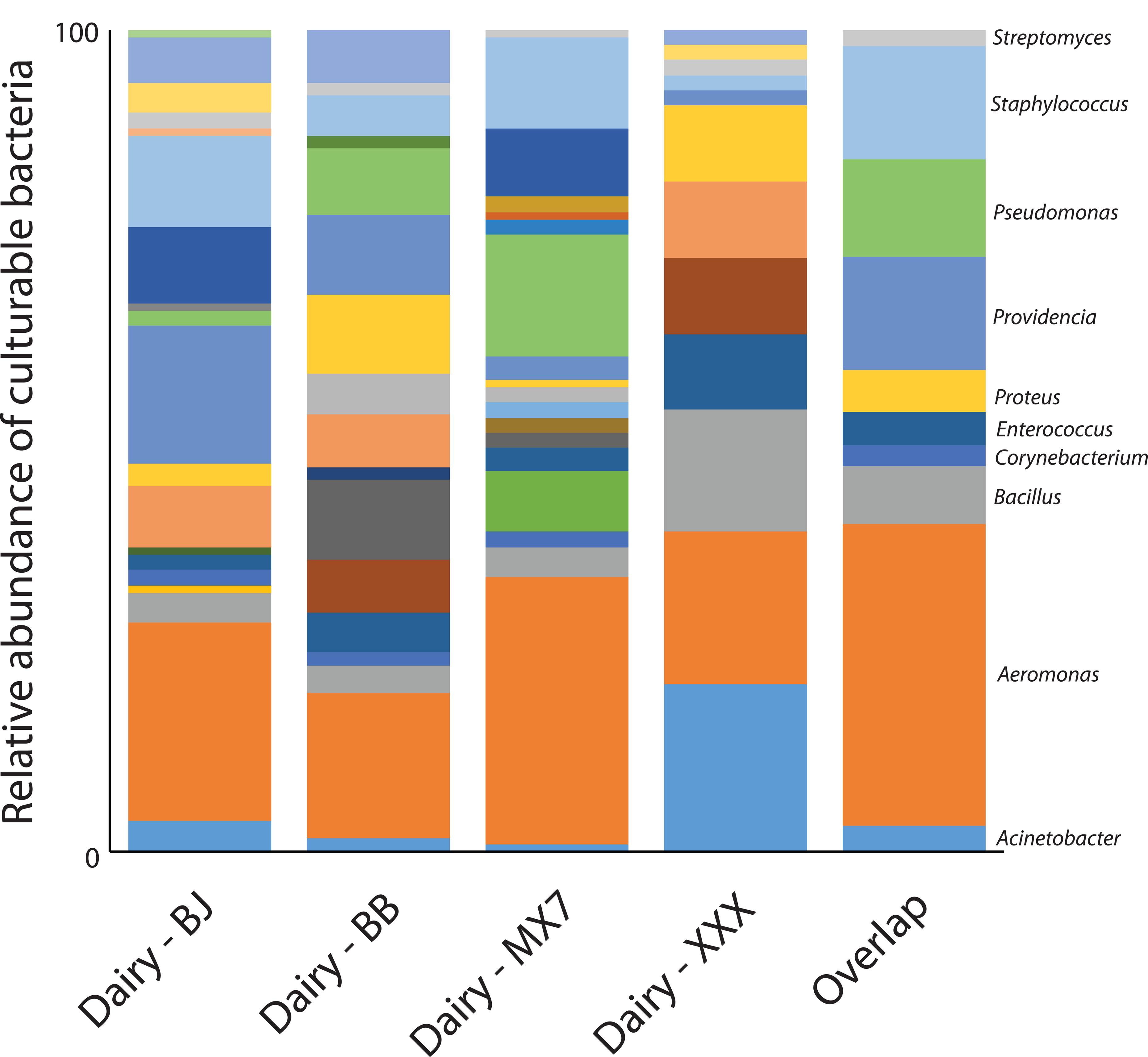
Microbiome analyses of culturable bacteria. Survey of the culturable bacteria associated with stable flies from four separate dairies in Texas. Overlap represents species present at all four sites. Complete list is available in Additional File 2, Table S32.

A pipeline was utilized for detecting bacterial to insect lateral gene transfers (LGT), and three candidate LGTs were detected (Additional File 1, Section 12). All three are derived from *Wolbachia*, a common endosymbiont found in arthropods [161] that infects 40-60 percent of insect species [162, 163]. While they are a common source of LGTs, likely due to their association with the germline of their insect hosts, the *Stomoxys* strain used for the genome sequencing is not infected with *Wolbachia* and there are no reports of natural occurrence of *Wolbachia* in *Stomoxys* populations. Presence of these LGTs, then, may be evidence of incomplete infection of the species or to LGT events from a past *Wolbachia* infection, that has been subsequently lost in the species. There is no evidence that any of the three LGTs have evolved into functional protein coding genes in *Stomoxys*, although one has detectable expression and occurs within the 3’ UTR of XM_013245585, which encodes a transcription factor containing a basic leucine zipper domain. Whether expression of the LGT is biologically significant is unknown, and further studies of these three LGTs are warranted.

### Evidence for transcription factors with putative role in regulation of salivation and reproduction

To determine transcription factors (TFs) that might control specific gene expression profiles, we first predicted TF-encoding genes by identifying putative DNA binding domains (DBDs), using a previously described approach [32, 37]. These analyses resulted in 837 predicted TFs, with the highest number coming from the C_2_H_2_ zinc finger and homeobox structural families (Fig. 11), consistent with previously analyzed insect genomes [32, 36, 37].

**Figure 11.**
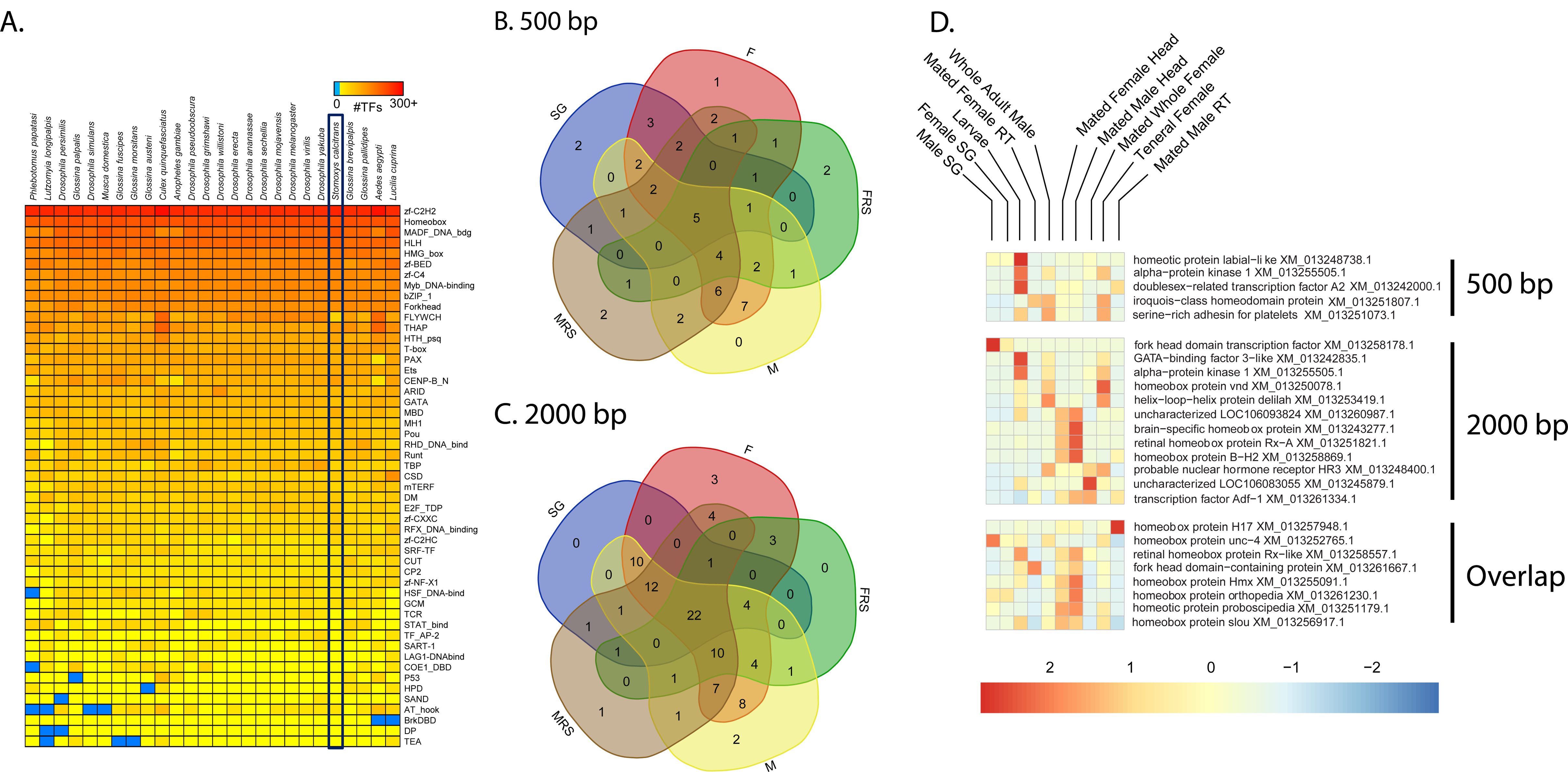
Transcription factors associated with stable flies. A. Number of transcription factors identified in *Stomoxys calcitrans* compared to other flies. (**B** and **C**) Overlap between transcription factors with increased binding sites in differentially expressed genes that have noted expression in the same tissue (F, teneral female; FRS, female reproductive system; M, male; MRS, male reproductive system; SG, salivary glands). (**D** and **E**). Expression of specific TFs associated with female, male, and salivary glands among multiple tissues and developmental stages.

We next predicted DNA binding motifs for as many of these putative TFs as possible using a previously developed method [164]. In brief, the percent of identical amino acids was calculated between each *Stomoxys* TF and each eukaryotic TF with a known motif, with values exceeding a TF family-specific threshold resulting in “inferred” motifs for the *Stomoxys* TFs. For example, the DBD of the uncharacterized XP_013101333 protein is 92.3% identical to the DBD of the *Drosophila melanogaster* gene *cropped* (FBgn0001994). Since the DNA binding motif of cropped has already been experimentally determined, and the cutoff for the bHLH family of TFs is 60%, we can infer that XP_013101333 will have the same binding motif as cropped. This procedure resulted in inferred motifs for 285 of the *Stomoxys* TFs (34%).

We then performed TF binding site motif enrichment using promoter regions for groups of genes with similar gene expression patterns in our RNA-seq experiments (see Methods). Promoters were defined as either 500 or 2000 bp upstream of the predicted transcription start site for each gene. We restricted the search to gene set/motif pairs with significant enrichment, and further filtered gene set/motif pairs to cases where (1) the given motif was present in at least 60% of the promoters of the gene set, (2) the given motif was present in less than 20% of all gene promoters, and (3) the difference between the presence of the motif in the gene set and promoters of all genes exceeded 40% (see Methods). Lastly, expression of each TF was verified in specific tissues using our RNA-seq datasets.

Based on these criteria and comparative analyses between samples, seven and nine TFs, respectively, were enriched in SG for the 2000 bp and 500 bp promoter regions (Fig. 11). Based on the 500 bp promoter regions, two specific TFs, *proboscipedia* (XM_013251179) and *orthopedia* (XM_013261230), likely regulate SG-based transcript expression. These two TFs have been associated with head and salivary development in *Drosophila* [165, 166], and the increased binding sites and specific expression profile suggest a role in saliva production.

Male-and female-enriched analysis based on stage and tissue specific RNA-seq analyses identified TF targets in each of the 500 bp and 2000 bp promoter regions (Fig. 11). The four most likely TFs associated with female specific genes are XM_013251807.1 (*iroquois-class homeodomain protein*) and XM_013252765.1 (*Unc4 homeodomain protein*) based on the 500 bp promoter region and two other likely homeodomain proteins, XM_013245879.1 (uncharacterized) and XM_013261334.1 (uncharacterized), in the 2000 bp regulatory region. The latter have high expression in females and female reproductive tract (Fig. 11). Two TFs within the 500 bp promoter region were for male enriched genes XM_013258869.1 (*BarH-2*) and XM_013251073.1 (uncharacterized), both of which are highly expressed in males, male heads, and/or the male reproductive system. When expanded to the 2000 bp promoter region, two additional putative TFs related to male enriched genes are XM_013260987.1 (uncharacterized) and XM_013257948.1 (drop), which are both highly expressed in male samples. The identification of these TFs could provide novel targets for the control of stable fly reproduction or the prevention of feeding.

## CONCLUSIONS

Our analyses reveal unique aspects related to stable fly biology, including molecular mechanisms underlying reproduction, chemical and host detection, feeding, and immune responses. These combined studies provide substantial advancement in the understanding of *Stomoxys* biology and provide the functional genomic resources to develop novel control mechanisms for this livestock pest. Importantly, our studies advance the knowledge of stable fly genomics and genetics to those of other non-model, but extremely important, dipterans such as tsetse and house flies. Recognizing expanded vision associated genes and chemosensory factors will inform the development of behavior modifying compounds and/or strategies, i.e., repellents, and enhance manipulation of visual attraction to improve traps for population suppression. Unique proteins, including transcription factors, that are associated with reproduction, feeding, and immunity will be ideal targets for gene editing strategies that modify sex-specific and tissue-specific genes to aid in population suppression. Further, defining the specific classes of genes that account for stable fly resistance to insecticides will enable the design of diagnostic tools to monitor insecticide resistance in field populations.

## METHODS

### Genome Sequencing, Assembly, and Annotation

Multiple male (F7 generation from inbred line 8C7A2A5H3J4) individual DNA isolates were provided as a pool in TE buffer. The sequencing plan followed the recommendations provided in the ALLPATHS-LG assembler manual [167]. This model requires 45x sequence coverage each of fragments (overlapping paired reads ∼ 180bp length) and 3kb paired end (PE) reads as well as 5x coverage of 8kb PE reads. For fragments and all jumping libraries (3 and 8kb) we used a DNA sample pooled from approximately 10 male individuals. Total assembled sequence coverage of Illumina instrument reads was 66X (overlapping reads 39x, 2kb PE 24.5x, 6kb PE 2.5x) using a genome size estimate of 900Mb reported by the ALLPATHS-LG software (Broad Institute). This first draft assembly was referred to as S_calcitrans 1.0. In the S_calcitrans 1.0 assembly small scaffold gaps were closed with Illumina read mapping and local assembly, and scaffolding was improved using SSPACE [168]. Contaminating contigs, trimmed vector in the form of X’s and ambiguous bases as N’s in the sequence were removed. NCBI requires that all contigs 200bp and smaller be removed. Removing these contigs was the final step in preparation for submitting the 1.0.1 assembly. The final *S. calcitrans*-1.0.1 assembly is made up of a total of 12,042 scaffolds with an N50 scaffold length of over 504kb (N50 contig length was 11kb). The total scaffold assembly including gaps and single contigs scaffolds spans over 971Mb. Data for the *S. calcitrans* genome have been deposited in the GenBank Bioproject database under the accession code PRJNA188117. The genome assembly has been deposited to GenBank under the accession GCA_001015335.1. RNA-seq datasets used in gene prediction have been deposited to the NCBI Sequence Read Archive under the accession codes SRX995857 -5860, SRX229930, SX229931, and SRX275910 (Additional File 2, Table S1). Methods related to the annotation are described within Additional File 1, Section 1.

### RNA-seq analyses

In conjunction with the genomic sequencing, RNA-seq analyses were performed to examine specific transcript differences between different stages and tissues (Additional File 2, Tables S5 – S13). RNA was extracted with the use of TRizol. DNA contamination was reduced via DNase treatment according to methods previously described [169, 170]. Briefly, RNA-seq datasets were trimmed and low-quality reads were removed with trimmomatic. Each dataset was mapped to the to the predicted gene sets (NCBI Annotation Release 100) using CLC Genomics (Qiagen). Reads were mapped with the following parameters: 95% match for over 60% of the read length with only two mismatches allowed. Reads alignments were converted to per million mapped to allow comparison between RNA-seq data sets with varying coverage. Expression was based upon transcripts per million (TPM). A Baggerly’s test followed by a Bonferroni correction at 0.01 (number of genes × α value) was used to establish significance. Each RNA-seq dataset was validated by qRT-PCR (Additional File 1, Section 3). Gene ontology assessment for specific groups were conducted by the use of gProfiler following conversion of *Stomoxys* gene IDs to *D. melanogaster* gene IDs [171]. Salivary gland RNA-seq analyses were conducted based on methods used in previous studies on insect vectors [130].

### Transcription factor analyses

To assess potential transcription factors regulating tissue and sex-specific expression, TFs were identified according to previously developed methods in other insect systems [32, 37]. Enriched TF binding motifs were identified in the 500 and 2000 bp regions upstream of the putative transcription start site using the HOMER tool [172] supplemented with the *Stomoxys* inferred binding motifs obtained from the CisBP database (build 0.90).

### Microbiome analyses and LGT Prediction

To identify culturable bacterial communities harbored by adult stable flies, fly specimens were collected at each of four Texas dairies in April and June 2015 (Lingleville and Comanche, Texas). Twenty flies per site per date were collected by aerial sweep nets in the area surrounding each dairy’s milking parlor. Within 4 hours, whole flies were surface sterilized in 1% sodium hypochlorite for 15 minutes, followed by two washes in 70% ethanol and three rinses in sterile water. Individual flies were macerated in Butterfield’s phosphate buffer, and the homogenate was diluted and plated on tryptic soy agar. Individual, morphologically distinct colonies were selected, suspended in Butterfield’s phosphate buffer, and the DNA isolated by rapid boiling. These DNAs were used as template in 16S PCR amplification with a universal primer pair (16SEub_61F: 5’ – GCTTAACACATGCAAG – 3’; 16SEub_1227R: 5’ – CCATTGTAGCACGTGT – 3’). Individual amplicons were sequenced in both directions and the sequences assembled. Data were processed and full-length sequences (N=170) were analyzed in mothur, v 1.38.1 [173]. Sequences were aligned using the silva.seed_v128.align file and were subsequently clustered and classified at a 97% similarity cut-off, average neighbor, silva.nr_v128.align/.tax.

A DNA based computational pipeline was used to identify “contaminating” bacterial scaffolds and bacterial to *Stomoxys* lateral gene transfer (LGT) candidates in the *Stomoxys* genome assembly. The pipeline was originally developed by Wheeler et al. [174] that has subsequently been modified. Details of the pipeline are provided in Poynton et al. [175] and Panfilio et al. [176], which have been summarized in Additional File 1, Section 12.

## Declarations

### Ethics Approval and Consent to Participate

Not applicable

### Consent for Publication

Not applicable

### Availability of Data

All genome sequence data are publicly available at the NCBI BioProject: PRJNA188117, and RNA-Seq transcriptome data at BioProject: PRJNA288986, with the genome assembly at NCBI accession number GCA_001015335.1. RNA-seq datasets used in gene prediction have been deposited to the NCBI SRA site under the accession codes SRX995857-5860, SRX229930, SX229931, and SRX275910. Annotation and gene model data are available at the Apollo instance hosted on VectorBase (https://www.vectorbase.org/organisms/stomoxys-calcitrans; https://www.vectorbase.org/apollo).

## Footnote

This article reports the result of research only. Mention of trade names or commercial products in this publication is solely for the purpose of providing specific information and does not imply recommendation or endorsement by the U.S. Department of Agriculture. The USDA is an equal opportunity provider and employer.

## Authors’ Contributions

PUO and JBB organized and directed the sequencing, analysis, and manuscript development. Grant funding was obtained for this project as part of the *Glossina* genome project by SA. Sequencing and assembly of the genome was performed by WCW and RKW. Hosting of the genome for annotation was provided by GLM, SJE, and DL. Gene predictions were performed by TDM. Male and female reproductive analyses were conducted by GMA and JBB. Transcription factor analyses were conducted by XC, ENJ, MTW, and JBB. BUSCO and phylogenetic analyses were conducted by EOM, ECJ, and JBB. JMCR, JBB, and PUO collected samples and performed sialome analyses. Chemosensation associated genes were annotated and analyzed by HMR, MD, GB, and PUO. Cuticle proteins were analyzed by AJR and JBB. Vision associated genes were analyzed by MF and JD. CJH and JBB assisted in the annotation of aquaporins. Annotation of CEGMA and autophagy genes was conducted by KJO and JBB. Immune genes were annotated and analyzed by TBS, DN, and PUO. DBT contributed to the writing of the introduction and interpretations of the results. Muller elements were assigned and analyzed by RPM. Bacterial community was assessed by SLS and PUO. Cytochrome P450s were analyzed by DRN. Carboxylesterases, GSTs, and CLGICs were analyzed by PUO. JHW conducted analyses of lateral gene transfers. OrthoDB analyses were conducted by EZ, RMW, and PI. Transposable elements and genomic repeats were analyzed AMT, SHS, CJC, and TJR. PUO, JBB, GMA, MF, RPM, DN, TJR, JMCR, HMR, TBS, AMT, DBT, MTW, RMW, and JHW wrote and edited the manuscript. All authors read and approved the final manuscript.

## Acknowledgements

We thank The Genome Institute at Washington University for library construction and sequencing. JHW thanks S Cheng for assistance with the LGT pipeline.

