## Additional File 1 for "Functional genomics of the stable fly, *Stomoxys calcitrans*, reveals mechanisms underlying reproduction, host interactions, and novel targets for pest control"

***Stomoxys calcitrans* Genome Sequencing Project**

**SUPPLEMENTARY INFORMATION**

**Contents**

1. DNA and RNA for Sequencing, Genome Sequencing and Assembly, In silico Annotation, and Community Curation	3
2. OrthoDB and RNASeq Analyses	5
3. RNASeq validation by quantitative real-time PCR (RT-qPCR)	6
4. Stable Fly Chromosome Arm Assignments	7
5. Presence and enrichment of repeat elements

in the *Stomoxys calcitrans* genome	9

6. *Stomoxys* Reproductive Biology	10
7. *Stomoxys* Chemosensory Gene Families	11
8. *Stomoxys* Vision	33
9. Immune System	38
10. *Stomoxys* Insecticide Resistance Gene Families	48
11. Bacterial Communities Harbored by Stable Flies from Texas Dairies	55
12. Lateral Gene Transfer	56
13. *Stomoxys* Aquaporin Proteins	57
14. *Stomoxys* cuticle proteins	58

**Additional File 1, Supplementary Figures**

| **Figure S1.** | Phylogenetic placement and genomic comparisons for *Stomoxys calcitrans* and other fly species |
| --- | --- |
| **Figure S2.** | Pearson's correlation of RNA-seq and RT-qPCR results |
| **Figure S3.** | *Stomoxys calcitrans* Gustatory Receptor Gene Family |
| **Figure S4.** | *Stomoxys calcitrans* Ionotropic Receptor Gene Family |
| **Figure S5.** | *Stomoxys calcitrans* Odorant Binding Protein Gene Family |
| **Figure S6.** | Duplicated OS-E-like and an OS-X orthologue in *Stomoxys* and *Musca* |
| **Figure S7.** | *Stomoxys calcitrans* Odorant Receptor Gene Family |
| **Figure S8.** | Maximum likelihood tree of dipteran opsin gene relationships |
| **Figure S9.** | Genomic organization and evolution of the S. calcitrans Rh1 opsin subfamily |
| **Figure S10.** | Phylogenetic analysis of the Stomoxys Rh1 gene cluster |
| **Figure S11.** | Analysis of tuning site 17 variation in Stomoxys and Musca Rh1 paralogs |
| **Figure S12.** | Maximum likelihood phylogenetic tree of PeptidoGlycan Recognition Protein (PGRP) sequences from *S. calcitrans* (red), *M. domestica* (black), and *D. melanogaster* (blue) |
| **Figure S13.** | Alignment of *Stomoxys* PGRP-S sequences with characterized *D. melanogaster* PGRP-SC1 (C0HK98) and –SC2 (Q9VX2) and N-acetylmuramoyl-L-alanine amidase (P00806). |
| **Figure S14.** | Alignment of *Stomoxys* PGRP-L sequences with characterized *D. melanogaster* PGRP-L proteins and N-acetylmuramoyl-L-alanine amidase. |
| **Figure S15.** | *Stomoxys calcitrans* genomic scaffold housing 11 defensin gene models, and phylogenetic relationship of defensins |
| **Figure S16.** | Phylogenetic relationship of catalytic carboxyesterases from *Stomoxys* relative to *Musca* and *Drosophila*. |
| **Figure S17.** | Phylogenetic analysis of carboxylesterases with a role in neuronal development |
| **Figure S18.** | Phylogenetic analysis of cytochrome P450 genes from *Stomoxys* |
| **Figure S19.** | Phylogenetic analysis of glutathione-S-transferases from *Stomoxys* relative to *Musca* and *Drosophila*. |
| **Figure S20.** | Phylogenetic analysis of Cys-Loop Ligand Gated Ion Channels from *Stomoxys* relative to *Musca* and *Drosophila*. |

**DNA and RNA for Sequencing, Genome Sequencing/Assembly, Automated Annotation, BUSCO Assessment, and Community Curation**

**Contributors:** Wes Warren, Terence Murphy, Garreth Maslen, Scott Emrich

*Genomic DNA sequencing and assembly.*Total genomic DNA was isolated from pooled, newly emerged/unfed, adult males of a *Stomoxys calcitrans* line that resulted from single paired mating for 7 generations (Sc8C7A2A5H3J4). High quality/ high molecular weight DNA was isolated from the pooled flies using the Genomic-tip purification column and the associated buffer kit (QIAGEN, Valencia CA), and samples were processed according to the protocol for tissue based DNA extraction. The pooled DNA isolates were utilized for sequencing on Illumina^®^ HiSeq2000 instruments. The sequencing plan followed the recommendations provided in the ALLPATHS-LG assembler [1]. Using this model, we targeted 45x sequence coverage each of fragments (overlapping paired reads ~180bp length) and 3kb paired end (PE) sequences as well as 5x coverage of 8kb PE sequences. The first draft assembly scaffold gaps were closed where possible with mapping assembly input sequences (overlapping paired reads ~180bp length) and local gap assembly [2]. Contaminating sequences and contigs 200bp or less were removed. The genome assembly, Stomoxys_calcitrans-1.0.1, was made publicly available in the Genbank sequence database, accession number GCF_001015335.1. All raw data used in assembly were made publicly available, and accession numbers are summarized in Additional File 2, Table S1.

*RNAseq data.* A variety of developmental stages and dissected tissues were collected for total RNA isolation (Additional File 2, Table S1). Specifically, RNA collected from whole females (teneral and mated, reproductive), whole males (teneral), male reproductive tracts, female reproductive tracts, male heads (fed, mated), female heads (fed, mated), third instar larva, and pooled female/male salivary glands were examined to assist in addressing core questions of this study. Poly(A)+ RNA was isolated, then measured with the Agilent Bioanalyzer for quality and only those samples with a minimum RIN score of 7 were used to build non-normalized cDNA libraries using a modified version of the Nu-GEN Ovation® RNASeq System V2 (http://www.nugeninc.com). We sequenced each cDNA library (0.125 lane) on an Illumina HiSeq 2000 instrument (~36 Gb per lane) at 100 base pair length. All raw data were made publicly available in the Genbank sequence database, and accession numbers are summarized in Additional File 2, Table S1.

*Automated annotation*

The NCBI Eukaryotic Genome Annotation Pipeline, version 6.4 was used to annotate the *Stomoxys* genome assembly (Stomoxys_calcitrans-1.0.1) with 1.3 billion supporting RNA-seq reads from 8 samples, existing transcripts in the expressed sequence tag (EST) database, RefSeq proteins from *D*. *melanogaster*, *M*. *domestica*, *C*. *capitata*, *B*. *cucurbitae*, and other insects, and a de novo assembled transcriptome …” “The resulting Stomoxys Annotation Release 100 is summarized and available for download at (Trinity-v2; transcriptome shotgun assembly (TSA) sequence database, Accession number GDIM00000000.1).

The resulting *Stomoxys* Annotation Release 100 is summarized and available for download at: <https://www.ncbi.nlm.nih.gov/genome/annotation_euk/Stomoxys_calcitrans/100/>

*Vectorbase, Web Apollo interface, and Community Curation*

Assembly and reference sequence data are also available as part of the Vectorbase platform (<https://www.vectorbase.org/organisms/stomoxys-calcitrans>). A conversion table for Vectorbase to NCBI IDs is available as Table S38.

The Web Apollo tool [3], adopted by VectorBase, was used by our community to review and edit automated gene predictions. Edited gene models were incorporated into a community annotation patch build, ScalU1.4 (release date: February 21, 2018).

*Benchmarking Universal Single-Copy Orthologs (BUSCO) and OrthoFinder analyses*

We used BUSCO to assess quality of the genome assembly and completeness of the predicted gene set. Based on the near-universal single-copy orthologs from dipterans (OrthoDB v8, [4]), 95.1% were found in the assembly and 92.2 in the final predicted gene set.

A species phylogeny based on 664 protein sequences (269,392 amino acids) was reconstructed to determine the evolutionary relationships among nine dipteran species (Fig. S1). The official protein set of *Sarcophaga* *bullata*, *Lucilia* *cuprina*, *Musca* *domestica*, *Glossina* *morsitans*, *Drosophila* *melanogaster*, *Mayetiola* *destructor*, *Aedes* *aegypti*, and *Anopheles* *gambiae* were downloaded from NCBI and searched against the *S*. *calcitrans* gene set using BLASTp. A significant e-value cut-off ≤1e-5 was applied and only genes that had a single hit across all eight species were included in further analysis. A total of 664 individual proteins were aligned with MAFFT [5] using default settings, and alignments were trimmed using gBlocks to remove gaps [6]. The aligned single-copy protein-coding genes were then concatenated and the phylogeny was reconstructed using RAxML version 8.2.8 [7] with the PROTGAMMAWAG model and 100 bootstrap replicates. The phylogeny was visualized with FigTree version 1.4.2 (http://tree.bio.ed.ac.uk/software/figtree/). Orthologous groups of genes were also determined among the nine species using OrthoFinder (v 2.2.7) [8] using default settings.

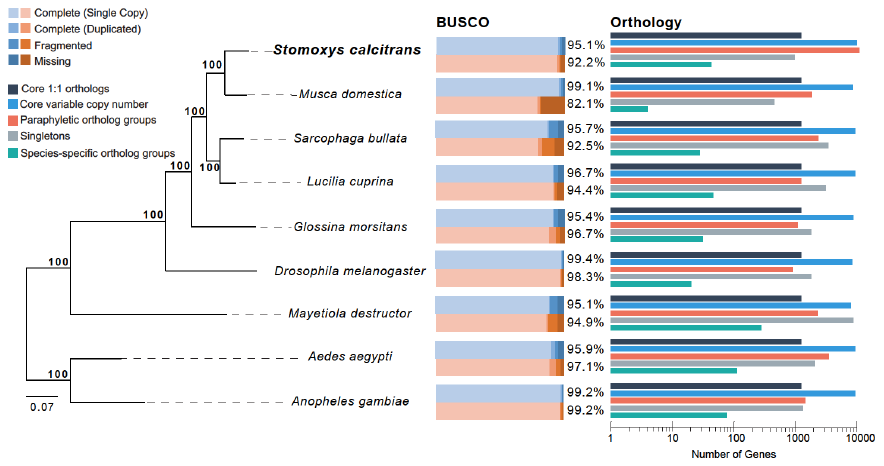

**Figure S1: Phylogenetic placement and genomic comparisons for *Stomoxys calcitrans* and other fly species.** Left, The phylogenetic analysis places *S. calcitrans* as a sister species to the house fly, *Musca domestica*. The phylogeny is built using RAxML and it is based on amino acid sequences from 664 single-copy genes that are present in all nine species. Bootstrap values are shown for every node. Middle, Benchmarking Universal Single-Copy Orthologs (BUSCO, [9] analyses based on the dipteran dataset (odb8). Blue, genome; Red, predicted genesets. Right, Orthology-based analyses of protein coding genes between eight fly species determined the number of genes in single-copy core clusters, variable-copy number core clusters, paraphyletic clusters (non-core, non-species specific), singleton, and species-specific clusters, based on OrthoFinder [8].

**OrthoDB Analysis**

**Contributors:** Evgeny Zdobnov, Panagiotis Ioannidis, Robert Waterhouse

The OrthoDB hierarchical orthology delineation procedure was employed to predict orthologous groups (OGs) of genes across 87 arthropods for OrthoDB v8 [4]. Briefly, protein sequence alignments were assessed to identify all best reciprocal hits (BRHs) between genes from each pair of species, which are then clustered into OGs following a graph-based approach that starts with BRH triangulation. The annotated proteins from the genome of *Stomoxys calcitrans* were first filtered to select one protein-coding transcript per gene and then mapped to OrthoDB v8 at the Diptera (37 species), Endopterygota (72 species), Insecta (80 species), and Arthropoda (87 species) levels. OrthoDB orthology mapping uses the same BRH-based clustering procedure but only allowing proteins from the mapped species to join existing OGs.

**RNA-seq analyses**

**Contributors:** Joshua Benoit, Pia Olafson

The proteome of the stable fly was organized on a hyperlinked spreadsheet (Additional File 2, Table S2) with accompanying information, *e.g.* the presence or absence of signal peptide indicative of secretion, presence of transmembrane domains, and similarities to several databases. Expression values present within this sheet are based on the RNA-seq data sets within Table S1 and analyzed according to previously described methods [10, 11].

RNA-seq analyses were conducted based on methods in Benoit et al. [12] and updated according to Rosendale et al. [13, 14]. RNA-seq datasets analyzed are summarized within Table S1. The main goals for analyzing these datasets were to 1. determine male, female, and larva-enriched gene sets, 2. identify reproductive specific genes, and 3. establish chemosensory genes associated with sexes and specific tissue. Each RNA-seq set was trimmed with CLC Genomics (CLC Bio). Quality of each RNA-seq set was assessed with FastQC prior to analyses. Each datasets was mapped to the predicted gene sets for *S*. *calcitrans* using CLC Genomics with each read requiring at least 95% similarity over 50% of length with three mismatches allowed. Transcripts per million (TPM) was used as a unit of gene expression. Significant enrichment was based on a Baggerly’s test (t-type test statistic) followed by Bonferroni correction at 0.01. Genes were identified by BLASTx searching against an NCBI non-redundant proteins databases for arthropods with an expectation value (e-value) of at least 0.001. Analyses are summarized in Additional File 2, Tables S5 – S13. Genes of interest were compared to the results in Additional File 2, Table S2 for verification before validation by qPCR.

**Validation of RNASeq Data**

**Contributors:** Pia Olafson, Greta Buckmeier

Twenty-five transcripts were randomly selected to evaluate correlation between log_2_ fold changes of quantitative real-time PCR (RT-qPCR) versus RNAseq, as there are no biological replicates of the transcriptome libraries. Total RNAs were isolated from tissues to represent those used for RNASeq, i.e. female or male heads (7d fed, mated), female or male reproductive systems (7d fed, mated, dissected), adult female or male (whole, teneral), adult female or male (whole, 7d fed, mated), third instar larvae. Samples were placed in TriZol, macerated, and stored at -80 °C until isolation using the Zymo Direct-zol**™** method (Zymo Research, Irvine CA) with on-column DNAse treatment (TURBO DNase, ThermoFisher Scientific, Waltham MA). cDNA templates were synthesized from 500ng total RNA in a 20ul volume using SuperScript**^®^** III reverse transcriptase (ThermoScientific) primed with a dTV_n_ oligonucleotide. Primers for RT-qPCR were designed with Beacon software to span exon-intron junctions (Additional File 2, Table S14), and primer efficiencies were estimated using serially diluted cDNAs. Reactions were prepared in a 20ul volume consisting of 250nM each of the forward and reverse primers, cDNA from 25ng RNA, and the iTaq™ Universal SYBR^®^ Green Supermix (Bio-Rad Laboratories, Hercules CA). Reactions were run in triplicate on a LightCycler**^®^** 96 System (Roche Life Sciences, Indianapolis IN). Data were analyzed using the comparative Ct method, and all values were normalized to the reference gene *RpS3*. Pearson’s correlation of log_2_ fold change for RNA-Seq and RT-qPCR results was calculated using GraphPad Prism version 7.00 for MacOSX (GraphPad Software, La Jolla CA).

A significant correlation (Pearson’s *R*^2^ = 0.8643, P<0.0001; Fig. S2) was observed, validating the use of stringent parameters [Baggerly’s test (t-type statistic) followed by a Bonferroni correction at 0.01 (number of genes x α value)] to analyze existing RNASeq data to identify differentially expressed genes.

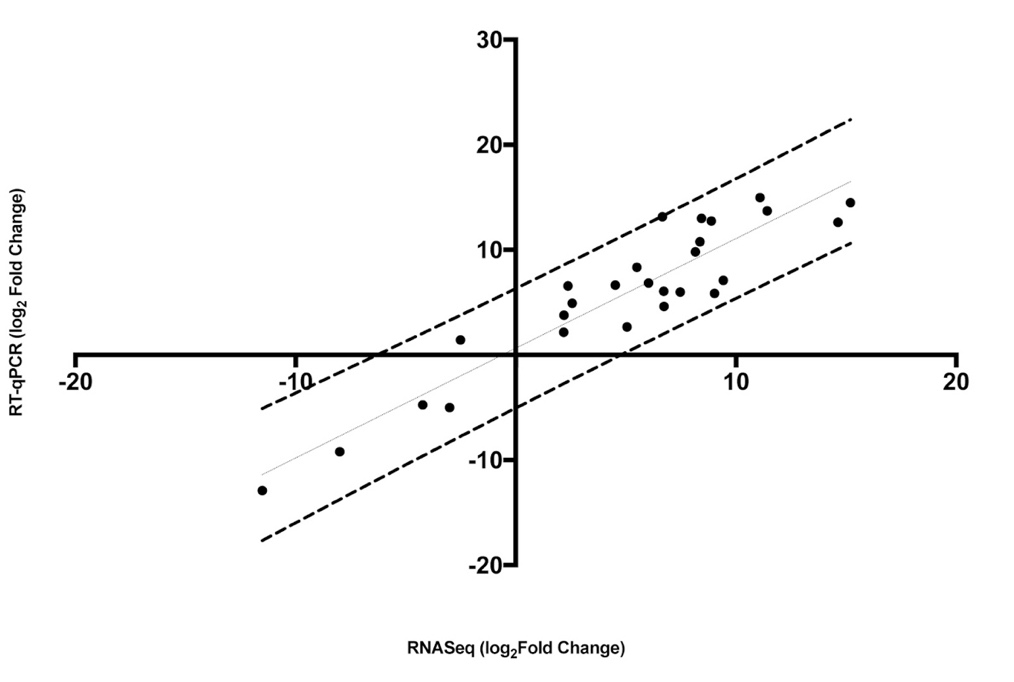

**Figure S2. Pearson's correlation of RNA-seq and RT-qPCR results.** The correlation of the log_2_ fold change analyzed by RNA-Seq (x-axis) with data obtained using RTq-PCR (y-axis). Dashed lines represent the 95% confidence interval.

**Stable Fly Chromosome Arm Assignments**

**Contributor**: Richard P. Meisel (University of Houston)

***Assigning autosomes to Muller elements***

Fly genomes are organized into six chromosome arms, known as Muller elements [15, 16]. Elements A–E correspond to the five gene-rich chromosomes that are autosomal in the most-recent common ancestor, and element F is a gene-poor heterochromatic X chromosome in the common ancestor [17]. The *S. calcitrans* karyotype is missing the heterochromatic element F [18-20], suggesting that it fused to one of the other elements. We have identified the chromosome fused to element F in a companion paper, which also reports the identity of the stable fly sex chromosomes. Here we report the reconciliation of the traditional stable fly chromosome numbering scheme with the Muller element nomenclature. Determining relationships between *S. calcitrans* chromosome numbers and the Muller elements is difficult because of a paucity of mutant markers that have been mapped to *S. calcitrans* autosomes. A mutation causing a black puparium (*bp*) was mapped to *S. calcitrans* chromosome 3 [21]. A similar mutation was mapped to house fly (*M. domestica*) chromosome 1 and sheep blowfly (*Lucilia cuprina*) chromosome 2, both corresponding to Muller element B [22, 23]. It is therefore reasonable to hypothesize that *S. calcitrans* chromosome 3 corresponds to Muller element B. The *S. calcitrans* rolled down wing (*rd*) mutation [24] is phenotypically similar to the house fly *ali-curved* mutation [25, 26], which also maps to house fly chromosome 1 (Muller element B). However, *rd* was mapped to *S. calcitrans* chromosome 4 [24], and not chromosome 3 as expected based on the location of *bp*. The caramine eye (*ca*) and malathion resistance (*malR*) mutations both map to *S. calcitrans* chromosome 2 [24, 27], and they have multiple possible orthologs in other fly species on Muller elements C and E [23, 28]. We therefore conclude that Muller element B either corresponds to *S. calcitrans* chromosome 3 or 4, and *S. calcitrans* chromosome 2 therefore either corresponds to Muller elements C or E.

**Methods**

***Chromosome mapping.*** We used 1:1 orthologs between *D. melanogaster* and *S. calcitrans* to map genes and scaffolds to chromosome arms (Muller elements). This approach works because chromosome arm gene content (synteny) is conserved across higher dipterans [23, 29], and we have previously demonstrated its efficacy with the house fly genome [30].

**Presence and enrichment of repeat elements in *Stomoxys calcitrans* genome**

**Contributors:** T. J. Raszick, S.-H. Sze, C. J. Coates, A. M. Tarone (Texas A&M University)

**Materials and Methods**

A hidden Markov model (HMM)-based approach was used to identify repeat elements in the *S. calcitrans* genome including Class I and II transposons, non-coding RNAs, and satellite sequences (Additional File 2, Table S15). The database of HMMs for 4150 unique repeat elements was obtained from Dfam [31] and evaluated against the *S. calcitrans* genome assembly using nhmmscan in HMMER [32]. Repeat elements identified as having at least one copy in the genome were then further investigated with regards to repeat sequence enrichment for each putative Muller element (Additional File 1, Section 4) using a Bonferroni-corrected hypergeometric test (α = 0.05/1165 = 4.291845e-05). Repeat elements occurring on contigs that were not assigned to any Muller element were also included in this analysis.

For all significantly enriched repeat elements with copy number >1000 found in association with a Muller element, DNA sequences for up to 25 copies were randomly subsampled from each Muller element where those elements occurred. Also, a custom BLAST database was created with the Dfam consensus sequences for those repeat elements. The random subset of sequences was then locally compared to the custom database in a reciprocal BLAST designed to ensure that the results of the HMM-based approach were consistent with the consensus sequences available from Dfam.

**Table S16:** Summary of Dfam repeat elements with >1000 copies including number of unique elements for each family of elements and total number of genomic copies for each family.

| Type of repeat element | Family | Unique elements | Total copies |
| --- | --- | --- | --- |
| Class I (retrotransposons) | LINE | 6 | 19282 |
|  | LTR | 6 | 8223 |
| Class II (DNA transposons) | Cut and paste | 28 | 104621 |
|  | Rolling circle | 8 | 112877 |
| Transfer RNA | Valine | 1 | 5130 |
| Satellite | Undefined *Mus* satellite | 1 | 6236 |

**Results**

The HMM-based search for repeat elements yielded a total of 1165 unique Dfam repeat elements with at least one copy in the *S. calcitrans* genome assembly. Of these, 50 occurred frequently, with >1000 copies genome-wide. The majority of those 50 were Class II transposons, but Class I transposons were also common (Table S16). Cut and paste transposons displayed the highest diversity of unique elements, and rolling circle transposons, while less diverse, occurred with the highest copy number. DNAREP1 was the most common across all types of repeat elements, accounting for 90,273 of the 112,877 occurrences of rolling circle transposons. A subset of 15 unique repeat elements with total genomic copy number >1000 were found to be significantly enriched in association with at least one Muller element (Table S17). Our reciprocal BLAST search reassigned these repeat elements to appropriate consensus sequences with 76.22% success (1010 best hits to the exact same element or to a similar element out of 1325 subsampled sequences).

The Muller D and F elements had distinct patterns of enrichment when contrasted with each other or with Muller A, B, C, and E elements. There were nine unique repeat elements that were significantly enriched in association with one or more of the Muller A, B, C, and E elements, and none of those nine were significantly enriched in association with the D or F. Conversely, there were three unique repeat elements that were significantly enriched in association with the Muller D element and no other Muller element. This pattern suggests that there is some distinct feature of the Muller D element that alters the way it interacts with repeat elements. No repeat elements were significantly enriched in association with the Muller F element. There were three repeat elements that were found to be significantly enriched in association with contigs that were not assigned to any Muller element in the assembly.

**Table S17:** Repeat elements with total genomic copy number >1000 that were found to be significantly enriched in association with at least one Muller element

|  |  |  | Significantly enriched in Muller Element (p < 4.291845e-05) | | | | | |
| --- | --- | --- | --- | --- | --- | --- | --- | --- |
| Repeat Element | **Dfam Taxon**  **Described From** | **Total Genomic Copies** | **A** | **B** | **C** | **D** | **E** | **Unassigned** |
| DNAREP1 | *D. melanogaster* | 90273 | 2.45E-09 | 2.39E-12 |  |  | 2.40E-45 |  |
| Ginger1-N1 | *D. rerio* | 42999 | 1.03E-100 | 1.28E-16 | 9.64E-09 |  | 1.45E-50 |  |
| Baggins1 | *D. melanogaster* | 9191 |  |  |  |  |  | 1.41E-14 |
| HAT1 | *D. rerio* | 7997 | 1.20E-19 |  | 9.83E-05 |  | 3.71E-09 |  |
| Kolobok-2 | *D. rerio* | 6927 | 1.03E-10 |  |  |  |  |  |
| IMPB_01 | *M. musculus* | 6236 | 4.58E-18 |  | 4.31E-05 |  | 1.19E-09 |  |
| HelitronY1 | *C. elegans* | 6096 | 1.49E-13 | 7.66E-61 | 6.61E-60 |  | 1.33E-39 |  |
| HelitronY1A | *C. elegans* | 5147 | 1.44E-12 | 2.37E-45 | 4.19E-51 |  | 9.51E-37 |  |
| Vingi-1 | *C. elegans* | 3988 |  |  |  | 5.09E-06 |  |  |
| HelitronY4 | *C. elegans* | 3987 | 8.85E-11 | 1.62E-31 | 3.16E-36 |  | 2.91E-20 |  |
| Helitron3Na | *M. musculus,*  *H. sapiens* | 2178 |  |  |  |  |  | 6.03E-06 |
| DNA-1-2 | *D. rerio* | 1543 |  |  |  | 9.32E-05 |  |  |
| Dada-tA | *D. rerio* | 1500 |  |  |  | 9.57E-06 |  |  |
| DMCR1A | *D. melanogaster* | 1286 |  |  |  |  |  | 9.85E-07 |
| Helitron1 | *C. elegans* | 1211 |  |  | 5.09E-18 |  |  |  |

***Stomoxys* Reproductive Biology**

**Contributor:** Geoffrey Attardo (University of California, Davis)

**Materials and Methods**

*Yolk protein gene identification and phylogenetic analysis*

Annotated yolk protein gene sequences from *Drosophila melanogaster, Glossina morsitans* and *Musca domestica* were used to identify yolk protein ortholog sequences from the *Musca domestica* and *Stomoxys calcitrans* predicted transcriptomes available at Vectorbase ([www.vectorbase.com](http://www.vectorbase.com)). Gene sequences with significant homology were aligned using the ClustalO software package [33]. The alignment was used to generate a maximum likelihood phylogeny using the tree generation software included in the CLC Main Workbench (Qiagen, Redwood City CA) software package using the following settings (construction method: neighbor joining, Protein substitution model: WAG, Bootstrap analysis: 1000 replicates). Sequences with homology closer to lipase enzyme sequences (the ancestors to yolk protein genes) that form outgroups relative to annotated yolk proteins were removed from the alignment and the phylogeny was recalculated.

*Identification of Male Reproductive Tract Genes and Reciprocal Orthology Analysis*

RNAseq libraries were generated from male and female reproductive tissues from teneral *Stomoxys calcitrans* (see Section 1). RPKM values from the read mapping of these libraries against the putative *Stomoxys* transcriptome from the male and female reproductive tissues were compared to identify genes with a male/female expression ratio of at least 5 and a minimum RPKM value to 50 in the male reproductive tract. The sequences meeting these criteria were extracted from the *Stomoxys* transcriptome using a custom PERL script. Male biased sequences were annotated and associated with GO terms using the BLAST2GO software package [34]. Orthologus sequences for these genes were identified using a  reciprocal hit analysis using the BLAST+ [35] software package against the predicted transcriptomes from *Glossina morsitans*, *Drosophila melanogaster*, *Aedes aegypti* and *Homo sapiens*. Best hits from each of these transcriptomes from these species against male biased *Stomoxys* genes were extracted and used to BLAST back against the predicted *Stomoxys* transcriptome. *Stomoxys* genes with reciprocal hits were detected and annotated as such utilizing a stom PERL script to parse the BLAST output.

***Stomoxys* Chemosensory Gene Families**

**Contributors:** Hugh Robertson (University of Illinois at Urbana-Champaign), Pia Olafson (USDA-ARS)

**Methods.**

TBLASTN searches of the genome assembly were performed with gustatory receptors (GR) from *M. domestica*, *D. melanogaster*, and where relevant the medfly *Ceratitis capitata* [36], and with all newly identified *Stomoxys* GRs. Models were built primarily in the WebApollo server at VectorBase, using a combination of existing automated models, RNAseq information from multiple lifestages, tissues, and sexes, and comparisons with the *Musca* and *Drosophila* Gr gene structures. A few models with regions missing in assembly gaps were repaired using raw genomic and/or RNAseq reads. Pseudogenes were translated as best possible to provide an encoded protein that could be aligned with the intact proteins for phylogenetic analysis. A 200 amino acid minimum was enforced for including pseudogenes in the analysis (roughly half the length of a typical GR), and there are several shorter fragments of genes that were not included in the analysis. All *Stomoxys* GRs were aligned in CLUSTALX v2.1 [37] using default settings with the GRs of *D. melanogaster* [38] and *M. domestica* [39]. Problematic gene models and pseudogenes were refined in light of these alignments. The final alignment was trimmed using TrimAl v1.4 [40] with the “strict” option. Phylogenetic analysis was performed by maximum likelihood analysis using the PHYML v3.0 webserver with default settings [41]. The resultant tree was formatted and colored using FigTree v1.4.2 (http://tree.bio.ed.ac.uk/software/figtree/), while the final version with labels was made in Adobe Illustrator.

Analysis of the Ionotropic Receptor (IR) family in *S. calcitrans* was generally similar to that for the GRs, except that pseudogenes were only included if they encoded at least 50% of the length of a related intact IR (because IRs vary considerably in length, far more than do GRs).

Odorant receptor (OR), odorant binding protein (OBP), and chemosensory protein (CSP) sequences from *Drosophila melanogaster* and *Musca domestica* were used in tBLASTN searches to identify orthologs in the Stomoxys calcitrans 1.0.1 genome assembly. Amino acid sequences from each family were aligned with the MUSCLE algorithm [42], and the alignments trimmed with the trimAl tool using the –strictplus option [40]. The trimmed alignment was used to construct a maximum likelihood phylogeny with the web server version of IQ-TREE software ([43]; best-fit substitution model, branch support assessed with 1000 replicates of UFBoot bootstrap approximation). The OR tree was rooted with the highly conserved ORCO. OBP domains of dimer OBPs were separated for phylogenetic analysis and labeled ‘a’ and ‘b’, and the Plus-C OBPs were not included in assembling alignments for tree construction.

***Stomoxys calcitrans* gustatory receptors**

The Gustatory Receptor family is the more ancient of the two families that make up the insect chemoreceptor superfamily [38, 44-46], and comprises several highly divergent lineages, most involved in taste but some in olfaction [47]. The Odorant Receptor family arose from a GR lineage near to or within a basal insect lineage [48, 49]. In *Drosophila melanogaster* the GR family consists of 60 genes encoding 68 proteins through an unusual form of alternative splicing where multiple long first exons are spliced into a shared single or set of exons encoding the conserved C-terminus of the protein. In *Musca domestica*, the family consists of 80 genes encoding 108 proteins ([39]– 77 genes encoding 101 proteins were reported therein, but as described below, three more genes encoding an additional 7 proteins are recognized here). The family is generally divided into three major and divergent subfamilies. The sugar or sweet receptors, the carbon dioxide receptors, and the bitter taste receptors, but with complications, for example, a lineage within the bitter taste receptor clade has evolved into an important receptor for fructose [50], while others are involved in courtship [51].

The ScalGr gene set consists of 74 models, encoding 113 potential proteins through alternative splicing of eight loci (Supplementary Dataset 1). Ten (9%) of these genes or isoforms are apparent pseudogenes, regions of seven models were repaired using raw genomic and/or RNAseq reads, five models were joined across scaffolds, and eight remain incomplete. Eight genes are modeled as being alternatively spliced and occasionally RNAseq evidence supporting these alternative splices was available. The resultant isoform proteins often differ considerably in most of their sequence, and hence presumably bind different ligands. They are indicated with a lower case letter after the gene name (not to be confused with the lower case letter after the Gr names in Drosophila, e.g. Gr2a, which instead indicates the cytological location of the gene, and hence to avoid confusion alternatively-spliced isoforms in Drosophila are indicated with an upper case letter, e.g. Gr39aC). As a result, the number of apparently intact GR proteins is 103. Less obvious pseudogenes (for example with small in-frame deletions or insertions, crucial amino acid changes, or promoter defects) would not be recognized, so this total might be high.

The GR repertoire was primarily compared with that of *M. domestica* [39], which is in the same family of flies, and *D. melanogaster* [38], as the functions of some of these proteins are at least partially known from the latter. Additional phylogenetic analysis with a more distantly related fly with a complete genome sequence and chemoreceptor analysis, the medfly *C. capitata* [36], was undertaken and is mentioned when relevant, but is not shown. Details of the major subfamilies and gene lineages are below, along with a heat map of normalized expression values for annotated *Gr* transcripts (Fig. S3; Additional File 2, Table S22).

*Stomoxys* has the same set of carbon dioxide receptors as *Musca*, with a duplication of the Gr1 lineage (DmelGr21a) in inverted orientation like *Musca*. These two genes are somewhat awkwardly named Gr1.1 and 1.2, as was done in *Musca*, to maintain a proposed convention that the carbon dioxide receptors be named Gr1-3 [52]. The absence of the Gr2 lineage (also absent from Drosophila) helps confirm that this loss occurred before the Muscidae and Drosophilidae split, but after they separated from the Tephritidae because *Ceratitis* has it.

*Stomoxys* has the same set of eight sugar receptors as *Musca* (Gr4-11). In *Musca* these genes were generally poorly assembled on mostly separate scaffolds, and so both their phylogenetic relationships to the *Drosophila* sugar receptors and their relative locations in the genome were unclear. In *Stomoxys* the situation is somewhat better in that most of these genes are full-length or the assembly could be repaired to be full-length, and Gr7-10 are in an array crossing two scaffolds, however Gr4-6 and 11 are all in separate scaffolds. It therefore remains unclear when these genes became separated as they are in *Drosophila* with the first and last genes (Gr61a and 5a) having moved from the contiguous tandem array in *Ceratitis* that is hypothesized to be the ancestral arrangement in flies [52]. Their phylogenetic relationships are also clarified, with the muscids having duplicated the DmelGr64b lineage (as Gr7/8), while *Drosophila* duplicated DmelGr64c/d. Gr7 is an apparent pseudogene in *Stomoyxs*, with a stop codon near the start codon, confirmed by multiple RNAseq and genomic reads, however as with all such young pseudogenes, this nonsense mutation might be unique to the sequenced strain.

*Stomoxys* has the same duplication of the highly conserved DmelGr43a lineage of fructose receptors as *Musca* (Gr12/13), a lineage independently duplicated to four genes in *Ceratitis* (CcapGr11-14). This lineage clusters with several well-known bitter taste receptors, e.g. DmelGr66a and 33a, and hence presumably evolved from a receptor that originally detected a bitter compound. It certainly did not evolve from within the sugar receptor subfamily.

Most of the remaining *Drosophila* GRs are implicated in perception of bitter tastants or have not yet been functionally characterized [53-66]. The *Stomoxys* representatives were named roughly in the order of the *Musca* genes, however because of gene duplications and losses, this naming system eventually becomes disconnected with orthology. In addition, some of these are rapidly evolving genes, so establishing orthology is not always clearcut, and when feasible microsynteny analysis was employed to assist with these decisions. They are described below in the order of their naming, in paragraphs corresponding to major lineages in the tree, with details of known functions, many of which are relatively new discoveries in *Drosophila*. In particular, the work of Delventhal and Carlson [55] has revealed ligand specificities for some of these bitter taste receptors, showing the breadth of their ability to mediate sensing of diverse bitter compounds.

Gr14 is the conserved ortholog of DmelGr32a, which is a well-known bitter receptor and one of five expressed in all four types of bitter taste neurons on the labellum. It is also involved in courtship through expression in a small set of gustatory receptor neurons on the male foreleg [67], and is implicated in inhibition of male-male courtship [63, 68], as well as mediating rejection of non-conspecific females as targets of male courtship [51], and mediating aggression [69]. DmelGr68a is a relative of Gr32a also involved in courtship [70-72], but it was lost from both *Musca* and *Stomoxys*. Gr15 is a complicated locus with an interesting history in these flies, and is also the next relative of Gr32a/68a and is likely to be involved in courtship [73]. *Ceratitis* has a single gene, Gr20. In *Stomoxys* this locus has two alternatively-spliced isoforms (Gr15a/b). In *Musca* each of these isoforms has been independently duplicated (Gr15a/b and c/d), and remarkably these two isoforms were again independently duplicated in *Drosophila* where they are known as Gr39aA/B and C/D, and the locus has undergone additional duplications in other Drosophila species [74]. DmelGr39aA is another of the five receptors expressed in all four bitter taste neurons in the labellum [73]. Unfortunately when annotating the *Musca* genome, two additional alternatively-spliced loci related to these were missed. They are therefore numbered after the current final *Musca* Gr, so are MdomGr77a-d and 78a/b. In *Stomoxys* they are Gr71a-e and 72 (*Stomoxys* has an additional alternatively-spliced first exon in Gr71, while *Musca* has duplicated the first exon of Gr78). The phylogenetic relationships of these two new genes, which are neighbors in each genome, are unusual. They appear to be confidently related to the DmelGr39aC/D, MdomGr15c/d, and ScalGr15b lineage, implying that they are a duplicate from the terminal alternatively-spliced part of this locus. This relationship is also supported in smaller trees using only the proteins described in this paragraph. *Ceratitis* does not have these genes. It appears then that *Stomoxys* has a slightly smaller repertoire of these GRs involved in the social behaviors of courtship and aggression than does *Musca*, but both have a set of GRs absent from *Drosophila*, while having lost the DmelGr68a ortholog.

ScalGr16-18 are together as neighbors in one scaffold, as is the case for their *Musca* relatives. *Drosophila* lost the ortholog of Gr17 (which is a frameshifted pseudogene in *Stomoxys*), but Gr16 is the ortholog of DmelGr2a while Gr18 is related to DmelGr23aA/B. Both of these genes in *Drosophila* are expressed in larval pharyngeal organs [54] and the adult labral sense organ and Gr2a is required for avoidance of high salt concentrations in food [75]. Delventhal and Carlson [55] find that Gr2a responds to several plant bitter compounds including the alkaloids caffeine, lobeline, and theophylline, as well as the quite dissimilar phenylpropanoid umbelliferone. Like DmelGr23aA/B, the Gr18 locus is also alternatively-spliced in *Stomoxys*, however this is an ancient duplication of the first exon, independent of the DmelGr23aA/B duplication, and *Musca* has lost the second isoform from its version of the Gr18 locus, so only has a single isoform related to ScalGr18a. Clustering with Gr17/18 in the tree is MdomGr19, which is the highly diverged ortholog of DmelGr39b (supported by microsynteny analysis as well). But *Stomoxys*, despite maintaining microsynteny of the flanking genes, has lost this gene. MdomGr20/21 are thought to be duplicates distantly related to DmelGr98a, although they do not cluster in the tree but do when CcapGr37 is included in the analysis, but this gene lineage was also lost from *Stomoxys*. *Stomoxys* has a single ortholog of DmelGr8a (ScalGr19), a receptor for a plant-derived insecticide L-canavanine [57, 76], but this lineage expanded to 7 genes in *Musca* (MdomGr22-28). Related to this lineage is the alternatively-spliced gene MdomGr29a-c, which has a relative in *Ceratitis* (CcapGr33), but was independently lost from *Drosophila* and *Stomoxys*. DmelGr98b-d are one of the few examples of a small tandem array of GR genes in *Drosophila*, and are related to an independent expansion of six genes, some in an array, in *Musca* (MdomGr30-35). *Stomoxys* has only two genes in this lineage (ScalGr20 and 21) quite separate in two large scaffolds, and their phylogenetic relationships suggest that they are separately related to the expansions of MdomGr30-32 and 33-35, respectively, and that some of these duplications occurred early in the *Muscid* lineage and were lost from *Stomoxys*. DmelGr98b is also involved in perception of L-canavanine [76], so this set of muscid expansions is likely involved in perception of similar bitter compounds. There is a single ortholog for DmelGr9a in *Musca* (MdomGr41), but it is an alternatively-spliced gene in *Stomoxys* (ScalGr27a/b) (this relationship is not reflected in the tree, but is found in larger trees including *Ceratitis*, and microsynteny analysis supports their orthology). Finally, at the base of the cluster of the above GR lineages in the tree are MdomGr76 and ScalGr70, a lineage with a single conserved ortholog in *Ceratitis* (CcapGr70), but which was lost from *Drosophila*. This phylogenetic cluster of bitter taste receptors has been extensively studied in *Drosophila*, where they are variously expressed in the labellum, abdominal neurons, and forelegs of adult flies, as well as in larvae and several are known to mediate perception of important plant bitter compounds. *Stomoxys* has just 10 proteins in this phylogenetic cluster, the same number as *Drosophila*, compared with 24 in *Musca*, resulting from gene losses in *Stomoxys* and duplications in *Musca*, suggesting that *Musca* has a greater chemical ecological need for this clade of bitter receptors.

ScalGr22 is the ortholog of MdomGr36 and DmelGr66a, one of the best-known bitter taste receptors and another of the five expressed in all four classes of bitter taste neurons on the labium. Like *Musca* (MdomGr37), *Stomoxys* has a paralog (ScalGr23) that is absent from *Drosophila*, so presumably this MdomGr37/ScalGr23 lineage, which is also present in *Ceratitis* (CcapGr39), is another ancient bitter receptor that *Drosophila* lost. ScalGr24 is the ortholog of MdomGr38 and DmelGr33a, another well-known bitter receptor and another of the five expressed in all four types of bitter taste neurons on the labium. Related to it is a complicated alternatively-spliced gene, ScalGr25a-h, MdomGr39a-g, and DmelGr28bA-E. In *Drosophila* the isoforms from this gene are involved in bitter taste as well as being expressed in other neurons [55, 77], and some have been implicated in sensing light and temperature, e.g. [78-80]. The muscid flies have orthologous isoforms for each of the five *Drosophila* isoforms, but some have been additionally duplicated in the muscids. All are highly conserved in sequence, and may well perform similar roles in muscids as in *Drosophila*. Immediately downstream of this complex gene in each genome is the similarly well-conserved receptor, ScalGr26, MdomGr40, and DmelGr28a, which has similar broad expression in both gustatory and other neurons [77] and responds to a wide variety of bitter compounds [55]. Note again that the fructose receptor DmelGr43a and its muscid orthologs cluster with these bitter receptors, having presumably changed its ligand from a bitter tastant to fructose at some point.

ScalGr28 is the ortholog of MdomGr42 and DmelGr10a, another well-known bitter receptor that has a remarkably similar response profile to a quite of bitter compounds as Gr2a, despite being phylogenetically far removed from it [55]. In *Stomoxys* this gene is at the start of a large and complicated array of genes spanning two large scaffolds, with additional related genes on four other large scaffolds implying that they moved from the original array of genes, for a total of 48 proteins (ScalGr29-57) including alternative splicing of Gr47a-t. These form a major expansion of candidate bitter taste receptors in *Stomoxys*, comparable to a similarly complicated set in *Musca* (MdomGr43-64, encoding 35 proteins), and together these are a major expansion compared with *Drosophila* and *Ceratitis*. In *Ceratitis* and *Musca* these genes are all adjacent to each other in a single scaffold, which indicates how an original expansion in a fly ancestor has become hugely expanded in the muscids, but not *Drosophila*, while in the latter the genes have been split up on different chromosome arms. The presence of subsets of these genes in four other large scaffolds in *Stomoxys* (Gr48/49, 50, 51-54, and 55-57) implies that some genomic shuffling has occurred on this fly lineage as well. The tree reveals that these genes form three major expanded clades in the muscids. CladeA is related to DmelGr59a/b and consists of independent expansions in the two muscids, with two genes encoding five proteins in *Musca* versus 23 in *Stomoxys*. Clade B has four genes in *Stomoxys* and 16 in *Musca*, all of which are *Musca*-specific duplicates. The relationship of this clade to the *Drosophila* GRs in the tree is unclear. Clade C is related to DmelGr36a-c and Gr59c/d and consists of a simple gene (ScalGr46 and MdomGr51) adjacent to a large alternatively-spliced gene in each species (ScalGr47a-t and MdomGr52a-k). Most of the duplications of these first exons occurred within either *Musca* or *Stomoxys*, presumably from a much smaller ancestral alternatively-spliced locus. In *Stomoxys* this large gene spans two large scaffolds and one contig. In *Drosophila*, Gr59c responds to the plant alkaloids berberine and lobelline, while Gr36a responds to a wide variety of bitter compounds including the alkaloid sparteine, the terpenoid saponin, and the phenanthrene aristocholic acid [55]. Finally, one member of this expansion in *Musca* (MdomGr43) is so divergent it does not cluster with the others, and similarly several *Drosophila* GRs that might be related to these expansions do not cluster confidently with them, e.g. DmelGr10b, 47a, and 85a. In larger analyses including the *Ceratitis* GRs, DmelGr10b clusters with MdomGr43, while DmelGr47a and 85a cluster with Clade B. DmelGr47a is a narrowly-specific receptor for the plant bitter compound strychnine [59]. Although neither of the muscids appears to have a convincing close relative for the recently-duplicated DmelGr22a-f genes in the tree, *Ceratitis* has six related and also recently-duplicated genes, and in larger trees including *Ceratitis*, ScalGr60a/b and MdomGr67 cluster with this lineage. In *Drosophila*, Gr22b responds to several bitter compounds from plants like the glycoside cucurbitacin, the terpenoid azadirachtin, and the alkaloids lobeline and berberine [55]. In summary, this large grouping of genes, originating from a small cluster in a fly ancestor and now distributed across the *Drosophila* genome, has been expanded in a major fashion in the muscids, and especially in *Stomoxys*, with a total of 49 proteins versus 36 in *Musca*, 17 in *Drosophila*, and 12 in *Ceratitis*.

ScalGr58 and 59 are simple orthologs of MdomGr65 and 66 and DmelGr47b and 57a, respectively, forming a divergent lineage in the tree. In Drosophila Gr57a is expressed along with 2a, 23a, and 93d in the larval pharyngeal organ [54] and the adult labral sense organ [75], but nothing is known about their ligand specificities.

ScalGr60-62 are together in one scaffold and related to MdomGr67-69 and DmelGr58a-c, however two of them are alternatively-spliced encoding nine proteins and constituting yet another Stomoxys-specific expansion of candidate bitter taste receptors. As noted above, ScalGr60a/b and MdomGr67 might be related to DmelGr22a-f. ScalGr61 and the alternatively-spliced Gr62a-f are related to MdomGr68 and DmelGr58a/b in another expanded Clade D, while *Stomoxys* appears to have lost the ortholog of MdomGr69/DmelGr58c. Gr58c was included in the study by Delventhal and Carlson [55] and responds to an overlapping set of bitter compounds as Gr22b, but also to additional plant bitter compounds such as the flavonoid myricetin, the benzopyrone coumarin, and the alkaloids strychnine and quinine. Presumably then this is yet another expansion of bitter receptors in *Stomoxys*.

ScalGr63-65 are together in a scaffold and related to MdomGr70/71 and DmelGr59e/f in another distinctive gene lineage, but little is known about Gr59e/f beyond expression in the larval sense organs [56].

ScalGr66-69 are rapidly evolving orthologs of several more *Musca* and *Drosophila* Grs, all clustering weakly phylogenetically with the large expansions described above. ScalGr66 is the ortholog of MdomGr72 and DmelGr77a, ScalGr67 is the ortholog of MdomGr73 and DmelGr89a (the fifth Gr expressed in all four bitter neuron types), ScalGr68 is the ortholog of MdomGr74 and DmelGr93a, which is required for response to caffeine [58], while the alternatively-spliced ScalGr69a/b is the ortholog of MdomGr75a/b and DmelGr94a and related to Gr97a, which along with Gr33a and 66a is involved in quinine perception [53]. This cluster includes a previously unrecognized gene in *Musca*, which is named MdomGr79, with ortholog ScalGr73, related to DmelGr92a and 93b-d, the last of which genes is expressed along with 2a, 23a, and 57a in the larval pharyngeal organ [54] and the adult labral sense organ [75].

In summary, while the carbon dioxide, sugar, and fructose receptors are relatively well conserved in these two muscids, as is the case for many other insects, the bitter taste receptors reveal considerable gene family evolution both with respect to the available relatives of these muscid flies, *Drosophila* and *Ceratitis*, and between these two muscids. *Stomoxys* has lost or not expanded several lineages implicated in courtship, but expanded several others. The ligands for some of these *Drosophila* bitter receptors are being elucidated [55] and provide an early glimpse into what these expansions might mean for the sensory capabilities of these muscids, however it will require determination of the ligand specificities of these muscid receptors to fully understand the ecological significance of the differential expansions and contractions of their bitter taste abilities.

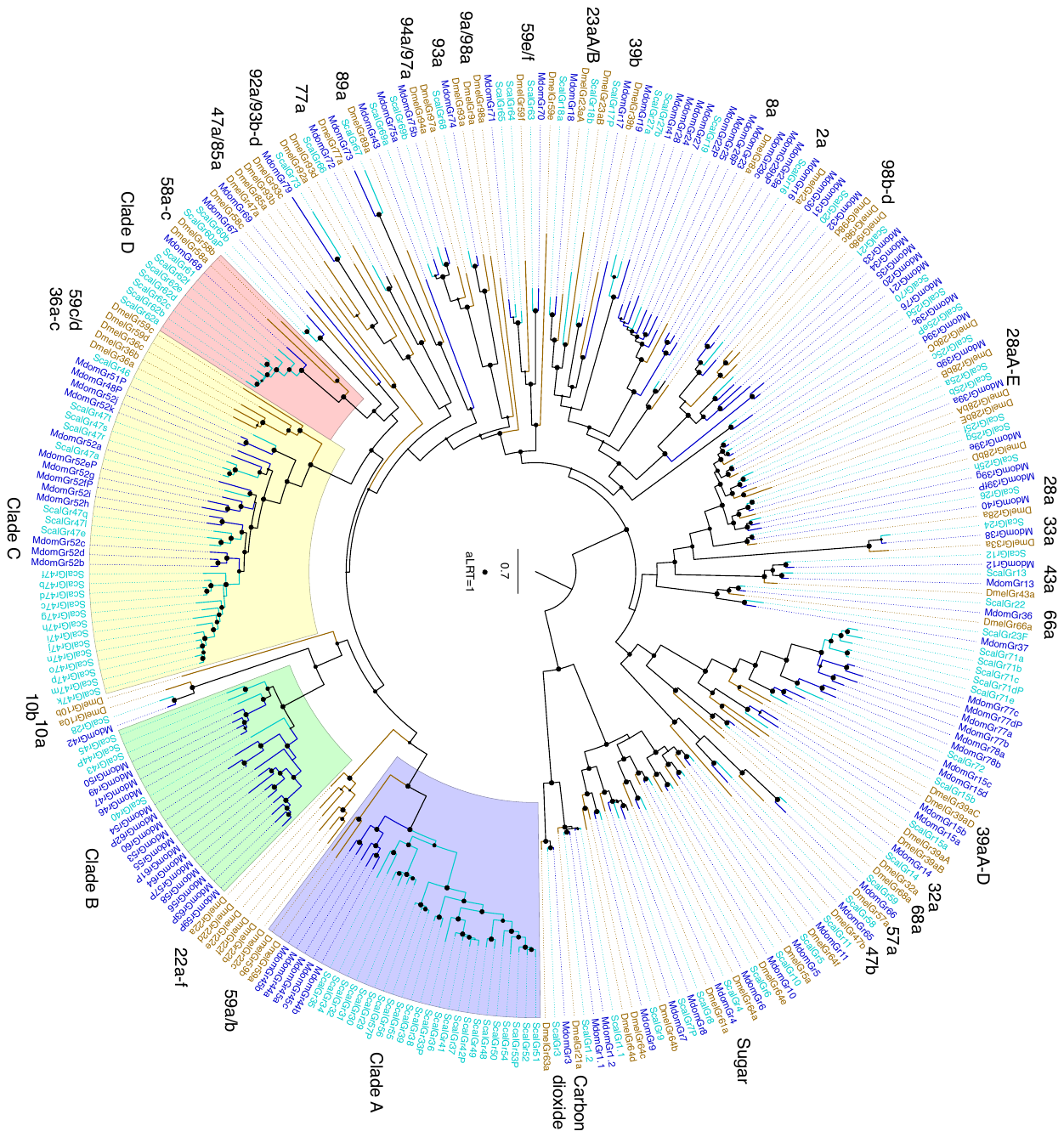

**A**

**Figure S3 (Fig. 5, main paper). *Stomoxys* Gustatory Receptor Gene Family A.** Phylogenetic tree of the *Stomoxys calcitrans* GRs with those of *Drosophila melanogaster* and *Musca domestica.* This is a maximum likelihood tree rooted by declaring the distantly-related and divergent carbon dioxide and sugar receptor subfamilies as the outgroup. The *S. calcitrans* and *M.* *domestica* gene/protein names are highlighted in blue and teal, respectively, while *D. melanogaster* names are in mustard. Support levels from the approximate Likelihood-Ratio Test (aLRT) from PhyML v3.0 are shown on branches. Subfamilies and individual or clustered *Drosophila* genes are indicated outside the circle to facilitate finding them in the tree. Four clades of candidate bitter receptors that are expanded in the muscids are highlighted. Pseudogenic sequences are indicated with the suffix P. **B.** Heat map of normalized expression values for *Gr* transcripts annotated from the *Stomoxys* genome.

**Figure S3 (contd).**

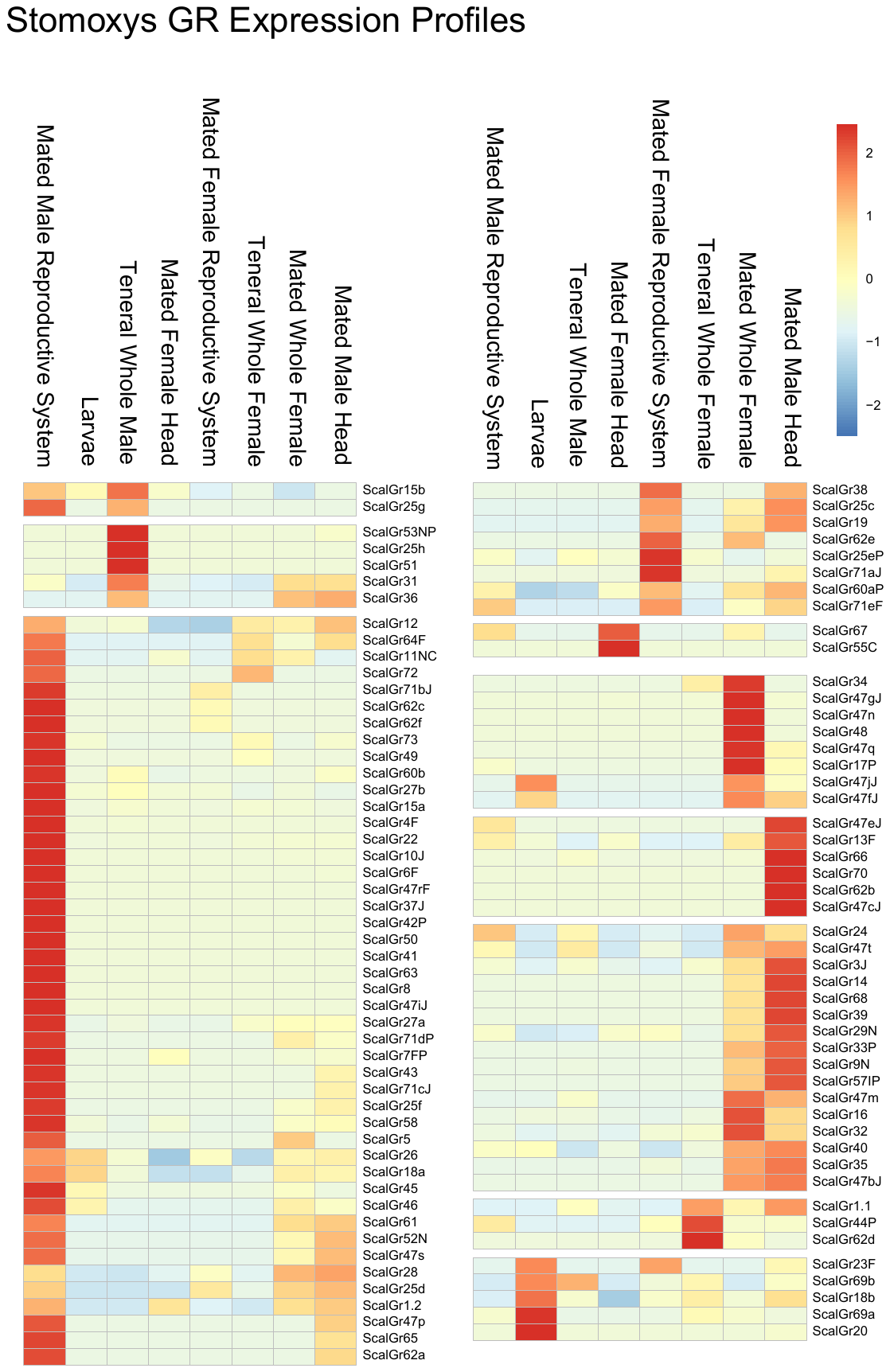

**B.**

**The Ionotropic Receptor (IR) Gene Family**

The Ionotropic Receptor family is a variant lineage of the ancient ionotropic glutamate receptor family [44, 81-83]. These proteins have three transmembrane domains. Like the GRs they are involved in both olfaction and gustation, as well as sensing light, temperature, and humidity [83]. In *D. melanogaster* the family consists of 60 intact genes and four long pseudogenes. *M. domestica* has a considerable expansion to 100 intact genes and nine pseudogenes [39].

The IR family consists of 131 intact genes and 14 pseudogenes for a total of 145 models (Supplementary Dataset 2), although nine remain partial with termini missing in gaps, while three were repaired and one was joined across scaffolds. The IRs were named using the same convention as employed for *Musca*, *Ceratitis*, and several other arthropods, with those showing clearcut orthology with *Drosophila* IRs named for their *Drosophila* orthologs, after which genes were named in a series from Ir101, to avoid any confusion with the *Drosophila* IRs, which only go up to Ir100a, having been named for their cytological locations. Details of the major subfamilies and gene lineages are below.

In *Drosophila* and most other insects examined to date, there are two IRs that are highly conserved both in sequence and length and in being phylogenetically most closely related to the ionotropic glutamate receptors from which this variant ionotropic receptor family of chemoreceptors evolved [81, 82]. These are Ir8a and 25a, both of which function as co-receptors with other IRs [83]. While *Stomoxys* has the expected single conserved ortholog of Ir8a, surprisingly it has four paralogs of Ir25a, named Ir25a1-4. The first exon is missing from the assembly for two of these, apparently in gaps, but otherwise they all appear intact and all but Ir25a2 have extensive RNAseq support. They therefore appear to be functional genes/proteins, but their functions are enigmatic as such duplications of Ir25a are rarely observed in other insects.

Most of the remaining conserved IRs with convincing orthologous relationships are indicated with their *Drosophila* names in Fig. S4. For the most part these are simple 1:1 orthologous relationships, and include proteins such as Ir76b, a third apparent co-receptor involved in sensing salt, amines, and amino acids [84-87], Ir41a implicated in perception of polyamines [86], Ir64a implicated in perception of acidic odors [88, 89], Ir84a which has phenylacetic acid and phenylacetaldehyde as ligands [90], Ir75a that mediates perception of acetic acid [Prieto-Godino, 2017 #115][91], Ir75b/c that respond to butyric and proprionic acid in *D. melanogaster* and hexanoic acid in *D. sechellia* [92], and Ir21a, 40a, 68a, and 93a implicated in temperature and humidity sensing [93-96]. Several instances of duplications in *Stomoxys* are evident, specifically Ir10a1/2, Ir41a1/2, and Ir75d1/2. The latter two lineages are commonly expanded in other insects, and in *Drosophila* are involved in sensing various acids and amines [86, 91, 94]. In *Musca*, in contrast, there are five duplicates of Ir10a (two are pseudogenes) and three duplicates of Ir76a, but the ligands are unknown for these two genes in *Drosophila*.

The Ir7a-g and 11a genes in *Drosophila* are expressed in larval and adult gustatory organs [94], but ligands for these receptors are unknown. This subfamily is considerably expanded in the muscids, and given the complexities of the relationships, in both *Musca* and *Stomoxys* they are not named for their *Drosophila* relatives, but rather begin the numbered series from Ir101, in the case of *Stomoxys* to Ir121 and in *Musca* to Ir126 (Fig. S4). These IR gene expansions strongly suggest an expanded gustatory capacity.

Finally, a large clade of “divergent” IRs in *Drosophila* is involved in gustation and is known as the Ir20a subfamily of 33 proteins (including the four pseudogenic ones) [97, 98]. This clade of mostly intronless genes is considerably expanded in *Musca* to 53 members (MdomIr127-179), and even more so in *Stomoxys* to 96 members (ScalIr122-217). This subfamily consists mostly of minor expansions in *Drosophila* and major expansions in the two muscids, labeled clades A-G in the tree (Fig. S4). Although support for these clades is low, they do appear to represent seven independent expansions of lineages that are represented by 0-6 genes in *Drosophila*. The largest of these is clade G consisting of 16 *Musca* genes (MdomIr163-178) and 50 genes in *Stomoxys* (ScalIr153-202), apparently related to DmelIr47a/b, 67a, and 94a-c. Also noteworthy is clade B, which has just one *Musca* gene (MdomIr149), but 15 *Stomoxys* genes (ScalGr203-217), apparently related to DmelIr60e and 67b/c. Putative ligands in *Drosophila* are limited to carbonation sensing and specific carbohydrates [99, 100], but the expansion in *Stomoxys* warrants study as feeding preference on carbohydrates is low. Expression pattern revealed one specific IR (ScalIR119) that is highly enriched in the male RS compared to all other stages (Fig. S4b; Additional File 2, Table S22). Of interest, the ortholog for this IR in *Drosophila* (Ir7a) has increased expression associated with both male and female reproductive organs [101], suggesting a potential critical role in fly reproduction. This Ir20a subfamily also has most of the pseudogenes, although the Ir7 subfamily also has some, indicating that these two subfamilies, in addition to numerous young duplicates in the muscids, also exhibits the most gene losses, and hence are the most evolutionarily dynamic parts of the IR family. The expansions of this and the Ir7 subfamily in the muscids, and particularly *Stomoxys*, mirror those of bitter taste receptors in the GR family.

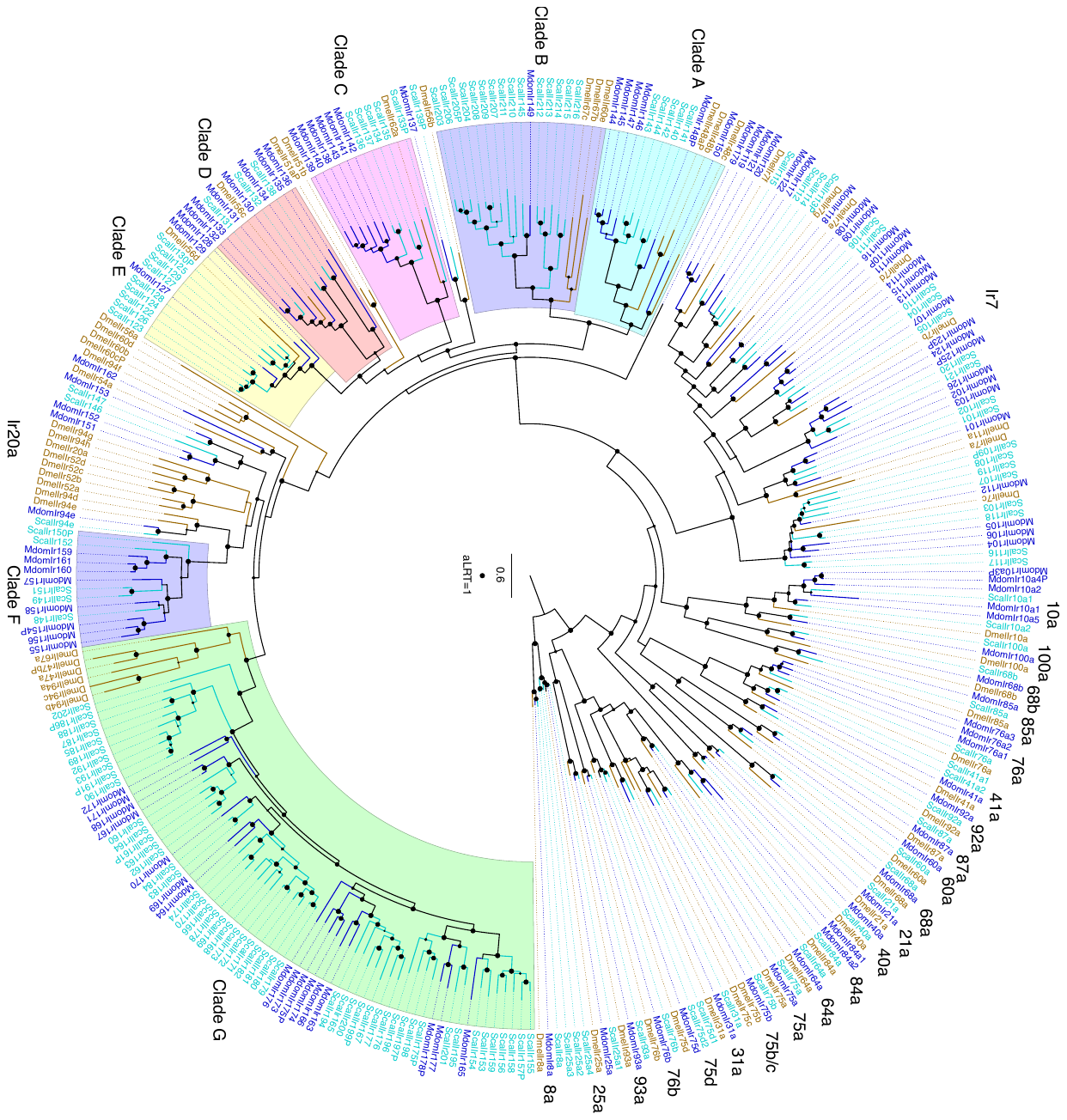

**A**

**Figure S4 (Fig. 6, main paper). Stomoxys Ionotropic Receptor Gene Family A.** Phylogenetic tree of the *Stomoxys calcitrans* IRs with those of *Drosophila melanogaster* and *Musca domestica.* This is a maximum likelihood tree rooted by declaring the Ir8a/25a lineage as the outgroup. The *S. calcitrans* and *M.* *domestica* gene/protein names are highlighted in blue and teal, respectively, while *D. melanogaster* names are in mustard. Support levels from the approximate Likelihood-Ratio Test from PhyML v3.0 are shown on branches. Subfamilies, clades, and individual *Drosophila* genes are indicated outside the circle to facilitate finding them in the tree. Pseudogenic sequences are indicated with the suffix P. **B.** Heat map of normalized expression values for *Ir* transcripts annotated from the *Stomoxys* genome.

**Figure S4 (contd.)**

**B**

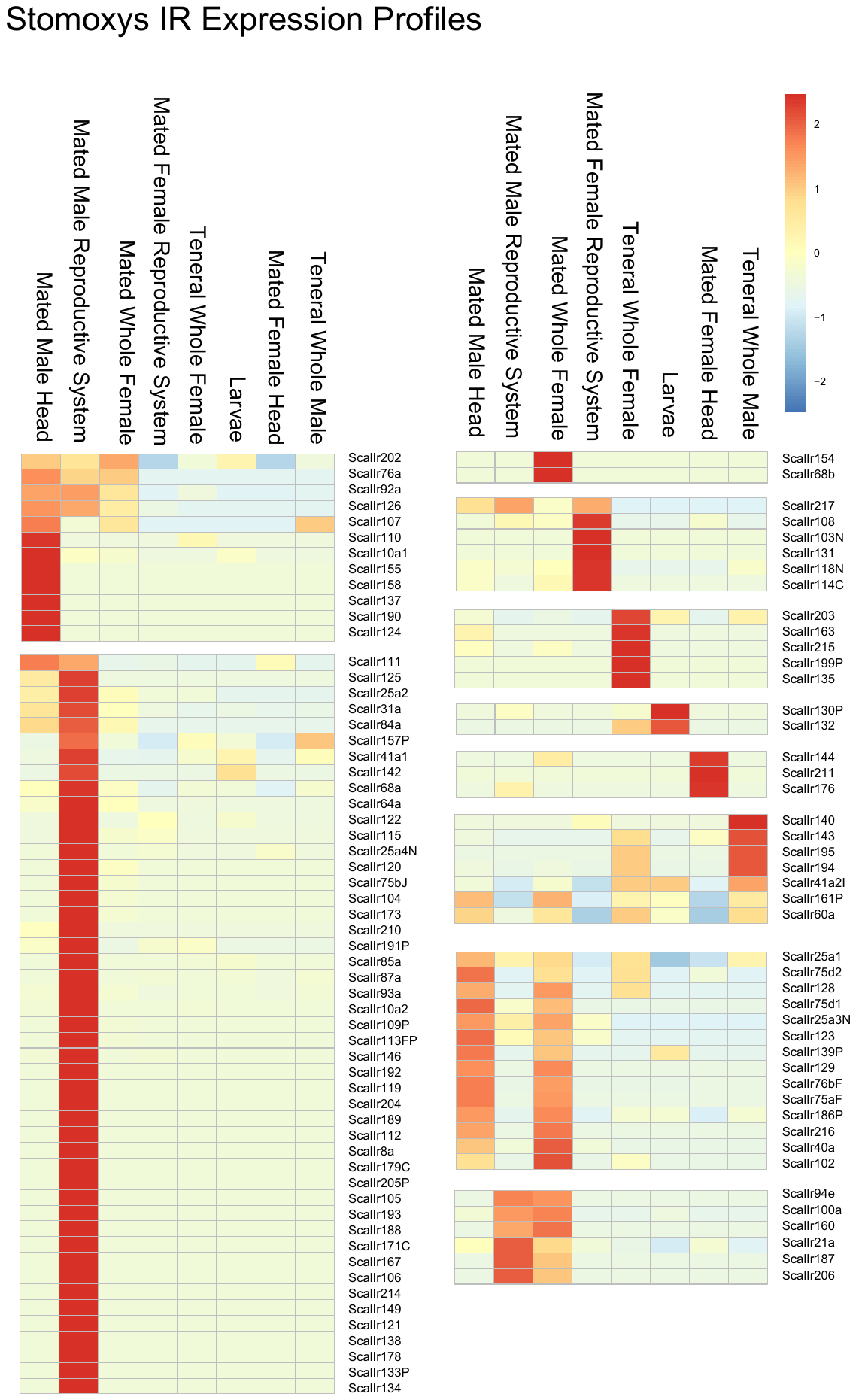

***Stomoxys* Odorant Binding Protein (OBP) Gene Family**

A tBLASTN search of the *Stomoxys* genome using annotated odorant binding protein (OBP) sequences from *D. melanogaster* (38 OBPs, 12 “Plus C” OBPs) and *M. domestica* (87 OBPs, 6 “Plus C” OBPs) resulted in 90 ScalObp gene models (Figure S5; Supplementary Dataset 3). Nine are new models constructed during manual annotation, not including ones resulting from fixing or splitting a model or joining across scaffolds (Additional File 2, Table S19). The 90 gene models include five “Plus-C” OBPs (*ScalObp86-90*), three “Minus-C” (*ScalObp61-63*), and two dimer OBPs (*ScalObp52* and *53*). Classical odorant binding proteins are typically small and encode six cysteines in conserved positions that form three disulfide bonds to stabilize the molecule. Variations in the number of cysteine residues in conserved positions of the OBP-encoding domain were grouped into “Plus-C” and “Minus-C” OBPs comprised of eight cysteine residues and four or five cysteine residues, respectively [102]. Evidence for duplication and subsequent fusion of classical OBPs are represented by dimer OBPs [102]. The *Stomoxys* “Minus-C” Obps are organized in tandem and are expanded relative to the Drosophila “Minus- C” *DmelObp99c* and the single Musca ortholog *MdomObp60*. The two *Stomoxys* dimer OBPs are orthologous to dimers from *Musca* and *Drosophila* MdomObp53/DmelObp83cd and MdomObp54/DmelObp83ef, although no orthologs to *Musca* dimers MdomObp30 and 34 were identified.

More than half the gene models are organized as tandem clusters across three scaffolds, which is consistent with OBP gene organization in other dipteran genomes [103]. Scaffold KQ080538 contains 13 gene models (Fig. S5, yellow shade; *ScalObp11-23*) that are members of a *Stomoxys* gene expansion of 7 genes related to *DmelObp56b*, 4 genes related to *DmelObp56a*, and 2 genes related to *DmelObp56d,e*. Eighteen gene models are present on scaffold KQ080743, all of which are members of a *Stomoxys* gene expansion related to *DmelObp56h* (Fig. S5, orange shade; *ScalObp24-43*) that includes 9 *Musca* OBPs (*MdomObp30-38*). Two additional *Stomoxys* OBPs that are part of this expansion, *ScalObp25* and *ScalObp34*, are located on small scaffolds where they are the only gene models and are likely part of the same chromosome. DmelObp56h has a role in male mating behavior, as gene silencing results in distinct changes in the cuticular hydrocarbon profile of male Drosophila and in reduction of 5-T, a hydrocarbon that is produced by males and thought to delay onset of courtship [104]. The expression pattern of *Stomoxys* transcripts in this expansion is diverse, with several robustly expressed in male/female heads (*ScalObp24*, *25, 30*, and *32*) while others have RNASeq reads only in larvae (*ScalObp31*) or male reproductive system (*ScalObp35*). Lastly, KQ079977 contains 20 gene models (Fig. S7, orange branches) comprising a muscid-specific expansion of 14 *Stomoxys* OBPs (*ScalObp64-77*) and 15 *Musca* OBPs, as well as six gene models (*ScalObp56-63*) related to *DmelObp99a-d*. *ScalObp58* and *70* cluster on this lineage, but are located on small, separate scaffolds where they are the only gene model present suggesting these are part of the same chromosome. There are at least 13 1:1:1 OBP orthologs shared between *Stomoxys*, *Musca*, and *Drosophila*. These include: ScalObp46/DmelObp69aA/MdomObp47; ScalObp3/DmelObp19b/MdomObp2; ScalObp4/DmelObp19c/MdomObp3; ScalObp5/DmelObp19d/MdomObp4; ScalObp54/DmelObp84a/MdomObp55; dimer ScalObp53/DmelObp83e-f/MdomObp54; dimer ScalObp52/DmelObp83c-d/MdomObp53; ScalObp51/DmelObp83g/MdomObp52; ScalObp10/DmelObp44a/MdomObp15; ScalObp60/DmelObp99b/MdomObp59; ScalObp58/DmelObp99d/MdomObp58.

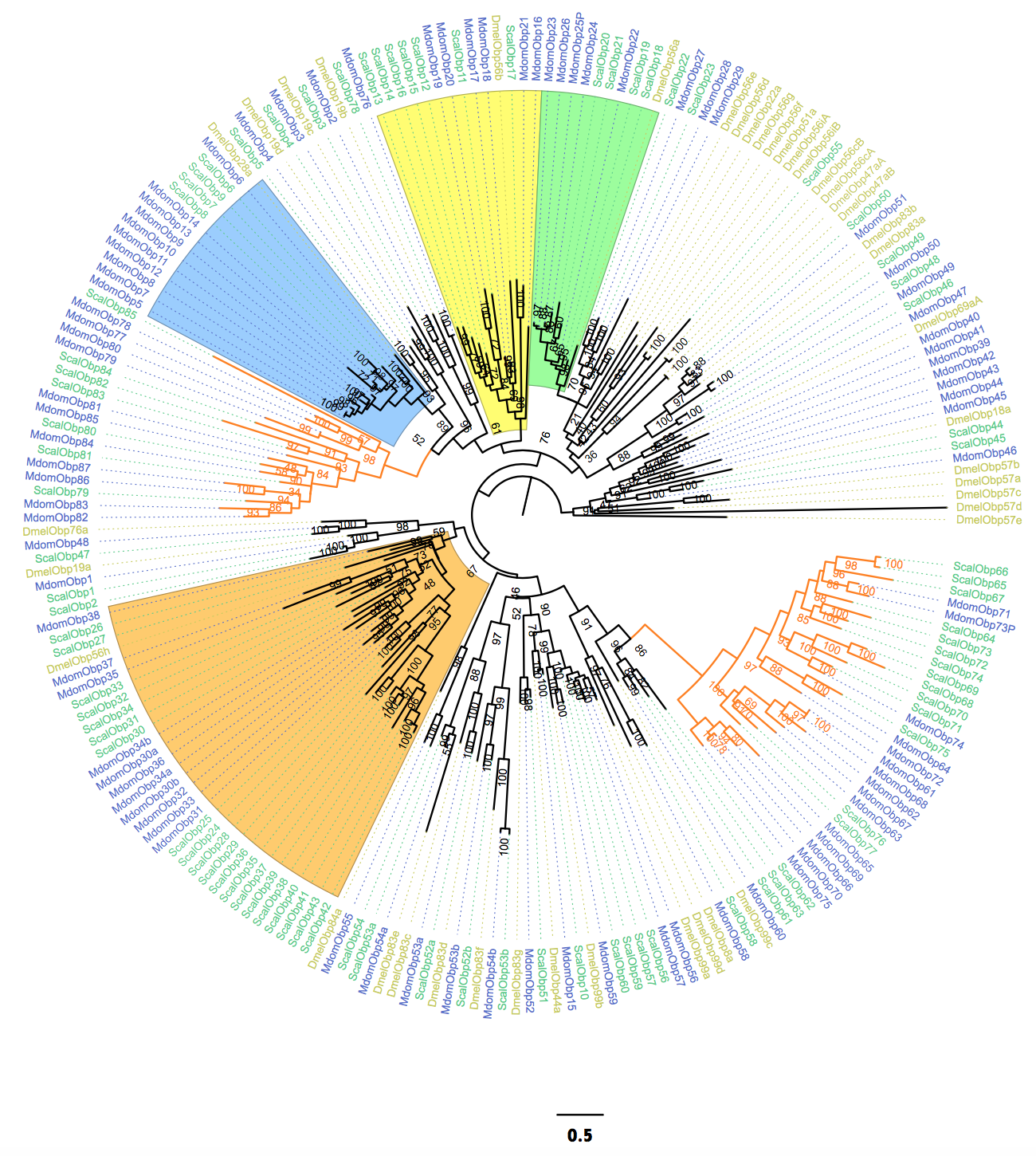

**A**

**Figure S5 (Fig. 4, main paper). Stomoxys Odorant Binding Protein Gene Family. A.** Phylogenetic tree of the *Stomoxys calcitrans* OBPs with those of *Drosophila melanogaster* and *Musca domestica.* The *S. calcitrans* and *M.* *domestica* gene/protein names are highlighted in teal and blue, respectively, while *D. melanogaster* names are in mustard. Maximum likelihood phylogeny was constructed using the web server version of IQ-TREE software (Trifinopoulos et al., 2016; best-fit substitution model, branch support assessed with 1000 replicates of UFBoot bootstrap approximation). **B.** Heat map of normalized expression values for *Obp* transcripts annotated from the *Stomoxys* genome.

ScalObp50/MdomObp51 is an OS-E-like protein, as it clusters with DmelObp83b (OS-E), which is co-expressed along with DmelObp83a (OS-F) and a third OBP (DmelObp76a) in a subset of *Drosophila* sensilla (trichoid). ScalObp48/MdomObp49 and ScalObp49/MdomObp50 are arranged in tandem on scaffold KQ081519, and they encode OS-E/-F-like proteins. Smaller trees constructed using related OS-E/-F proteins from a variety of *Drosophila* species supported an absence of an *OS-F* encoding gene in *Stomoxys*; rather, ScalObp49/MdomObp50 appears to be related to the newly identified OS-X proteins (Fig. S6; [105]). All three *Stomoxys* sequences have conserved intron/exon boundary locations and ScalObp50, while arranged at the terminal end of a separate scaffold (KQ081571), is likely part of the same chromosome.

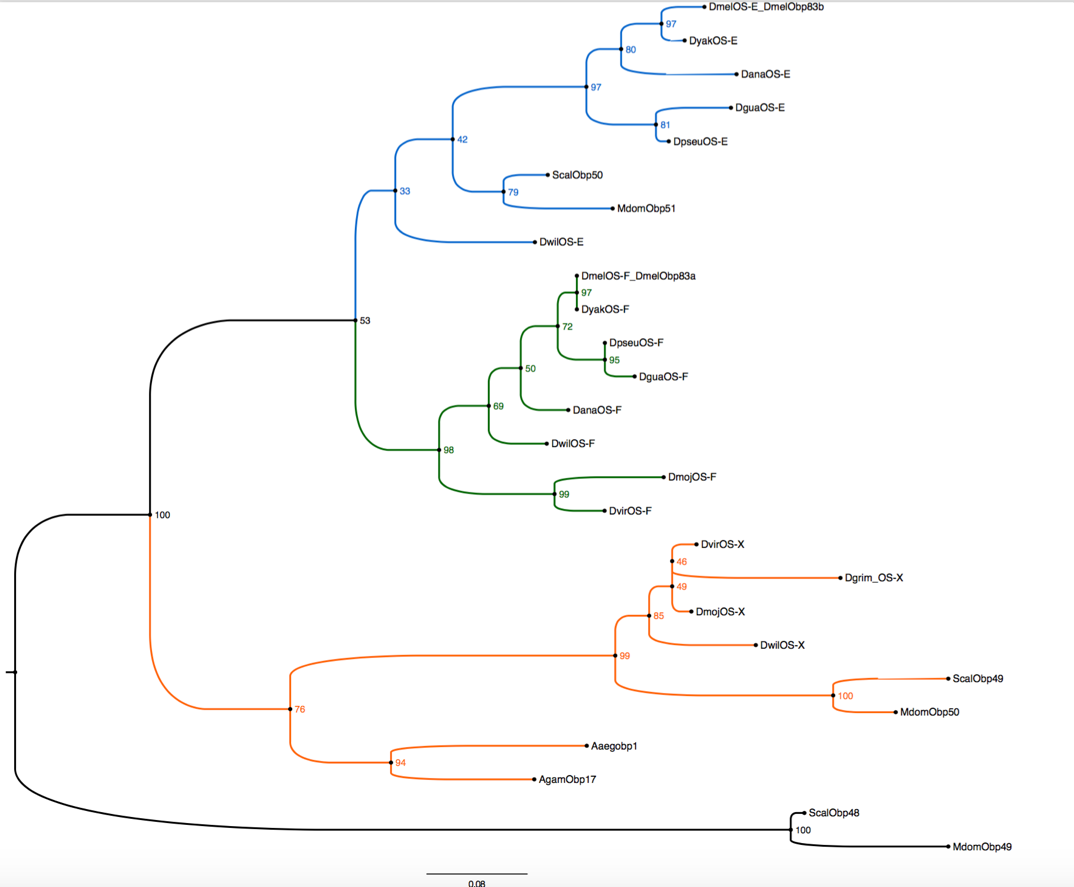

**Figure. S6.** **OS-E-like and an OS-X orthologue in *Stomoxys* and *Musca*.** Phylogenetic analysis with OS-E, OS-F, and OS-X proteins from *Drosophila* sp.

Eighty-eight *Obp* transcripts were detected in RNASeq from at least one tissue, and expression was not limited to olfactory organs with transcripts detected in both female and male reproductive systems (Fig. S5.B, Additional File 2, Table S22. This was not unexpected, as *Obp* expression in non-sensory tissues and roles not related to chemosensation have been reported for dipteran species [106, 107].DmelObp28a, orthologs of which are expanded in *Stomoxys* (ScalObp6-9) and *Musca* (MdomObp5-14), is abundant in a single sensillum on *Drosophila* antennae but does not appear to be essential for chemosensation in the classical sense [108]. Rather, DmelObp28a and its orthologue in *Glossina morsitans*, GmObp6, have a unique role in the bacterial-induced hematopoietic pathway that ultimately regulates melanization [109]. A similar non-sensory role was postulated for *DmelObp99b*, of which ScalObp60 and MdomObp59 are orthologs, as its expression in fat body is induced in response to activation by transcription factors that are typically associated with promoting fly longevity and survival [110]. A blowfly protein (PregObp56a) with orthology to DmelObp56a binds fatty acids, possibly functioning to transport byproducts from decaying meat to the blowfly gut for absorption and eventual use in reproduction [111]; orthologs in *Stomoxys* are expanded (*ScalObp18-21*).

Of interest are several muscid-specific lineages, the largest of which was mentioned above (Fig. S5; orange branches) and another that includes 7 *ScalObps* (*ScalObp79-85*) and 11 *MuscaObps* (*Mdom77-86*). Expression of these transripts in *Stomoxys* is diverse, suggesting the members serve multiple functions. Smaller musicd-specific lineages were present, including: ScalObp56-59/MdomObp56-57; ScalObp78/MdomObp76.

**The Odorant Receptor (OR) Gene Family**

Annotated odorant receptor sequences from *Drosophila melanogaster* (62 ORs) and *Musca domestica* (87 ORs) were used in tBLASTN searches of the *Stomoxys* genome to identify orthologs. This resulted in 74 ScalOr gene models, 71 of which were built by the NCBI automated annotation pipeline (Fig. S7; Additional File 2, Table S21; Supplementary Dataset 4). Three new gene models were constructed, not including ones that resulted from fixing or splitting a model or from joining across scaffolds. Four gene models were short and included premature stop codons, and these are predicted to be pseudogenes identified with PSEU. An additional three gene models were partial, missing either the C terminal or N terminal portions, identified with CTE or NTE, respectively. The *Stomoxys* OR sequences were named based on relationship to *Musca*, as it is the closest relative with a sequenced genome. As in *Musca*, the OR naming system starts with the ortholog of DpOrN. The highly-conserved odorant co-receptor ORCO is present in *Stomoxys* (ScalORCO) and has high sequence similarity to *Drosophila* (88%) and *Musca* (96%). The transcript is detected in all life stages and tissues evaluated, including reproductive systems from mated females and males, supporting previous reports [112].

Sixteen simple 1:1:1 OR orthologs are shared between *Stomoxys, Musca*, and *Drosophila*, and these have relatively high amino acid sequence similarities. ScalOr64/MdomOr78 are orthologous to DmelOr85e (60% similarity), which detects 1R-(-)fenchone, a monoterpene that has a repellent effect towards insects [113]. *DmelOr85e* is co-expressed with *DmelOr33c*, which does not appear to have an ortholog in either *Stomoxys* or *Musca*. ScalOr10/MdomOr11 and ScalOr11/MdomOr12 are orthologs of DmelOr10a (60% similarity) and DmelOr13a (70% similarity) that are known receptors for methyl benzoate [114] and 1-octen-3-ol (octenol) [115], respectively. Methyl benzoate is a component of clove bud essential oil, known to be repellent towards *Stomoxys* [116], while octenol is a cattle-associated compound known to elicit strong EAG responses [117]. *DmelOr10a* is tandemly arranged and co-expressed with *DmelGr10a* [118]. In *Stomoxys*, DmelGR10a is dramatically expanded into 48 proteins, 30 of which surround *ScalOr10* at this locus; it is unclear whether any are co-expressed with *ScalOr10*. ScalOr60/MdOr70 are orthologs of DmelOr82a (50% similarity), which is known to selectively respond to geranyl acetate [119]; geranyl acetate elicits a moderate EAG response in *Stomoxys* (Hieu et al., 2014). ScalOr34/MdomOr42 are orthologs of DmelOr49b (63% similarity), which is responsive to *o*-, *m*-, and *p*-cresols [119], methylphenols that are volatile components of cattle and cattle dung and elicit strong responses in *Stomoxys* [120, 121]. ScalOr21/MdomOr23 and ScalOr15/MdomOr16 are orthologs of DmelOr43a (57%) and DmelOr24a (60%). DmelOr43a and DmelOr24a are receptors for 1-hexanol and propyl acetate, respectively [122]; 1-hexanol is typically identified from blends of green plant volatiles, which can comprise decaying vegetation of *Stomoxys* larval breeding substrates, while propyl acetate is a decomposition product of cattle manure. ScalOr62/MdomOr75 is orthologous to DmelOr85d (45%), which is a receptor for ethyl pentanoate and 2-heptanone [123], the latter of which elicits a weak EAG response in *Stomoxys* [124]. *Stomoxys* has several ORs that are orthologous to larval-specific or larval-expressed *Drosophila* ORs, yet do not share the larval expression pattern. These include: *ScalOr6* (female/male antennae, male proboscis)/DmelOr2a (larval, adult)/MdomOr4; *ScalOr43* (female/male antennae and proboscis)/DmelOr63a (larval)/MdomOr49; *ScalOr46* (female/male antennae)/DmelOr67b (larval, adult)/MdomOr51-52; *ScalOr58* (female/male antennae and proboscis)/DmelOr35a (larval, adult)/MdomOr67/68. *ScalOr66-67*/DmelOr94a-94b (co-expressed, larval-specific)/ MdomOr80. Other 1:1:1 orthologous relationships were: *ScalOr45*/DmelOr22c/MdomOr15; *ScalOr65*/DmelOr88a/MdomOr79.

ScalOr61 shares 50% identity with DmelOr85b and DmelOr85c, which are tandemly arranged in *Drosophila*. In *Musca*, *MdomOr71-74* are duplicated in tandem relative to *DmelOr85b/c*. *Stomoxys* microsynteny analysis supports orthology with *Drosophila*. However, *ScalOr61* and the neighboring TMEM135 domain containing gene are the only models on this scaffold possibly due to the presence of numerous gaps preventing meaningful gene model annotation. *DmelOr85b* is co-expressed with *DmelOr98b* in adults, and it is a receptor for butyl acetate and E3-hexenol; there does not appear to be an ortholog of *DmelOr98b* in either *Stomoxys* or *Musca*. *DmelOr85c* is expressed in larvae and is a receptor for 3-octanol and 1-heptanol [125]. *ScalOr61* expression was limited to adult tissues (adult heads, female/male antennae, male proboscis).

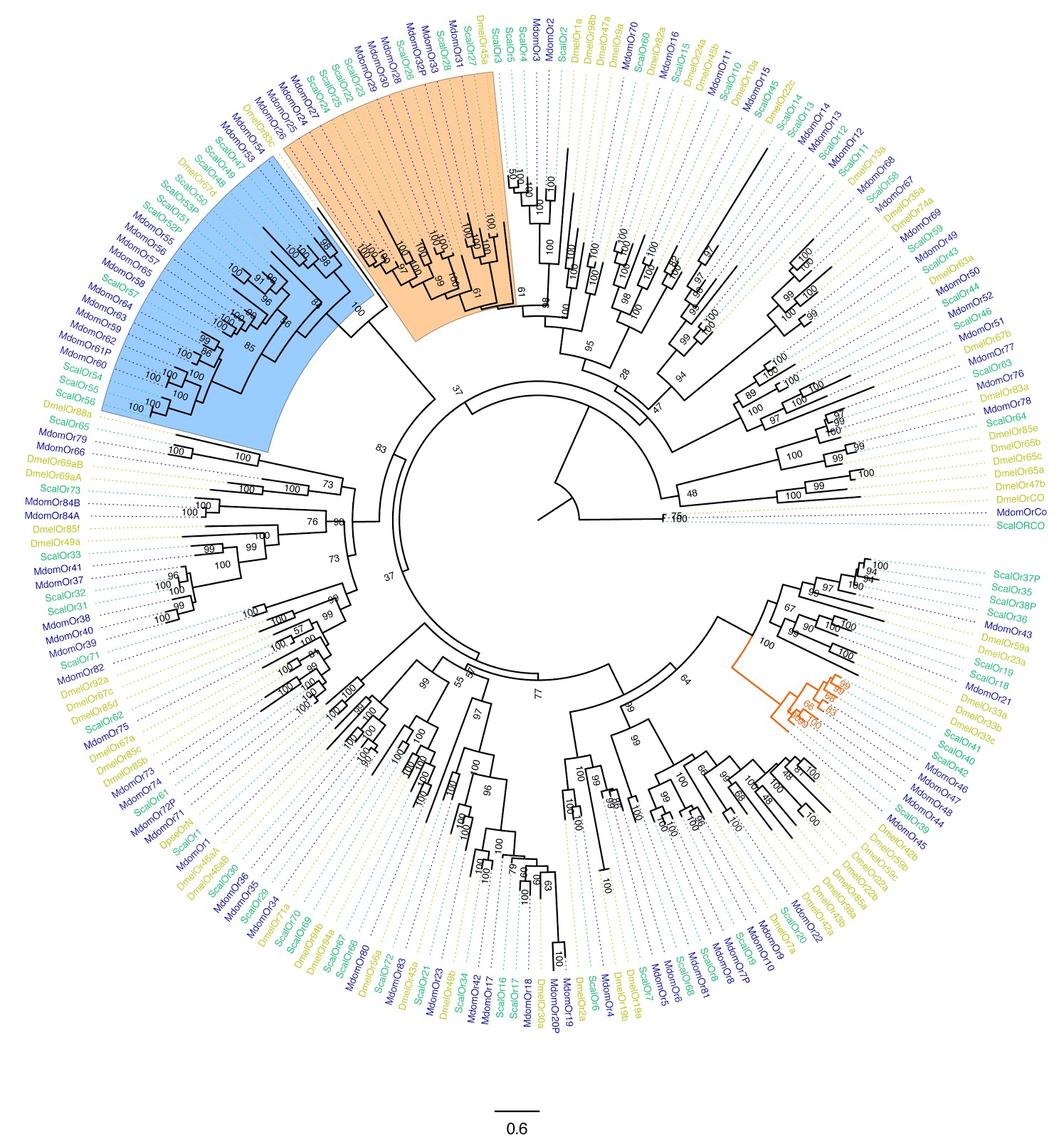

**A**

**Figure S7. Stomoxys Odorant Receptor Gene Family. A.** Phylogenetic tree of the *Stomoxys calcitrans* ORs with those of *Drosophila melanogaster* and *Musca domestica.* The *S. calcitrans* and *M.* *domestica* gene/protein names are highlighted in teal and blue, respectively, while *D. melanogaster* names are in mustard. Maximum likelihood phylogeny was constructed using the web server version of IQ-TREE software (Trifinopoulos et al., 2016; best-fit substitution model, branch support assessed with 1000 replicates of UFBoot bootstrap approximation). **B.** Heat map of normalized expression values for *Or* transcripts annotated from the *Stomoxys* genome.

**Figure S7 (contd).**

**B.**

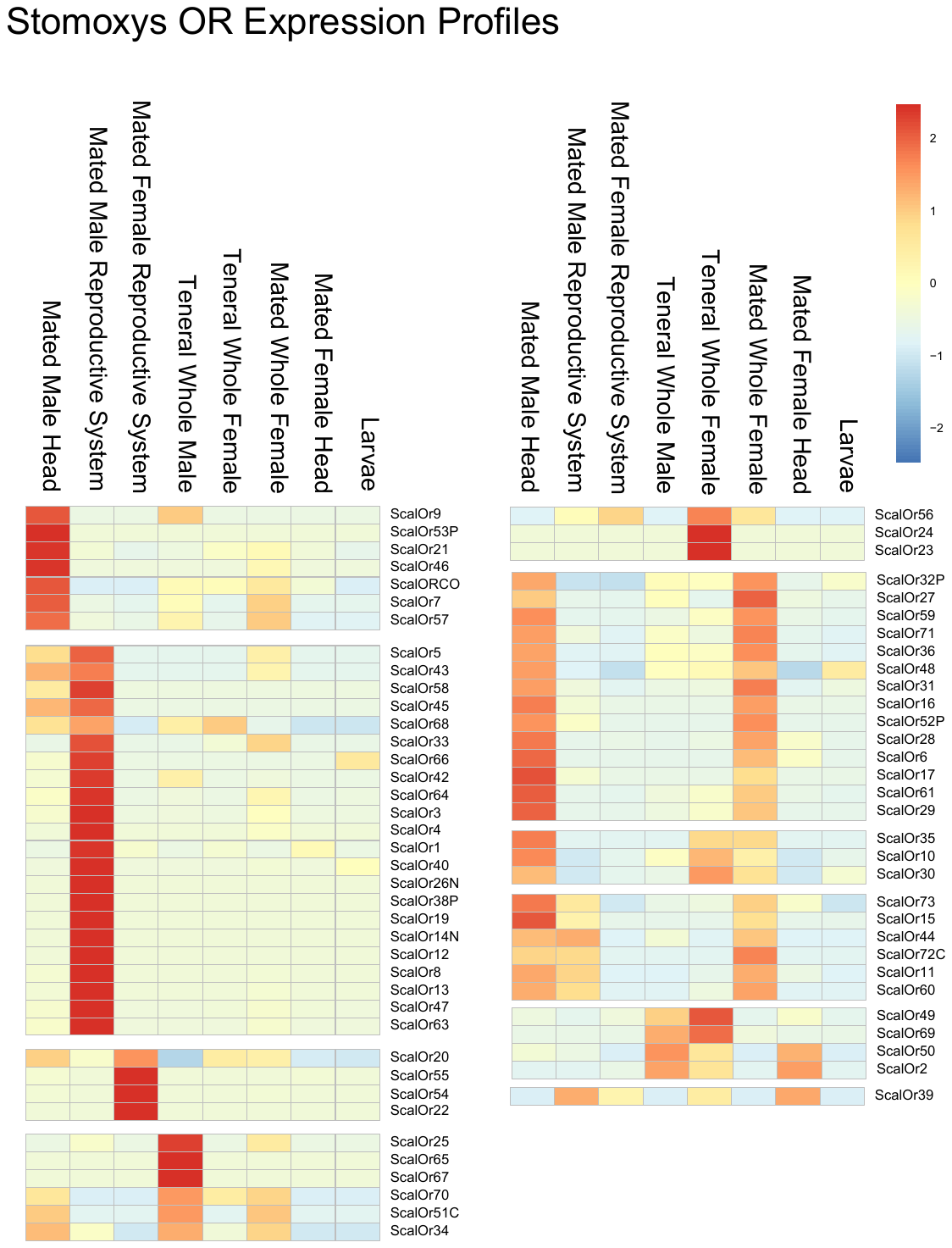

Smaller subsets of *Stomoxys* ORs duplicated relative to *Drosophila* are present in the genome. ScalOr35-38 clusters with DmelOr59a, which is a larval-specific receptor that responds to *o*-cresol. Within this group, ScalOr37 and 38 are pseudogenes and no larval RNASeq is detected for *ScalOr35* and *36* although *ScalOr36* was detected in female/male antennae and male proboscis. *ScalOr16* and *17* are expressed in female/male antennae but not in larvae, and these cluster with DmelOr30a, which is a larval-specific receptor that responds to *o*-cresol and *p*-cresol. ScalOr29-30 is in a sister clade to DmelOr49a A and B, which are the products of alternatively spliced *DmelOr49a* that is expressed in larvae and adults and binds *p*-cresol. There is no evidence for alternative splicing to produce these *Stomoxys* receptors. Although these *Stomoxys* receptors do not share a larval expression pattern with *Drosophila*, the response to methyphenols prevalent in cow dung could be co-opted for adult behaviors to locate hosts or ovipositional sites. Additional duplications relative to Drosophila ORs include ScalOr31-33/MdomOr37-41 that are in sister clades to DmelOr49a and DmelOr85f. *ScalOr31-33* are tandemly arranged, and *ScalOr33* is expressed in female/male antennae and proboscis. *DmelOr49a* and *DmelOr85f* are actually co-expressed, but unlinked in the genome (on different chromosomes), and together these mediate adult *Drosophila* detection and ultimate avoidance of semiochemicals produced by a parasitoid wasp, allowing the fly to detect ‘danger’ [126]. *ScalOr69* and *70* are tandemly arranged on a single scaffold and are duplicate orthologues of *DmelOr71a*, which is expressed in adults and binds ethylguaiacol and eugenol, components of host odors. ScalOr2 – 5, arranged on a single scaffold, are duplicates related to DmelOr1a, which is larval-specific, yet there is no evidence for larval expression of these *Stomoxys* receptors.

Muscid-specific lineages were also present, the most expansive being *ScalOr39-42* (single scaffold)/Mdom44-48. Others included *ScalOr7*/MdomOr5-6; *ScalOr8*/MdomOr7P-8; *ScalOr9*/MdomOr9-10; *ScalOr68*/MdomOr81; *ScalOr72*/ MdomOr83; *ScalOr73*/MdomOr84(A/B)

Fifty-seven of the 74 OR transcripts were detected in female and male head tissue, and a subset of 39 *ScalOrs* were further evaluated by non-quantitative RT-PCR to define spatial expression in pooled antennae and proboscides from fed adult females and males (Additional File 2, Table S21). As expected, all were detected in antennae, 17 were detected in both female and male proboscides, and an additional 13 were detected in only male proboscides, supporting an olfactory role for the proboscis beyond processing gustatory input, as has been suggested for mosquito species [127]. RNASeq detected 11 *Ors* in third instar larvae, and further evaluation by non-quantitative RT-PCR detected an additional 9 *Or*s expressed in first and second instar but not in third instar larvae. This suggests stable flies differentially utilize odorant receptors throughout immature development; all 20 of these *Ors* were not exclusive to the larval stages, as they were detected in adult tissues as well. Absence of larval-specific receptors in the stable fly may be a result of exposure to related compounds during the immature and adult stages, e.g. host dung, detritus. This is in contrast with mosquito species that occupy a larval aquatic habitat that is distinct from that of the adult. Similarly, both female and male stable flies are optional blood feeders and sex-biased receptors to enhance host localization in one gender over the other may be less critical.

**Chemosensory Proteins (CSP)**

Chemosensory proteins (CSPs), also referred to as OS-D like and sensory appendage proteins, are a class of small, highly soluble molecules that have been associated with insect sensory organs and other non-chemosensory tissues (Wanner et al., 2004). CSPs have no sequence similarity to members of the OBP family, and they encode four cysteine residues believed to play a role in disulfide bridge formation. A diverse functional role for this family is proposed due to its expression in various tissues, such as pheromone glands (Gong et al., 2007) and the Drosophila ejaculatory duct (PebIII), making them attractive as targets for control. Seven CSPs were previously described, and manual curation identified 3 additional CSP sequences (Supplementary Dataset 5). Eight of these ten CSPs are tandemly arranged on a single scaffold (KQ080226; Additional File 2, Table S20).

***Stomoxys* Vision**

**Contributors**: Markus Friedrich, Jeffery Jones, and Justin Dykema (Wayne State University)

**Table S23. Stomoxys opsin gene compilation.**

| **Organism** | **Homolog name** | **VectorBase ID** | **NCBI Gene ID** | **Gene ID** |
| --- | --- | --- | --- | --- |
| Stomoxys calcitrans | Rh1.1.1.1 | SCAU002283 | XM_013259150.1 | LOC106092322 |
| Stomoxys calcitrans | Rh1.1.1.2.1 | SCAU002003-RA | XM_013259153.1 | LOC106092325 |
| Stomoxys calcitrans | Rh1.1.1.2.2 | SCAU002003-RB | XM_013259155.1 | LOC106092325 |
| Stomoxys calcitrans | Rh1.1.2 | SCAU010102 | XM_013259152.1 | LOC106092323 |
| Stomoxys calcitrans | Rh1.2.1 | SCAU009433 | XM_013259157.1 | LOC106092326 |
| Stomoxys calcitrans | Rh1.2.2 | SCAU001182 | XM_013259158.1 | LOC106092327 |
| Stomoxys calcitrans | Rh2 | SCAU000443-RA | XM_013258302.1 | LOC106091691 |
| Stomoxys calcitrans | Rh3 | SCAU016603 | XM_013258948.1 | LOC106092178 |
| Stomoxys calcitrans | Rh5 | SCAU010746 | XM_013262197.1 | LOC106094954 |
| Stomoxys calcitrans | Rh6 | SCAU014862-RA | XM_013243463.1 | LOC106081480 |
| Stomoxys calcitrans | Rh7 | SCAU012939-RA | XM_013258762.1 | LOC106092023 |

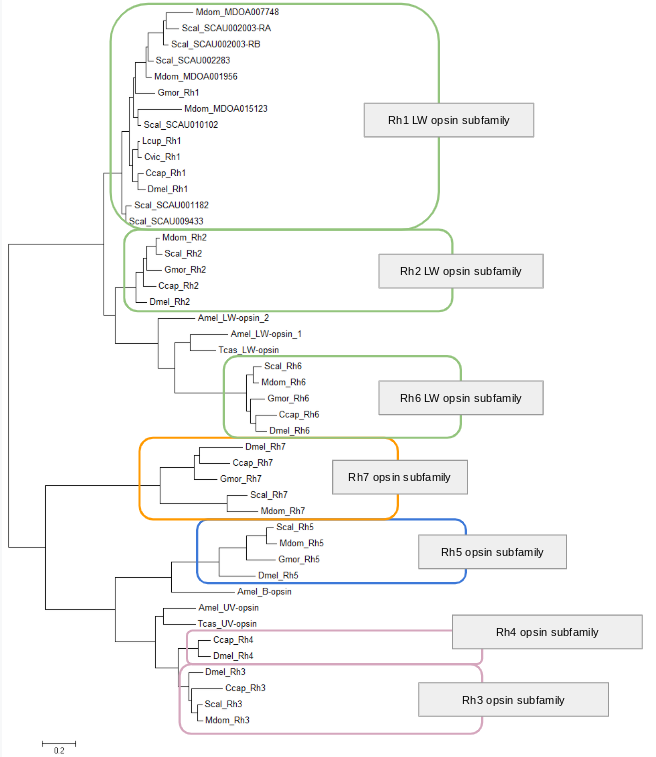

**Figure S8. Maximum likelihood tree of dipteran opsin gene relationships.** Protein sequences were aligned with Muscle [149]. Ambiguous alignment regions were filtered using Gblocks [150] using least stringent settings resulting in a final alignment with 291 sites.Bayesian tree analysis was performed out with MrBayes v3.2.6 [128] in the CIPRES Science Gateway V 3.3 environment [129], applying the GTR model of protein sequence evolution and correcting for across site substitution variation with a four rate category gamma distribution. All dipteran opsin ortholog clades were supported with credibility values higher than 0.9. . Species abbreviations: Amel = *Apis mellifera*, Ccap = *Ceratitis capitata*, Cvic = *Calliphora vicina*, Dmel = *Drosophila melanogaster*, Gmor = *Glossina morsitans*, Lcup = *Lucilia cuprina*, Mdom = *Musca domestica*, Scal = *Stomoxys calcitrans*, Tcas = Tribolium castaneum. Alignment available on request.

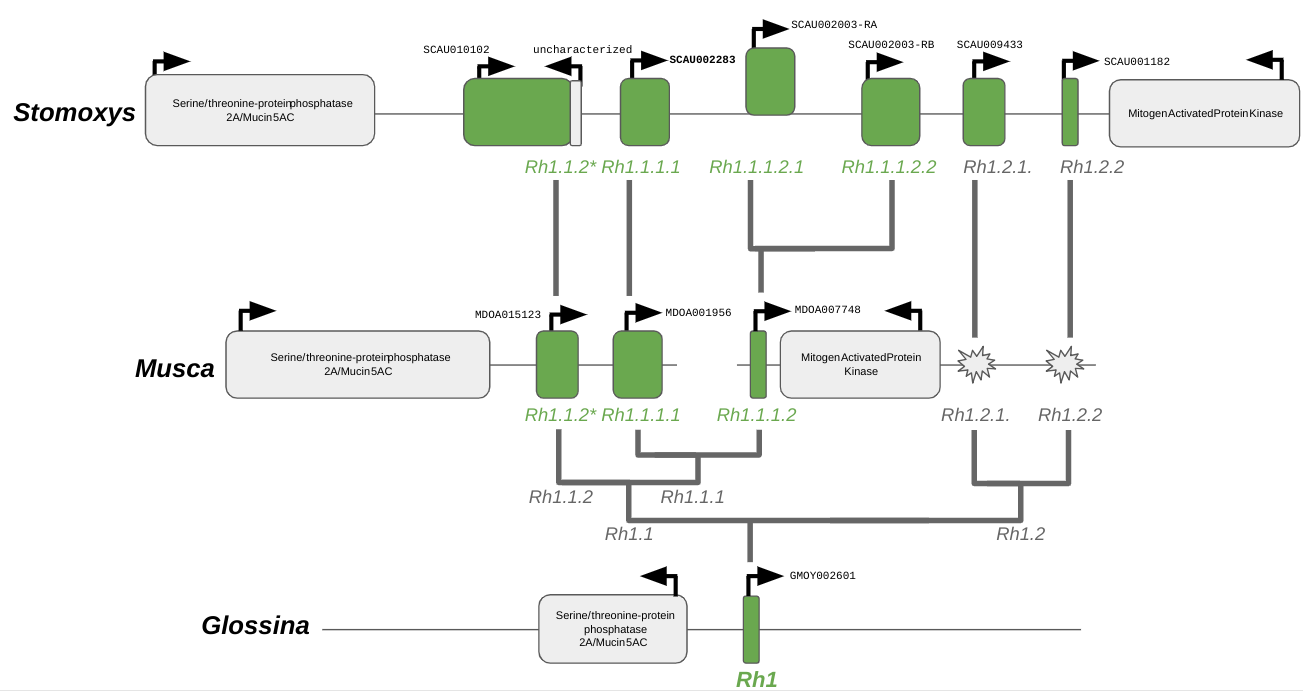

**Figure S9. Genomic organization and evolution of the Stomoxys Rh1 opsin subfamily in comparison to Musca and Glossina.**

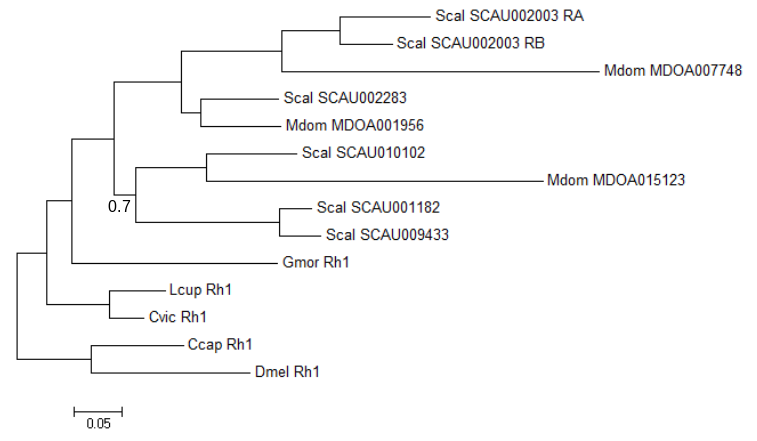
 **Figure** **S10: Phylogenetic analysis of the Stomoxys Rh1 gene cluster.**Bayesian analysis of the calyptrate expansion of Rh1 opsins. Protein sequences were aligned with Webprank [152]. Ambiguous alignment regions were filtered using TrimAl (v. 1.3) [39] as implemented on the Phylemon 2.0  server [153] applying User defined settings (Minimum percentage of positions to conserve: 10, Gap threshold: 0.9, Similarity threshold: 0.0, Window size: 1.0). Bayesian gene tree estimation and species abbreviations same as for Figure S9. All branch credibility values 1 except for one internal branch with 0.7 as indicated. Alignment available on request.

**Table S24. Opsin homolog expression levels in *Drosophila*, *Stomoxys* and *Musca*.** Transcription levels normalized as Fragments Per Kilobase of transcript per Million mapped reads (FKPM) for female and male head transcriptomes.. Data from this study and [30].

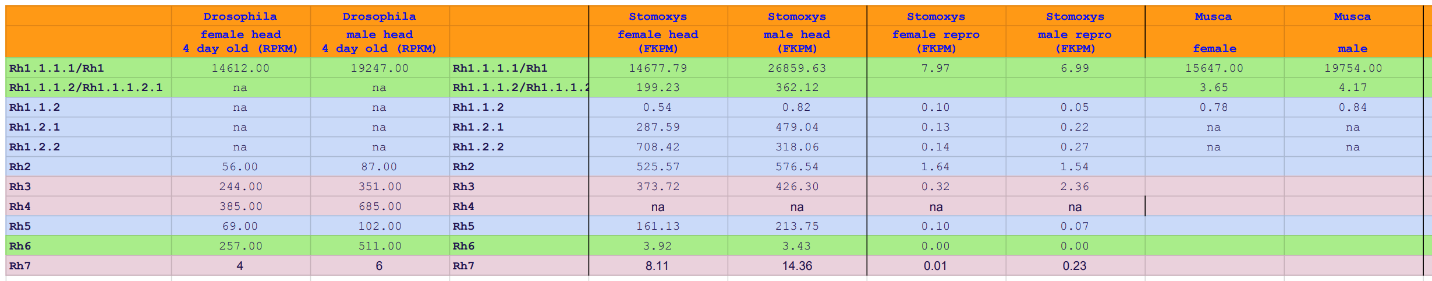

```
                             1
                             7
Scal_Rh1.1.1.1    DPMWNKILAAYLLTIGILAWIGNGTVIYIFGTTKSLRTPANLLVINLAVS
Mdom_Rh1.1.1.1    DPIWSKILAAYLLTIGILAWIGNGTVIYIFGTTKSLRTPANLLVINLALS
Scal_Rh1.1.1.2.1  DPMWNKILAIYLVVIGILAWIGNGTVLYIFATTKSLRTPANLLVINLALS
Scal_Rh1.1.1.2.2  DPIWYKILSTYLFTIGILAWIGNGTVIYIFGTTKSLRTPANLLVINLAIS
Mdom_Rh1.1.1.2    DPIWNKILTVYLIIIGMMAWFGNGTVIYIFATTKSLRTPANLLVINLAIS
Scal_Rh1.1.2      DRMWYNILTLYMVLIGIISWCGNGVVIYVFSTTKSLRTPANLLVINLALS
Mdom_Rh1.1.2      DREWYNLLTLYMLIIGIVSWCGNGVVIFIFSSSRALRTPANLLIINLALS
Scal_Rh1.2.2      DPMWNKILMWFMILIGIISWCGNGVVIYIFSTTKSLRTPANLLVINLALS
Scal_Rh1.2.1      DPMWNKILMWFMILIGIISWCGNGVVIYIFSTTKSLRTPANLLVINLALS
Lcup_Rh1          DPMWAKLLTAYMIVIGLISWCGNGVVIYIFSTTKSLRTPANLLVINLAIS
Cvic_Rh1          EPKWAKFLAAYMVLIATISWCGNGVVIYIFSTTKSLRTPANLLVINLAIS
Gmor_Rh1          DPMWNKILTTYMIMIGCISWCGNGVVIYIFSTTKSLRTPANLLVINLALS
Dmel_Rh1          DPIWAKILTAYMIMIGMISWCGNGVVIYIFATTKSLRTPANLLVINLAIS
Ccap_Rh1          DPMWAKILTAYMILIGTISWCGNGVVIYIFSTTKSLRTPANLLVINLALS
Dmel_Rh6          EPMWFGIIGFVIAILGTMSLAGNFIVMYIFTSSKGLRTPSNMFVVNLAFS
Ccap_Rh6          EQIWFHIIGFIITILGVMSLSGNFIVMYIFTSTRSLRTPSNIFVVNLAFS
Gmor_Rh6          EPLWFGIIGFIITVLGIMSLTGNFIVMYIFTSSKSLRTPSNMFVVNLAFS
Scal_Rh6          EPMWFGIIGFIITILGIMSLAGNFVVIYIFTSAKPLRTPSNMFVVNLAFS
Mdom_Rh6          EPMWFGIIGFVITVLGIMSLTGNFVVIYIFTSAKSLRTPSNMFVVNLAFS
Dmel_Rh2          DPMMSKILGLFTLAIMIISCCGNGVVVYIFGGTKSLRTPANLLVLNLAFS
Ccap_Rh2          DSTMSQILGLFTLVLLLISACGNGVVVYIFGGTKSLRTPANLLVLNLAFS
Gmor_Rh2          DTKMNQILGVFTFVIMVISLCGNGMVVFIFGSTKSLRTPANLLVLNLAFS
Mdom_Rh2          PSATSQLFGIFTAAIMVVSCCGNGVVVYIFGGTKSLRTPANLLVLNLAFS
Scal_Rh2          PSATSQLFGIFTAAIMVISCCGNGVVVYIFGGTKSLRTPANLLVLNLAFS
```

**Figure S11. Analysis of tuning site 17 variation in Stomoxys and Musca Rh1 paralogs.**

**Immune System**

**Contributors**: Tim Sackton (Harvard University), Pia Olafson (USDA-ARS), Dana Nayduch (USDA-ARS)

We use two computational approaches combined with targeted manual annotation to produce our annotation of immune-related genes and gene families in the *S. calcitrans* genome. First, we generated an initial list of *S. calcitrans* gene models that likely have an immune function based on homology to annotated and functionally characterized *D. melanogaster* genes. Homologous genes were identified based on OrthoDB groups, and include both orthologs and paralogs (e.g., every *S. calcitrans* gene in an OrthoDB group that includes a *D. melanogaster* immune-related gene is annotated as immune-related). To complement these homology-based annotations, we screened all predicted *S. calcitrans* proteins for similarity to HMM profiles of well-characterized Dipteran immune-related protein families (Additional File 2, Table S25) [130, 131], using hmmscan (from the HMMER software package). Finally, because antimicrobial peptides in particular can be difficult for computational pipelines to properly annotate (due to their small size), we also include some manual annotation of *S. calcitrans* AMPs. However, in order to maintain comparability to prior studies, we focus on the computational approaches for comparisons with *M. domestica,* *G. mortisans,* and other Dipterans. A full list of putative computationally annotated immune-related genes in *S. calcitrans*, including NCBI gene ids and putative functions when inferable from homology, is included in Additional File 2, Table S25. Major components of the Imd, Toll, and JAK/STAT pathways are summarized in Table S25.

In addition to our genome-wide computational annotation of immune genes, we focused our manual annotation efforts on antimicrobial peptides (AMPs), which can be challenging for *in silico* annotation algorithms to detect due to their short size (Additional File 2, Table S26; Supplementary Dataset 6). We focus on four classes of AMPs: defensins (DEF), attacins (ATT), diptericins (DIPT), and cecropins (CEC), all well-characterized and broadly distributed AMPs among insects. Four of the five computationally predicted defensins in the *S. calcitrans* genome are clustered on a single scaffold (KQ079966), and manual examination of that region of the genome reveals an additional six defensin gene models that were missed by the computational screen, including one previously reported (Stomoxys midgut defensin (Smd) 1, [132]) (Additional File 2, Table S26). Thus we believe that the *S. calcitrans* genome likely encodes at least 11 total defensins. The 12 predicted Attacins are largely clustered on four scaffolds [KQ080058 (5), KQ080155 (1), KQ080308 (4), and KQ082105 (2)]; one gene model (LOC106086445) was split to produce two better representative models. Of the three predicted diptericins, likely only one (LOC106086283) is in fact an antimicrobial peptide: while LOC106086281 does encode an ATTC domain, the size and gene structure (16 introns) is not consistent with other insect diptericins and no conserved domains are detectable in LOC106086726. Computation annotation recovers five cecropin-like sequences on a single scaffold (KQ080281). Three additional cecropin gene models were identified on this scaffold, along with two additional cecropin-like gene models with similarity to Stomoxyn [133] on scaffold KQ080227 (Additional File 2, Table S26), for a total of 10 cecropins in the *S. calcitrans* genome. Finally, we find a cluster of 9 gene models on scaffold KQ079975 that were *in silico* annotated as lncRNAs, but there is ample RNASeq support to indicate they encode short peptides with low sequence similarity to cecropin; these encode predicted signal peptides and alpha-helixes at the C terminus. In the absence of proteomic evidence it is difficult to know for sure whether these sequences are indeed translated into AMP-like peptides, but it seems likely that this cluster represents another set of cecropin-like AMPs in the *S. calcitrans* genome.

**Table S27**. Components of the immune deficiency, Toll, and JAK/STAT pathways identified from the *Stomoxys* genome

| **Immune Deficiency Pathway** | | |
| --- | --- | --- |
| PGRP-L | *See Additional File 2, Table S24* |  |
| Imd (immune deficiency) | XP_013107116 | Death domain superfamily; receptor-interacting serine/threonine-protein kinase 1 |
| FADD | XP_013114910 | Death domain; Death effector domain |
| DREDD (caspase-8) | XP_013105770 |  |
| Relish (nuclear factor NF-kappa p110 subunit) | XP_013099422 | Rel homology domain (RHD), IPT domain, ankyrin repeats |
| Relish (nuclear factor NF-kappa p110 subunit) | XP_013100924 | Partial, RHD, IPT, ankyrin |
| akirin | XP_013103618 |  |
| IAP (inhibitor of apoptosis) | XP_013112429 | 2 BIR domains, I ring_Ubox domain |
| TAB2 (tak associated binding protein) | XP_013103394 | Zinc finger, CUE-TAB2/TAB3 ubiquitin binding domain |
| TAK1  Transforming growth factor activated kinas (MAP3K) | XP_013107045 | STKc-TAK1 domain (catalytic domain of serine-threonine kinase) |
| IKK (ird5, IKK-beta) | XP_013109447 | SPS1 (serine/threonine protein kinase) domain |
| IKK (kenny, IKK-gamma) | XP_013118213 | UBAN motif |
| Poor imd response upon knock-in (PIRK); e.g. PIMS | XP_013117266 | e-32; no conserved domains, but Dmel PIRK doesn’t have any either |
| PGRP-SC2/3 | *See Additional File 2, Table S24* |  |
| Caspar (fas-associated factor 1, FAF) | XP_013102431 | Faf-1 UBX domain; UBA domain |
| caudal | XP_013114483 | Homeobox domain |
| mustard | XP_013098141 | TLDc domain; LysM domain |
| **Toll Pathway** | | |
| PGRP-SA, PGRP-SD | *See Additional File 2, Table S24* |  |
| Gram Negative Binding Proteins (GNBP)- GNBP3-like | XP_013116957 | Carbohydrate binding, Laminin G domains |
| GNBP3-like | XP_013108539 | Carbohydrate binding, Laminin G domains |
| GNBP1-like | XP_01311306 | Carbohydrate binding, Laminin G domains |
| GNBP2-like | XP_013113076 | Carbohydrate binding, Laminin G domains |
| spaetzle | XP_013115944 | Signal peptide |
| spaetzle processing enzyme (SPE) | XP_013102919 | Tryp-Spc, CLIP domain |
| Toll Receptor | XP_013107363 | N-terminal Leu-rich repeat, toll interleukin-1 resistance (TIR), TPKR_C2 domains |
| Myeloid differentiation primary response protein 88 (MyD88) | XP_013115653 | Death, TIR-2 domains |
| pelle (interleukin-1 receptor associated kinase [IRAK] 1) | XP_013102217 | C-terminal STKc_IRAK, N-terminal Death domains |
| tube (IRAK4) | XP_013116710 | Death domain |
| cactus (Rel-1 inhibitor) | XP_013100640 | Ankyrin repeats |
| dorsal | XP_013118266 | N-terminal Rel homology domain (RHD), C-terminal IPT_NKkB domain |
| dorsal immunity factor (dif) | XP_013118268 | N-terminal RHD, C-terminal Rel homology dimerization domains |
| deaf1 | XP_013116039 | SAND, MYND finger domain |
| drifter (dfr)/ventral veinless (vvl) | XP_013117919 | POU domain |
| persephone (psh) | XP_013119056 | Tryp_SPc (trypsin-like serine protease) domain |
| TNF receptor associated factor 6 (TRAF6) | XP_013107963 | RING, MATH, Sina domains |
| **JAK-STAT Pathway** | | |
| unpaired (upd); ligand of domeless | XP_013114134 | UPD domain |
| upd | XP_013099717 | UPD domain |
| upd4-like | XP_013099693 | UPD domain |
| domeless receptor | XP_013097750 | Transmembrane domains; cytokine receptor motif; Fibronectin type 3 (FN3) domain |
| domeless receptor | XP_013108583 | Transmembrane domains; cytokine receptor motif; Fibronectin type 3 (FN3) domain |
| hopscotch | XP_013112402 | TyrKc, SH2 JAK domains |
| suppressor of cytokine signaling (SOCS);  SOCS36E-like | XP_013114164 | C-terminal SOCS box, SH2 domain, phosphotyrosine binding pocket |
| SOCS16D-like | XP_013103755 | C-terminal SOCS box, SH2 domain, phosphotyrosine binding pocket |
| SOCS44A-like | XP_013104964 | C-terminal SOCS box, SH2 domain, phosphotyrosine binding pocket |
| Signal transducer and activator of transcription (STAT) | XP_013119408 | SH2-STAT domain, STAT DNA binding and STAT protein interaction domains |
| Protein inhibitor of activated STAT (PIAS) | XP_013104891 | MIZ-SP/RING zinc finger, PINIT domains |

**Table S28 (Table 1, main paper). Comparison of Insect Immune System Gene Families**
*numbers in parenthesis represent manually curated counts

|  |  | *stoCal* | *musDom* | *gloMor* | *aedAeg* | *droMel* |
| --- | --- | --- | --- | --- | --- | --- |
| canonical effectors | ATT | 11 (12)* | 10 | 4 | 1 | 4 |
|  | DEF | 5 (11)* | 7 (11)* | 0 | 4 | 1 |
|  | DIPT | 3 (1)* | 4 | 0 | 1 | 3 |
|  | CEC | 5 (10)* | 12 | 2 | 9 | 5 |
|  | LYS | 23 | 32 | 4 | 7 | 13 |
| non-canonical effectors | TPX | 5 | 6 | 6 | 5 | 8 |
|  | PPO | 19 | 23 | 4 | 25 | 10 |
|  | GPX | 1 | 1 | 0 | 3 | 2 |
|  | HPX | 12 | 12 | 8 | 19 | 10 |
|  | TSF | 4 | 6 | 3 | 5 | 3 |
| canonical recognition | NIM | 25 | 23 | 10 | 8 | 17 |
|  | PGRP | 17 | 17 | 4 | 10 | 13 |
|  | BGBP | 4 | 3 | 3 | 7 | 7 |
|  | TEP | 16 | 22 | 4 | 8 | 6 |
| other recognition | CTL | 78 | 41 | 11 | 43 | 38 |
|  | FREP | 49 | 38 | 7 | 34 | 14 |
|  | GALE | 15 | 13 | 8 | 12 | 6 |
|  | IGSF | 1 | 1 | 1 | 0 | 1 |
|  | MD2L | 8 | 12 | 5 | 26 | 8 |
|  | SRCA | 3 | 3 | 2 | 2 | 3 |
|  | SRCB | 15 | 18 | 11 | 13 | 14 |
|  | SRCC | 7 | 8 | 4 | 5 | 9 |

To better understand the biology of PGRPs in *S. calcitrans*, we examined patterns of gene expression from both RT-PCR assays in larval, pupae, and adult guts, and from whole genome RNA-seq assays of larvae and adults. We identified 17 PGRPs in the *S. calcitrans* genome (11 in the short subfamily, 5 in the long subfamily, and 1 ambiguous; Fig. S12). Of the members of the short subfamily, six – all homologs of the PGRP-SC gene family in *D. melanogaster* – appear to be only expressed in larvae (XP_013110193, XP_013110191, XP_013106717, XP_013106719, XP_013109922, and XP_013097596; Fig. S13), and based on sequence properties and conservation of residues required for amidase activity [134] are predicted to be both secreted and catalytic (Fig. S14). Interestingly, PGRP-SC genes in both *D. melanogaster* [135] and *M. domestica* [136] are also expressed in larvae where they likely have a role in modulating activation of the IMD pathway. Larval expression is not exclusive in *D. melanogaster*, though, suggesting the possibility that larval-specific expression may be a *S. calcitrans* innovation.

The remaining five members of the short family have more variable expression patterns, as well as variable evidence for amidase activity. The *Stomoxys* PGRP-SB orthologue (XP_013114626) likely retains catalytic activity, while the *Stomoxys* PGRP-SA (XP_013098814, XP_013098815, and XP_013098810) and –SD (XP_013108408) orthologues likely lack catalytic activity but may serve a receptor function [134]. The five members of the long subfamily, in contrast, all appear to have relatively broad expression patterns. The PGRP-LB orthologue (XP_013109943) encodes a predicted signal peptide and conserved residues supporting peptidoglycan binding and amidase activity, which is in keeping with the characterized *Drosophila* PGRP-LB that is regulated by the IMD pathway and cleaves peptidoglycan of gram negative bacteria [137]. The *Stomoxys* PGRP-LE (XP_013111331, XP_013111330) orthologues were identified at two loci tandemly arranged in opposite directions, and these are predicted to bind peptidoglycan but are likely non-catalytic, which has been demonstrated for *Drosophila* PGRP-LE [138]. The PGRP-LA orthologue (XP_013109991) does not encode residues associated with peptidoglycan binding, but may have a regulatory role [139], and the ambiguous gene (XP_013113924) encodes a PGRP-LD like polypeptide that is only very weakly expressed in any stage based on RNA-seq data; the predicted sequence does not support peptidoglycan binding. The expanded PGRPs in *S. calitrans*, then, are from the short subfamily, especially the SC-like members, and tend to have larval-specific expression patterns.

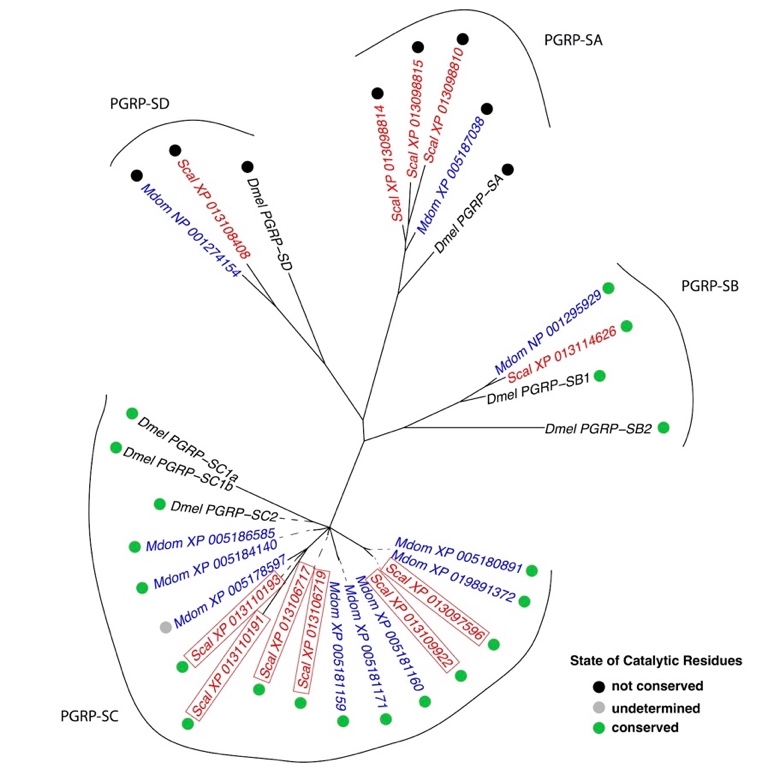

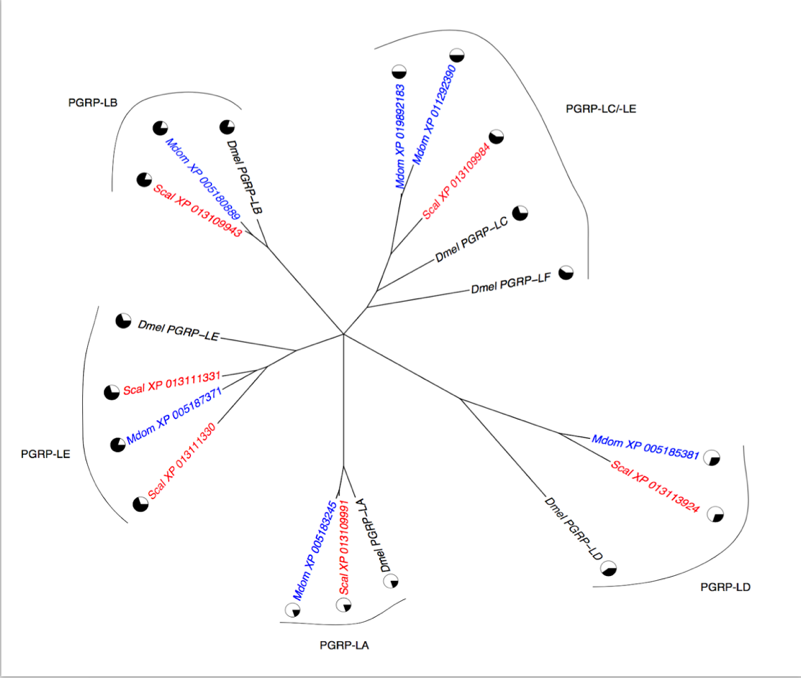

**Figure S12. Maximum likelihood phylogenetic tree of PGRP protein sequences from *S. calcitrans* (red), *M. domestica* (black), and *D. melanogaster* (blue)**. Alignments of PGRP-S and PGRP-L sequences were produced separately using mafft (einsi option), and then trimmed to remove columns with >75% gaps with trimal. Phylogenetic trees were computed with RAXML using the PROTGAMMAWAG model and bootstrapped 100 times. Nodes with < 50% bootstrap support are collapsed. Larval-specific *S. calcitrans* PGRP-S sequences are indicated with a box. Circles indicate presence/absence of catalytic residues (PGRP-S), or the proportion of sites important for PGRP binding conserved (PGRP-L).

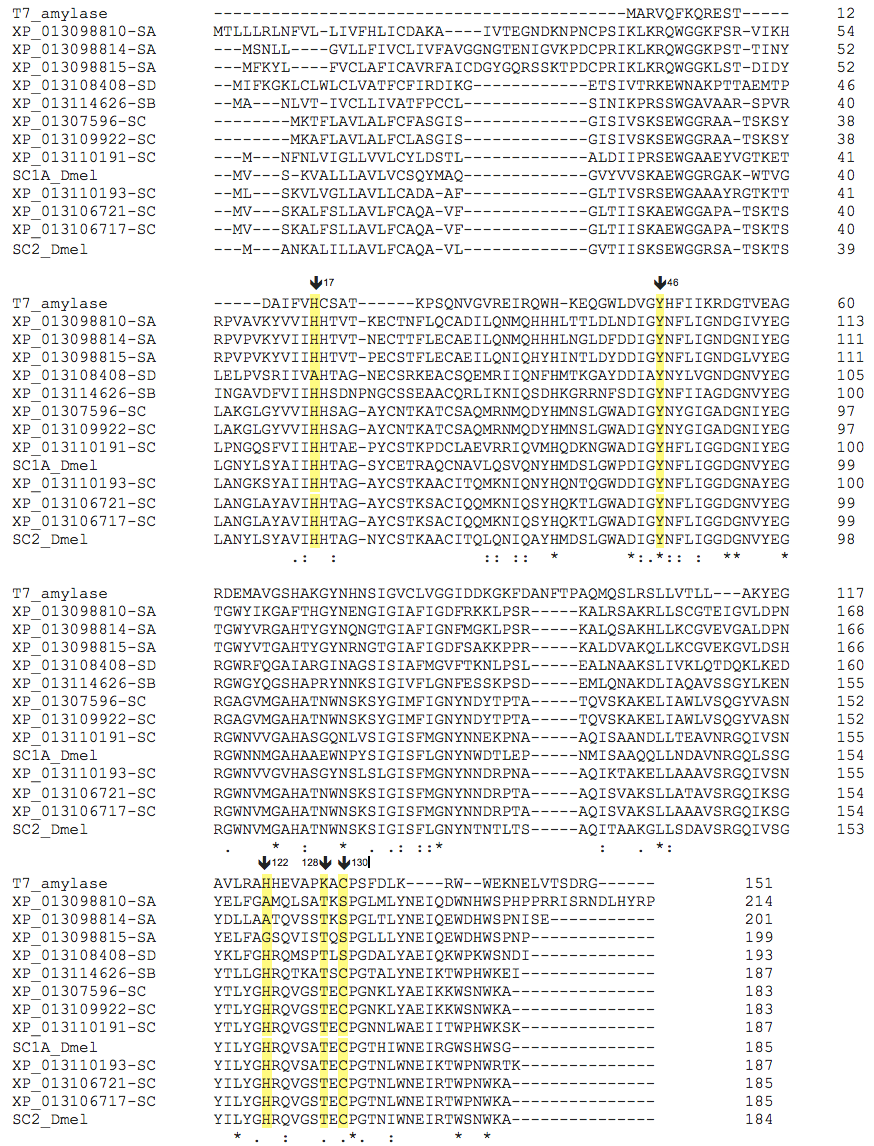

**Figure S13.** Alignment of *Stomoxys* PGRP-S sequences with characterized *D. melanogaster* PGRP-SC1 (C0HK98) and –SC2 (Q9VX2) and N-acetylmuramoyl-L-alanine amidase (P00806). Conserved amino acid residues required for catalytic amidase activity are highlighted in yellow [134]; conservation of cysteine at position 130 supports potential catalytic activity of *Stomoxys* PGRP-SC sequences.

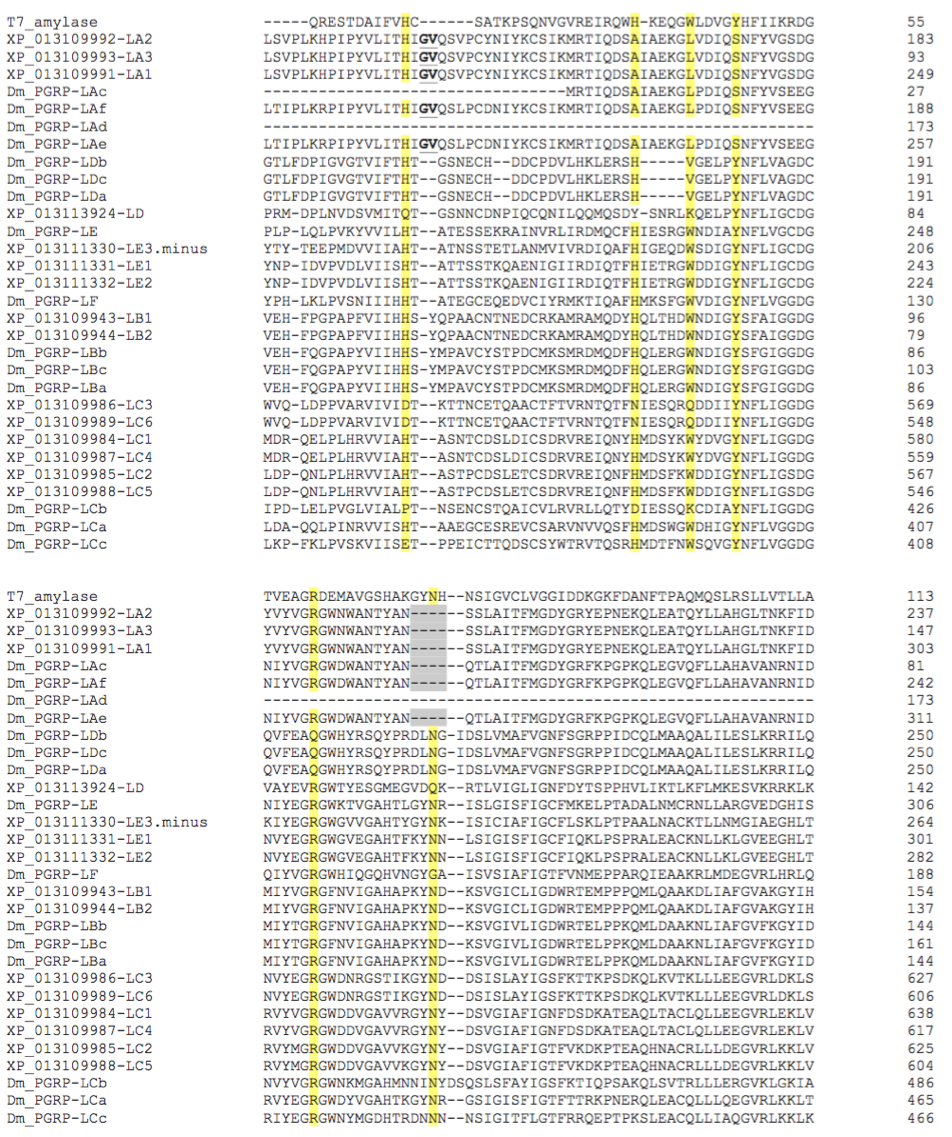

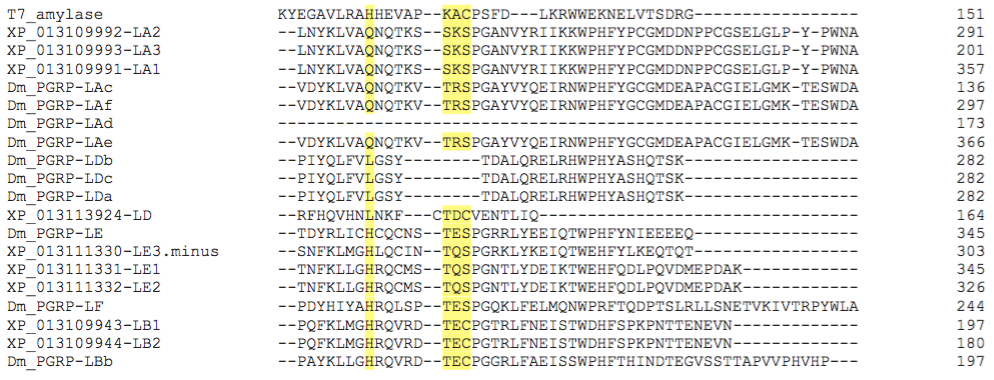

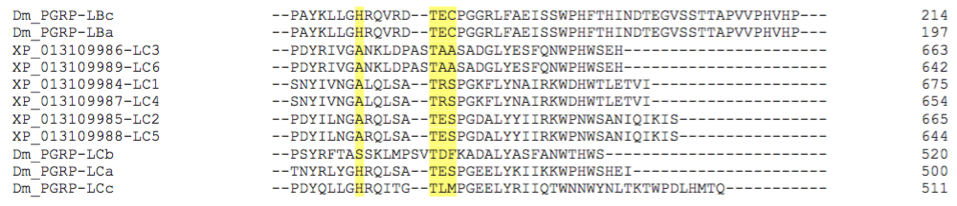

**Figure S14.** Alignment of *Stomoxys* PGRP-L sequences with characterized *D. melanogaster* PGRP-L proteins and N-acetylmuramoyl-L-alanine amidase. Conserved amino acid residues required for peptidoglycan binding and catalytic amidase activity are highlighted in yellow [134, 140]. Of the ten residues that are critical for peptidoglycan binding [140], the *Stomoxys* -LB, -LC and –LE sequences encode 9, 8, and 8 of these. Only 2 of these residues are conserved in the *Stomoxys* –LA and –LD sequences. Further, the *Stomoxys* -LA sequences encode a 2 amino acid insertion (bold, underlined text) and a 4 amino acid deletion (grey shaded) in regions of the molecules that are critical to binding [140]. These suggest that –LA and –LD do not bind peptidoglycan but may have a regulatory role [139].

Several of the expanded AMP gene families were found clustered on individual scaffolds, possibly arising from tandem duplications. The 11 *Stomoxys* defensin genes are located on a single scaffold (Fig. S15). Ten defensins were identified from scaffold KQ079966, which has a 3.3kb gap in the assembly, while a smaller scaffold (LDNW01121039) housed a single defensin gene model. Primers designed to amplify across the assembly gap confirmed that LDNW01121039 resides in this gap.

A)

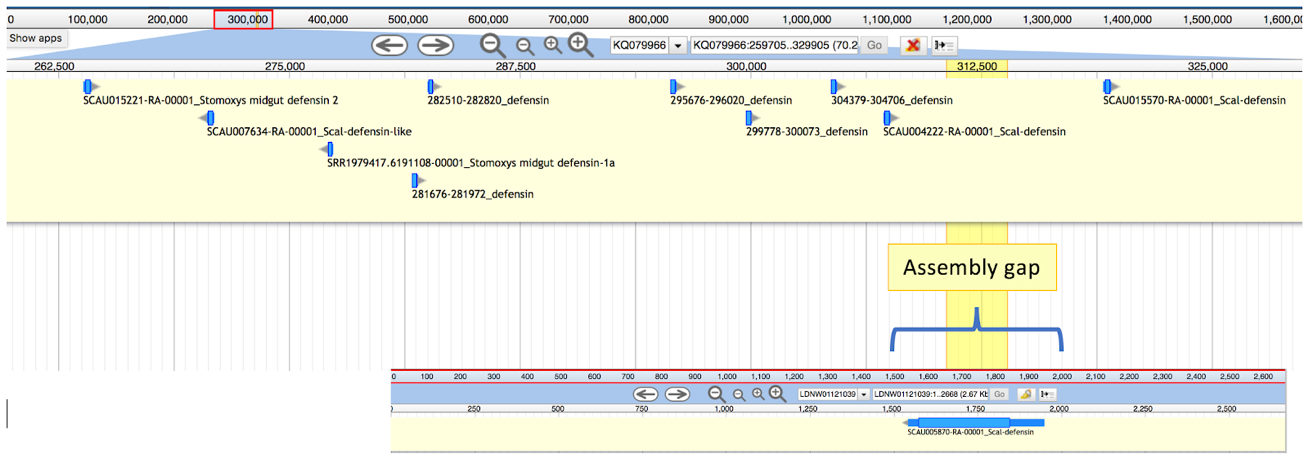

B)

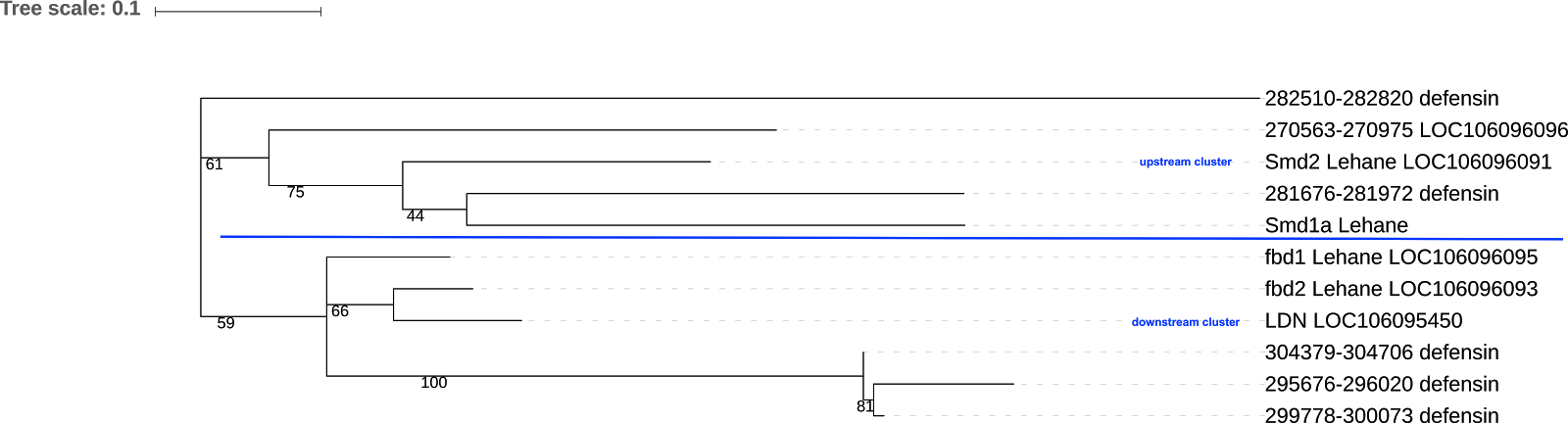

C)

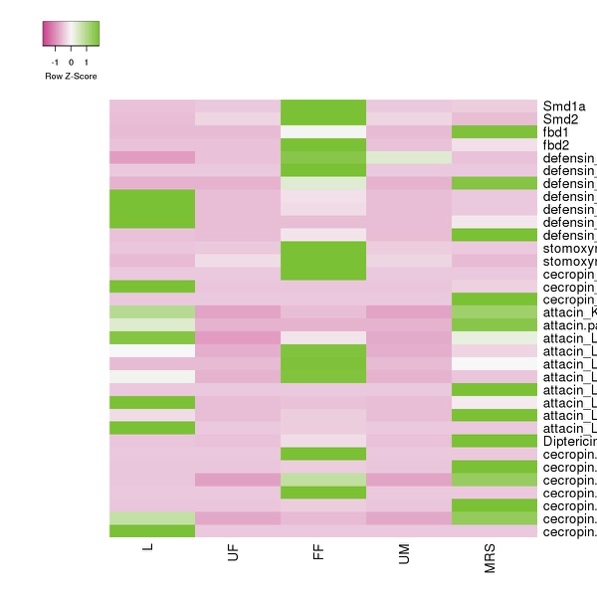

**Figure S15. *Stomoxys calcitrans* genomic scaffold housing 11 defensin gene models (A), phylogenetic relationship of defensins (B), and relative expression levels of antimicrobial peptide transcripts across tissues and developmental stages (C).**

Phylogenetically, the 11 defensins separate into three lineages (Figure S15.B), two lineages of which are occupied by five defensins present in an ‘upstream’ cluster from the remaining six defensins that form the third lineage. Within the downstream cluster, three *defensins* appear to be larval-biased (304379-304706, 295676-296020, and 299778-300073) and three are not detected in newly emerged adults, but are induced upon blood-feeding (fbd1, fbd2, LDN). Defensin genes in the upstream cluster are all detected in newly emerged adults and are upregulated in response to blood-feeding. Three defensins are also expressed relatively robustly in the male reproductive sytem (fbd1, 282510-282820, and LOC106095450), which is not unexpected given the difficulty in removing all of the fat body tissue that encapsulates the male testes. The *stomoxyn* expression pattern is consistent with previously published studies [133]. A cluster of five attacins were present on a single scaffold, and they displayed varying expression profiles: three appeared induced upon feeding in adult females, one was detected only in RTs of mated, adult males, and one was detected only in larvae.

Overall, our analysis of the *S. calcitrans* immune system as predicted by its genome reveals a dynamic system that recapitulates many of the trends first noted in *M. domestica*, suggesting that expanded repertoires of recognition and effector proteins may be a general feature of muscid fly genomes.

**Gene families associated with insecticide resistance**

**Contributors**: David Nelson (University of Tennessee) and Pia Olafson (USDA-ARS)

**Methods.**

Carboxylesterase, glutathione-S-transferase, and Cys-loop gated ion channel protein sequences from *Drosophila melanogaster* were used in tBLASTN searches to identify orthologs in the *S. calcitrans* 1.0.1 genome and M*. domestica* 2.0.2 genome assemblies. Amino acid sequences from each family were aligned with the MUSCLE algorithm [42], and the alignments trimmed with the trimAl tool using the –strictplus option [40]. The trimmed alignment was used to construct a maximum likelihood phylogeny with the web server version of IQ-TREE software ([43]; best-fit substitution model, branch support assessed with 1000 replicates of UFBoot bootstrap approximation). Resulting phylogenetic trees were unrooted.

**Results.**

Carboxylesterases (COE; EC 3.1.1) have a role in the detoxification of organophosphate, carbamate, and pyrethroid classes of insecticides [141]. Insecticide resistance conferred by COEs can occur by different mechanisms, including amplification of COE gene copy number, elevated expression of COE transcripts, and gene mutations that enhance COE enzymatic activity [142, 143]. The COE gene family in *Stomoxys* is comprised of 44 genes encoding 47 catalytic and non-catalytic proteins (Fig. S16; Supplementary Dataset 7). Thirty-six genes encode 38 secreted and intracellular catalytic proteins that separate into 7 clades, the largest of which includes 17 alpha-esterases (Additional File 1). Majority of the alpha-esterase like genes are clustered along a single scaffold (KQ079923) with only 3 genes located separately, one of which encodes a partial sequence. Separate clades with representative secreted beta-esterases, juvenile hormone esterases, integument esterases, and glutactins are also present in *Stomoxys* (Fig. S17). The COEs are presumed to be catalytic based on the presence of catalytic triad residues, with the exception of one glutactin protein (XP_013111473) and one juvenile hormone esterase-like protein (XP_013118535). Eight genes encode 9 non-catalytic proteins that are primarily involved in neurological development, including 4 neuroligins, gliotactin, and neurotactin. A single acetylcholinesterase gene is also present in the *Stomoxys* genome, as previously described by Temeyer and Chen [144].

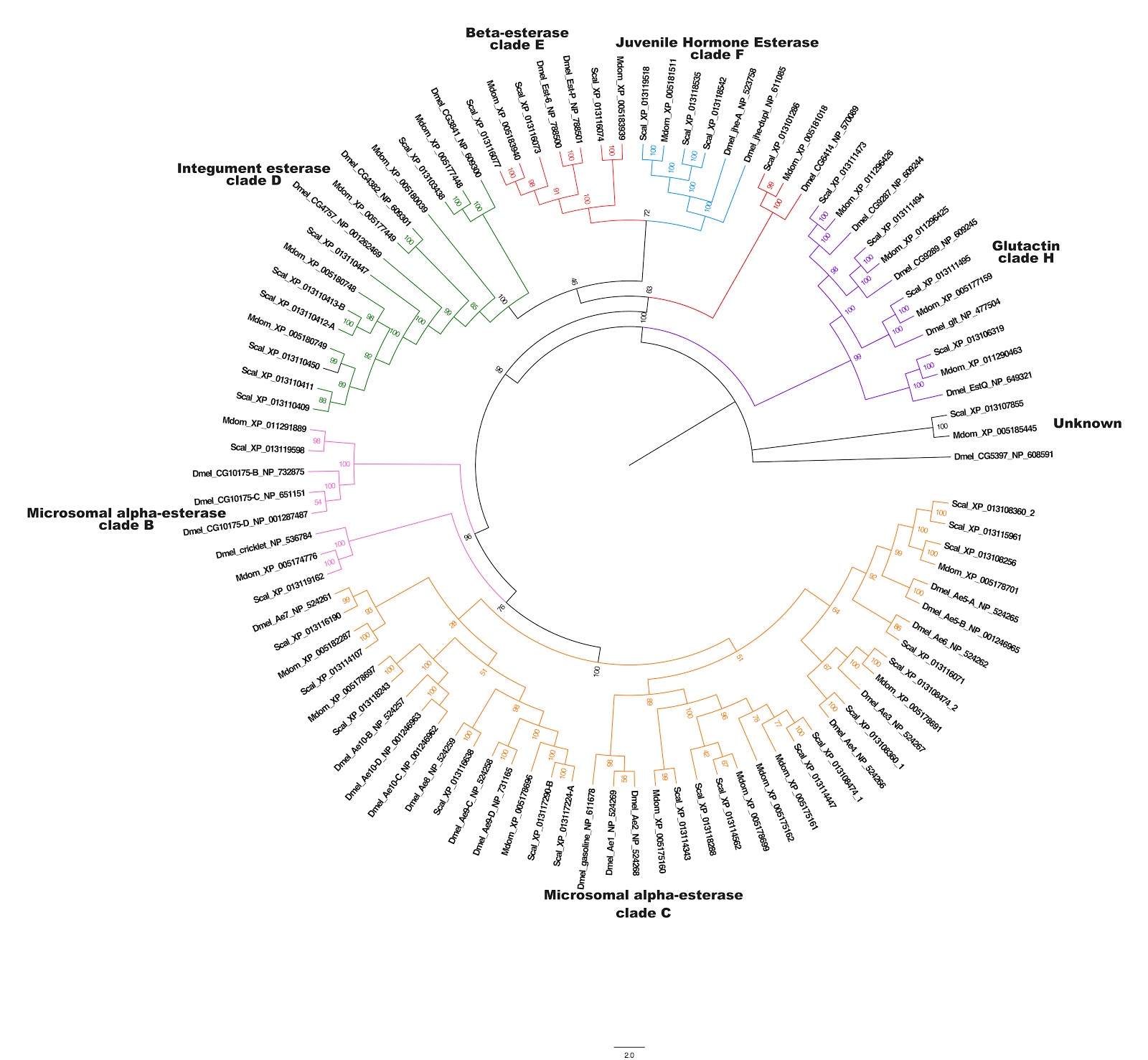

**Figure S16. Phylogenetic relationship of catalytic carboxyesterases in *Stomoxys*, *Musca*, and *Drosophila*.** Clades were designated according to nomenclature by Oakeshott et al. [145].

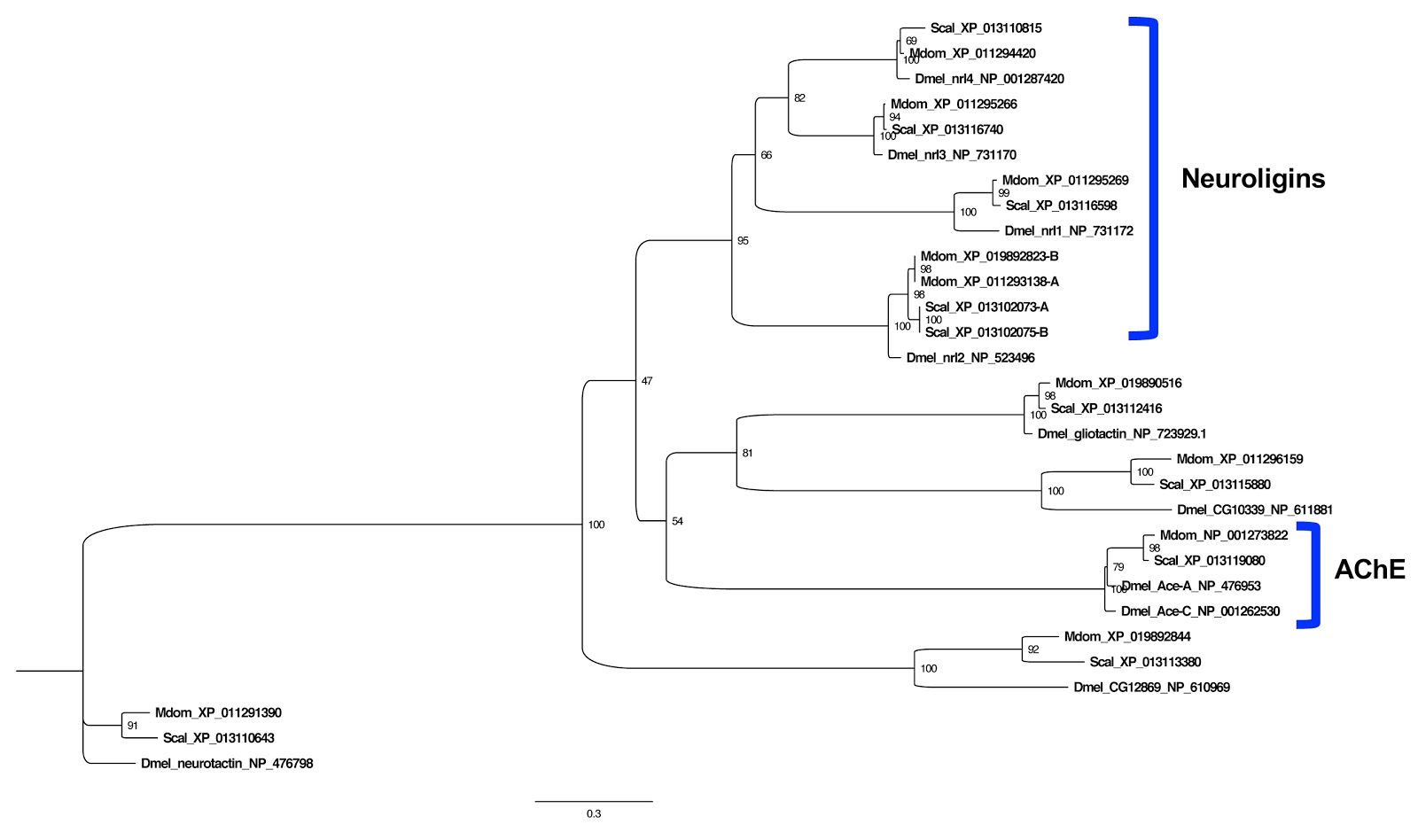

**Figure S17. Phylogeny of carboxylesterases with a role in neuronal development.**

Arthropod cytochrome P450s (CYPs) have diverse roles in insect physiology, including ecdysteroid biosynthesis and xenobiotic detoxification [146, 147]. The CYP gene family size varies among insects with dipterans having large arrays, i.e. 145 in *Musca*, 86 in *Drosophila*, and 77 in *Glossina*, and the 214 *Stomoxys* CYPs identified from the current genome assembly represent a substantial expansion relative to these sequenced dipteran genomes (Fig. S18; Supplementary Dataset 8; Additional File 2, Table S31). The *Stomoxys* CYPs have representatives from each of the CYP clans that are typically found in insects: mitochondrial, CYP2, CYP3, and CYP4. Approximately 17 were short fragments and not included in subsequent phylogenetic analysis. The *Stomoxys* mitochondrial and CYP2 clans contain orthologues of the Halloween genes that mediate the pathway for ecdysteroid biosynthesis, namely Cyp306A1, Cyp302A1, Cyp314A1, and Cyp315A1, and the mitochondrial clan (21 genes) contains tandemly arranged genes along three scaffolds: 9 CYP12A genes on KQ080363 and 7 CYP12G genes on KQ080439 and KQ082110. The CYP4 clan (62 genes) is primarily represented by the CYP4 (51 genes) family, while the CYP3 clan (107 genes) comprises the largest expansion of CYPs in the *Stomoxys* genome, predominated by the CYP6 (81 genes) and CYP9 (16 genes) families. Upregulation of CYP4, CYP6, and CYP9 genes has been associated with resistance to spinosad and pyrethroid insecticides in *Musca*, *Anopheles*, and *Drosophila* [148-150], but this has yet to be investigated in *Stomoxys*. Tandemly duplicated arrangements of 11 CYP4D (scaffold KQ080140), 18 CYP6A (scaffold KQ080692), and 16 CYP9F (scaffold KQ080085) genes are present in the *Stomoxys* genome. Given that *Stomoxys* inhabits conventional livestock production settings that utilize chemical fly control measures, the initial gene duplication that eventually lead to these expanded clusters may have been the result of environmental exposure to xenobiotic pressure [151].

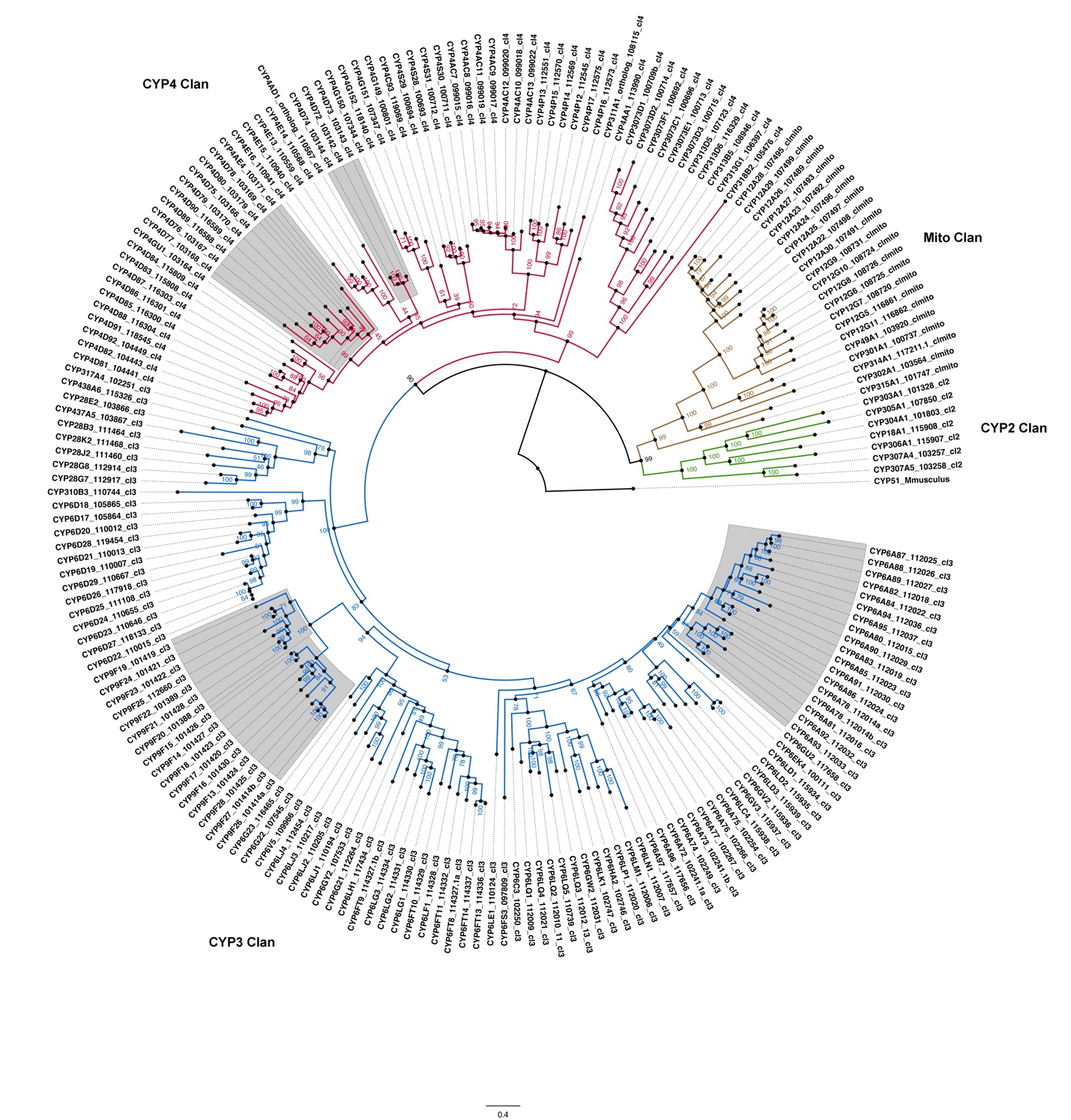
**Figure** **S18 (Fig. 9, main paper).** Phylogenetic analysis of cytochrome P450 genes from *Stomoxys*. The four insect CYP clades are presented in different colors, and examples of CYP gene clusters that are found in tandem within the genome are shaded in grey. P450 gene names were assigned by DR Nelson.

Glutathione S transferases (GST) are a family of enzymes that mediate the binding of reduced glutathione to xenobiotic or exogenous compounds that are electrophilic; this renders the compounds less reactive (detoxification) and promotes their excretion. GSTs typically encode residues for glutathione binding (G site) and substrate binding (H site), and these sites are located in the N- and C- terminus of the protein, respectively. Mutations within these binding sites and upregulation of GST expression have been correlated with resistance to insecticides, likely a result of increased capacity for xenobiotic detoxification [142, 152]. Insects encode microsomal and cytosolic GSTs, the latter of which are separated into six different classes (Delta, Epsilon, Omega, Sigma, Theta, and Zeta). The cytosolic gene family is comprised of 28 – 29 genes in mosquito species, 37 in *Drosophila*, and 26 in *Musca* [39, 153-155], while 36 genes encode 40 cytosolic GSTs and 5 genes encode 7 microsomal GSTs in the current *Stomoxys* genome assembly (Fig. S19; Supplementary Dataset 9). Cytosolic genes encoding members of the Delta (16 genes) and Epsilon (9 genes) classes comprise 69% of the *Stomoxys* GST family, which is comparable to the distribution in *Drosophila* and *Musca*. Delta and Epsilon GSTs are unique to insects and have been implicated in insecticide resistance [156]. The *Stomoxys* Delta genes are clustered along a single scaffold; 15 of the genes in the cluster, all with no introns, are arranged uninterrupted, which is consistent with D1 – D10 genes in *Drosophila* [143]. The five *Musca* delta genes also have no introns and are arranged uninterrupted on a single scaffold. The last *Stomoxys* Delta gene on this scaffold is alternatively spliced and shares sequence similarity and gene structure with *Drosophila* D11. In *Drosophila*, this gene is part of the uninterrupted cluster with D1 – D10, while the *Stomoxys* orthologue is separated from its cluster by a single, uncharacterized transcript.

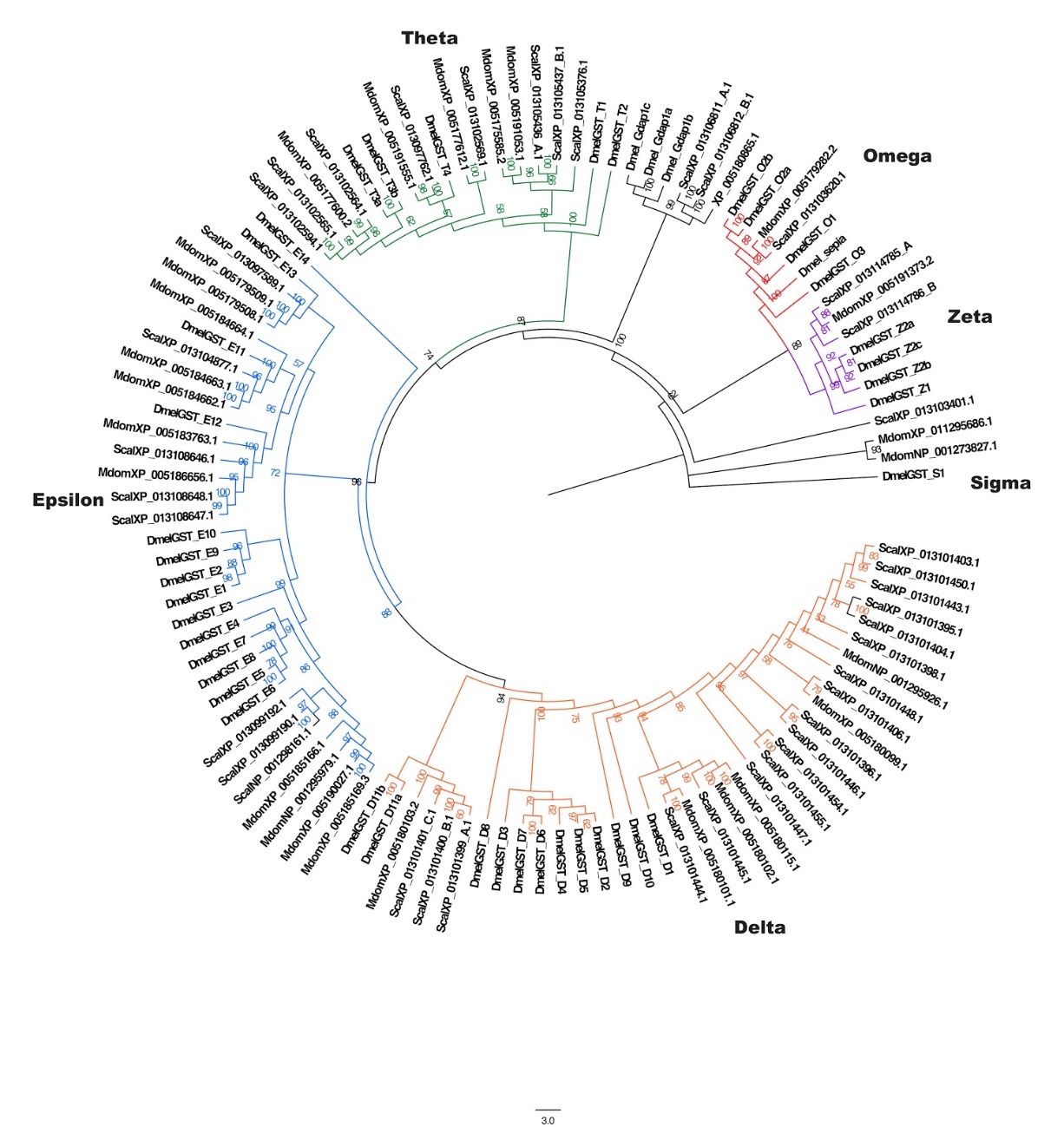

**Figure S19.** **Phylogeny of glutathione-S-transferases.** Dmel: *Drosophila melanogaster*; Mdom: *Musca domestica*.

*Cys-loop gated ion channels***.** Members of the Cys-loop gated ion channel (CysLGIC) superfamily are found throughout the insect nervous system mediating synaptic transmissions. As such, they are targets for several classes of insecticides [157, 158]. These channels are typically comprised of five subunits, each subunit of which encodes four transmembrane segments, an extracellular N-terminus that mediates ligand binding (loops A – F), and two cysteines that form a disulfide bond (‘Cys-loop’). In the *Stomoxys* genome, 26 genes were identified as putative CysLGIC subunits with orthologues to anionic and cationic channel subunits described from *Drosophila* (Fig. S20; Supplementary Dataset 10). The nicotinic acetylcholine receptors (nAChRs), activated by acetylcholine, are targets for neonicotinoid (imidacloprid) and spinosyn insecticides, and are comprised of either 5 α subunits or three α and two β subunits. Alpha subunits are characterized by the presence of neighboring cysteine residues within one of the N-terminal loops (Loop C; YxCC motif), and these cysteines are required for acetylcholine binding [159]. As the β subunits do not encode these residues, it is thought they do not participate in ligand binding. In the *Stomoxys* genome, 12 nAChR genes were identified, nine of which encode the YxCC motif and were thus characterized as α subunits (α1, α2, α3, α4, α5, α6, and two α7, one of which was a partial sequence); those without were categorized as β1 and β2. While orthologs of *Drosophila* β2 from sequenced genomes of non-drosophilid insect species are categorized as a subunits [160-162], the β2 orthologues from *Stomoxys*, *Musca* (Scott et al., 2014), and Drosophila species [163] are of the β -type. Two divergent subunits (low sequence similarity to others) were also identified in *Stomoxys*, one each of an α- and β type. The majority of the loci were located on separate scaffolds, with the exception of α1/ α2 and the two divergent subunits that are arranged in tandem on separate scaffolds. The *Stomoxys* genome also encodes GABA-gated ion channel orthologues of Rdl, Lcch3, and GRD, and one glutamate-gated ion channel, which are known targets of cyclodiene, phenylpyrazole (fipronil), and avermectin insecticides. Further, two histamine-gated and two pH-sensitive chloride channels are present in the genome, as well as orthologues of as yet uncharacterized *Drosophila* CG8916, CG12344, NtR, and the ‘Insect Group 1’ class of CysLGICs [164].

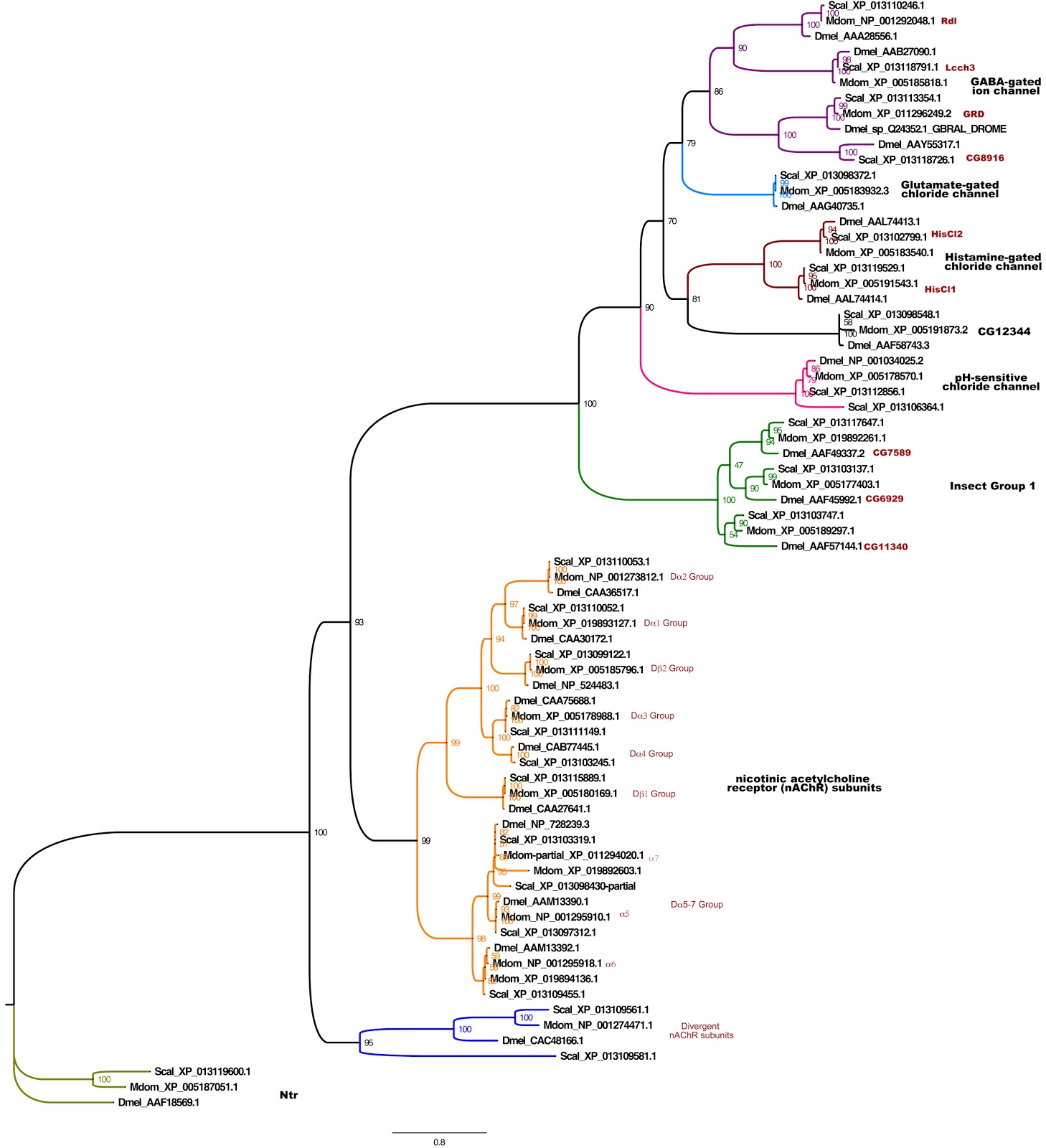

**Figure S20.** **Phylogeny of Cys-Loop Ligand Gated Ion Channels.** Scal: *Stomoxys calcitrans,*  Dmel: *Drosophila melanogaster*; Mdom: *Musca domestica*.

**Bacterial Communities Harbored by Stable Flies from Texas Dairies**

**Contributors**: Pia Untalan Olafson (USDA-ARS) and Sonja Swiger (Texas AgriLife)

**Methods.** Adult stable flies were collected at each of four Texas dairies in April and June 2015 (Lingleville and Comanche, Texas). The dairies differed in confinement method: Twenty flies per site per date were collected by aerial sweep nets in the area surrounding each dairy’s milking parlor. Sweep nets were wrapped in aluminum foil and sterilized by autoclave and were removed from the foil immediately prior to use. After each collection site, used nets were removed from the handle, placed in a plastic zip-top bag, and stored in an ice cooler until transport to the laboratory. Within 4 hours, whole flies were surface sterilized in 1% sodium hypochlorite for 15 minutes, followed by two washes in 70% ethanol and three rinses in sterile water. Individual flies were macerated in Butterfield’s phosphate buffer, and the homogenate was diluted and plated on tryptic soy broth (TSB) agar. Individual, morphologically distinct colonies were selected, suspended in Butterfield’s phosphate buffer, and the DNA isolated by rapid boiling. These DNAs were used as template in 16S PCR amplification with a universal primer pair (16SEub_61F: 5’ – GCTTAACACATGCAAG – 3’; 16SEub_1227R: 5’ – CCATTGTAGCACGTGT – 3’). Individual amplicons were sequenced in both directions, the sequences assembled, and used to query the NCBI database. Data were further processed and full-length sequences (N=170) were analyzed in mothur, v 1.38.1 [165]. Sequences were aligned using the silva.seed_v128.align file and were subsequently clustered and classified at a 97% similarity cut-off, average neighbor, silva.nr_v128.align/.tax.

**Results**. Culturable bacteria were recovered from 46% and 56% of flies screened in April and June, respectively. In total, 27 genera were identified, and the most abundant phyla represented were Proteobacteria (61%; *Aeromonas, Providencia, Pseudomonas, Serratia, Vibrio*) and Firmicutes (25%; *Bacillus, Staphylococcus, Enterococcus*) with isolates from Actinobacteria (11%; *Corynebacterium, Streptomyces*) and Bacteroidetes (2%; *Chryseobacterium*) also identified. A complete list of the individual isolates and results from querying the NCBI database is presented in Additional File 2, Table S32.

**Lateral Gene Transfer**

Contributor: Jack Werren (University of Rochester)

**Methods.** A DNA based computational pipeline was used to identify “contaminating” bacterial scaffolds and bacterial to Stomoxys lateral gene transfer (LGT) candidates in the Stomoxys genome assembly. The pipeline was originally developed by David Wheeler and John Werren [166], and has subsequently been modified. Details of the pipeline are provided in [167], and the procedure is summarized here. First, genome scaffolds are broken into 1000-bp units, which are screened against an bacterial genome database containing over 2,100 representative bacterial species. Scaffolds were then evaluated for proportion of bacterial matches along their length – those with greater than 20% bacterial matches were identified as likely bacterial scaffolds in the insect genome assembly, and were removed from further consideration. Follow-up analyses were conducted on scaffolds of greater than xx kb in length, because confidently assigning a candidate LGT requires confirmation of eukaryotic flanking sequences to the candidate. Positive bacterial hits in each 1000-bp region were compared to an “animal” database, which contains transcripts from the following eukaryotes:: *Xenopus*, *Daphnia*, *Strongylocentrotus*, *Mus*, *Homo sapiens*, *Aplysia*, *Caenorhabditis*, *Hydra*, *Monosiga*, and *Acanthamoeba*. The purpose of this step is to exclude highly conserved sequences that are shared between bacterial and eukaryotes from further analysis. We then focused our attentions on strong LGT candidates with bitscore >75 in the bacterial match and zero bitscore in the animal match. Regions were combined when adjacent 1Kb fragments each had a zero animal match and a bacterial match to the same bacterial source. These initial candidate LGTs were then manually curated, through a series of steps including (1) blastn to NCBI nr database, (2) blastx to NCBI protein database, and (3) examination of flanking regions in the scaffold to confirm presence of flanking eukaryotic (insect) orthologous sequences and gene models, (4) removal of matches due to repetitive DNA, and (5) examination of RNA sequencing data for evidence of expression. Additional steps are performed for promising LGT candidates,(e.g. expression analysis, phylogenetic reconstruction, read depth examination), but were not pursued here because of the paucity of candidates that passed manual curation in *Stomoxys calcitrans*.

**Results.** Results from the pipeline are summarized in Additional File 2: Table S33. Three candidate LGTs were detected, all of which were derived from *Wolbachia*. The *Stomoxys* strain used for the genome sequencing was not infected with *Wolbachia*; however, this is not uncommon, and detection of these segments can be due either to incomplete infection of the species, or simply due to a *Wolbachia* infection in the progenitor. One LGT occurs on scaffold NW_013171927.1 (positions 130698-130899; bitscore 242) and corresponds to a portion of the *Wolbachia* protein coding gene Dna translocase ftsk-like. A second LGT occurs on scaffold NW_013171876.1 (position 680353-680446) and is from a B group Wolbachia (bitscore 134), with similarity to the porin gene. The third is found on NW_013172024.1 (positions 765568-765650; bitscore 98.7), but it’s similarity to a specific *Wolbachia* gene is uncertain, possibly due to divergence subsequent to lateral transfer. Expression of only one of these LGTs was detected based on our current RNASeq data: the LGT region in NW_013171876.1 (680353-680446; Wolbachia porin surface protein) shows expression within the predicted 3’ UTR for a transcription factor containing a basic leucine zipper domain. This apparent 3’ UTR region was not in the *in silico* annotated gene model, but it appears to be associated based on RNAseq data examined in the WebApollo browser at VectorBase. Whether expression of the LGT is biologically significant is unknown. It should also be kept in mind that the gene annotation may be in error, and this LGT could correspond to an expressed protein coding region. Further studies of these three LGTs are warranted.

***Stomoxys* Aquaporin Proteins
Contributors:** Christopher J. Holmes and Joshua B. Benoit (University of Cincinnati)

**Summary paragraph**

Nine putative aquaporin (aqp) proteins were identified within the *Stomoxys calcitrans* genome with observed expansion in the Prip/entomoglyceroporin gene families. Unsurprisingly, *S. calcitrans* shared the greatest similarity, as they both reside in the *Muscinae* family, with *Musca domestica* but also shared some similarity with *Lucilia cuprina* aqp homologues, with seven and two respectively. Gene expansion in Drip/Prip designations was previously observed and described in *M. domestica* and *Glossina morsitans* and was suggested as a means of increased or specialized transport of water in higher-order flies [36]. Interestingly, the expansion of Prip/entomoglyceroporin genes in *S. calcitrans* shares the greatest number of homologues with *M. domestica* (4/5), and *L. cuprina* (1/5). The total number of identified aquaporin genes in *S. calcitrans* (9) is comparable to other closely related dipterans, such as *G.* *morsitans* (10), *Aedes spp.* (6), and *Drosophila* *melanogaster* (8) [168]. A recent study with *L. cuprina* (7 aqps) postulated that aqp proteins act as major contributors to osmotic pressure regulation, saliva hydration, efficient digestion, *in vivo* offspring hydration, and cold/heat tolerance [169]. Another potential function of aqps is the provision of water for milk and subsequent lactation, as was observed in *G. morsitans* [170]. It is befitting that *S. calcitrans* should display similar specialized water transport, regulation, and offspring provisions as other members of the same order. These findings prompt further investigation on the role of *S. calcitrans* aqps in regards to survivability and fecundity.

**Highlights sentence**

Nine genes including three aquaporins, three entomoglyceroporins, two Pyrocoelia rufa integral proteins (PRIP), and a big brain protein were identified as putative aquaporin-like proteins in *S. calcitrans*.

**Supplementary Materials**

The annotated *Stomoxys* aqp genes are provided as a list within Additional File 2, Table S35.

**Materials and Methods**

Putative aquaporin genes in the *S. calcitrans* genome were initially postulated via Gene Ontology term searches in Flybase (*Drosophila melanogaster*) by generating a query with proteins related to aquaporin-like functionality. The recovered nucleotide sequences were translated to peptide sequences and searched within the peptide models of *S. calcitrans*. For each possible aquaporin-like protein recovered from *S. calcitrans* models, the highest unique BLAST hit (blastp) was retrieved and confirmed via NCBI’s non-redundant arthropod database. Confirmed models were BLAST searched (blastp) against peptide sequences of *D. melanogaster, G. morsitans, L. cuprina,* and *M. domestica* for closest sequence homology. The *S. calcitrans* gene model was aligned to the closest homolog, manually annotated in WebApollo using homologues and RNA-seq data, and searched against NCBI’s non-redundant arthropod database for a final confirmation.

***Stomoxys* Cuticle Proteins
Contributors:** Andrew J. Rosendale (Mount St. Joseph University) and Joshua B. Benoit (University of Cincinnati)

**Summary**

Searching the *Stomoxys* genome with sequences typical of different cuticle protein families, as established by Willis [171], 321 genes encoding for putative cuticle proteins were identified. CutProtFam-Pred [172] was used to assign these genes to one of eight families (CPR, CPAP1, CPAP3, CPF, CPCFC, CPLCA, CPLCG, and TWDL). These results are summarized in Additional File 2, Tables S36 and S37. The total number of cuticle protein genes identified in *Stomoxys* was greater than most other insects but within the range of other dipterans [36, 172]. The number of CPR genes (196) constituted the largest group of cuticle protein genes in the *Stomoxys* genome*. Stomoxys* showed a large expansion, as compared to other insect orders, in in the TWDL (Tweedle) family, which is similar to other dipterans [172]. Generally, expansions in a cuticle protein family likely reflects adaptive evolution [173]; however, the precise role of these protein families is unknown, and the functional implications of expanded gene families needs further examination.

**Methods**

The official gene sets for *Stomoxys calcitrans* (ScalU1.1) were obtained from VectorBase and searched (BLASTp;[174]) using protein sequences that are distinctive of several families of cuticle proteins [171]. Putative cuticle proteins were then assigned to specific families of cuticle proteins using CutProtFam-Pred, a cuticle protein prediction tool [172]. Gene models coding for these putative cuticle proteins were manually annotated in WebApollo using RNA-seq data or homologous sequences. Finished models were analyzed with CutProtFam-Pred for a final confirmation.
